# Kinetic model of a determinate legume root nodule reveals plant metabolic characteristics for more efficient nitrogen fixation symbiosis

**DOI:** 10.64898/2026.04.28.721409

**Authors:** Rourou Ji, Joshua A.M. Kaste, Megan L. Matthews

**Affiliations:** Department of Civil and Environmental Engineering, Grainger College of Engineering, University of Illinois Urbana-Champaign, Urbana, IL 61802, US; Carl R. Woese Institute for Genomic Biology, University of Illinois Urbana-Champaign, Urbana, IL 61801, US

**Author notes:** Corresponding author: Megan L. Matthews.

**Keywords:** Nitrogen fixation, Kinetic modeling, Root nodule metabolism, Enzyme kinetics, Plant engineering

## Abstract

While nitrogen fertilizers are widely used in agricultural production, their application incurs significant environmental and energetic costs. In contrast, some crops are less dependent on these fertilizers because they engage in symbioses with rhizobia, nitrogen-fixing bacteria provide ammonium to the plant in exchange for carbon. However, the carbon cost associated with nitrogen fixation can negatively impact crop yields. Improving the efficiency of this metabolic process could alleviate this impact on crop productivity. Mathematical models can help us quantitatively explore metabolic behavior and identify potential targets for metabolic engineering. In this work, we developed a kinetic model of determinate root nodule metabolism, where this symbiotic exchange of carbon from the plant and nitrogen from the bacteria occurs.

We used this model to evaluate how the predicted metabolic behavior differs between inefficient and efficient nodules, and to identify potential engineering targets for improving nitrogen fixation efficiency and rate. We show that the enzymes phosphoenolpyruvate carboxylase and pyruvate kinase have significant influence on the predicted rate and efficiency of nitrogen fixation, especially when their expression is varied in combination with oxidative Pentose Phosphate Pathway enzymes like glucose-6-phosphate dehydrogenase and 6-phosphogluconolactonase. The model predicts that pairing a 3-fold decrease in glucose-6-phosphate dehydrogenase activity along with either a 3-fold increase in phosphoenolpyruvate carboxylase activity or decrease in pyruvate kinase activity could increase nitrogen fixation rate by 5.51% while improving nitrogen fixation efficiency by 7.74%.

## 1 Introduction

Nitrogen is one of the main nutrients that limit plant growth and crop yield (Du et al., 2020). To alleviate this growth limitation and maximize yields, synthetic nitrogen fertilizer is routinely applied to crops. However, runoff of these fertilizers leads to eutrophication and substantial environmental damage and biodiversity loss (Penuelas et al., 2023). Some plants, like soybean and other legumes, have evolved to form symbiotic relationships with rhizobia, a bacteria that can fix free nitrogen into ammonia. During symbiosis, these plants send carbon to the rhizobia in exchange for ammonia. This relationship allows plants to meet their nitrogen needs without relying on fertilizers. However, it also requires that the plants provide a portion of their photosynthate to the bacteria. It’s been estimated that up to 13-28% of a plant’s photosynthate could be directed to support this nitrogen fixation (Holland et al., 2023; Vance, 2008), a cost that is not incurred by plants that rely on fertilizers and take up nitrogen directly from the soil.

Previous modeling work has suggested that the carbon cost associated with nitrogen fixation can decrease soybean yields by up to 27% (Holland et al., 2023). Decreasing the amount of carbon needed to support N-fixation and improving the efficiency of this process could reduce its potential negative yield impacts, supporting both more sustainable agricultural practices and maintained crop productivity.

The exchange of metabolites between the plant and the bacteria occurs in root nodules, specialized plant structures created specifically for this symbiotic relationship. In these nodules, the carbon supplied by the plant, in the form of sucrose, is converted into dicarboxylates (malate, succinate, and fumarate) that are transported into the bacteria that have infected the nodule structure (Engelke et al., 1987). The bacteria use the energy provided by these dicarboxylates to catalyze the reduction of N_2_ to ammonia via nitrogenase (Yurgel and Kahn, 2004). Ammonium is then transported from the bacteria to the plant nodule cells where it is assimilated by the GS/GOGAT cycle into glutamate and glutamine. Glutamate and glutamine are then further converted into aspartate and asparagine in indeterminate nodules in plants of subtropical and temperate origin, or enter *de novo* purine synthesis and are converted to ureides in determinate nodules formed by legumes of tropical origin (Liu et al., 2018; Meilhoc et al., 2025).

The efficiency of nitrogen fixation, or the amount of carbon required to support this symbiotic relationship, has been shown to vary, with most observations and model estimates ranging from 2-8 g C g^-1^ N (Minchin and Witty, 2005). More efficient nitrogen fixation could come either from more efficient bacteria that need less carbon to support nitrogenase activity or from coordinating nodule metabolic pathways to reduce inefficient use of carbon. On the plant side, understanding the metabolic landscape of efficient versus inefficient nodules could provide potential plant engineering targets to improve the efficiency of this process, leaving more carbon for plant growth and yield.

Models can be useful tools for exploring metabolic inefficiencies and understanding emergent metabolic pathway behavior. While constraint-based flux balance analysis (FBA) models of nodule metabolism have been developed (diCenzo et al., 2020; Holland et al., 2023), these models are limited in their biotechnological utility because they are unable to evaluate how perturbations to individual enzymes will affect the rate and efficiency of nitrogen fixation. Kinetic metabolic models describe the change in metabolite concentrations as a system of differential equations composed of reaction fluxes that are functions of the kinetic parameters and enzyme and substrate concentrations. As such, these models can be useful for understanding the nonlinear behavior of a metabolic system and identifying metabolic engineering targets.

Kinetic models have been developed for a range of pathways and organisms, however, due to their complexity, most existing models have focused on small metabolic systems, particularly those related to central carbon metabolism or bacterial networks (Chassagnole et al., 2002; El-Mansi et al., 1994; Hernández-Esquivel et al., 2025; Jahan et al., 2016; Joshi and Palsson, 1989; Khodayari et al., 2014; Millard et al., 2017; Peskov et al., 2012; Pettersson and Ryde-Pettersson, 1988; Schoeberl et al., 2002; Wiechert, 2002). In plants, kinetic models have been developed to describe processes and identify engineering strategies in systems including photosynthesis (He et al., 2024; Zhao et al., 2021; Zhu et al., 2013), lignin biosynthesis (Sulis et al., 2023; Wang et al., 2018, 2014) and the benzenoid pathway (Colón et al., 2010). These mechanistic kinetic models have been used to reveal the influence individual enzymes have on overall metabolic pathway behavior, while also capturing system characteristics that are typically beyond the scope of constraint-based frameworks.

Here we present a kinetic model of determinate nodule metabolism integrating reactions from glycolysis, the tricarboxylic acid (TCA) cycle, the pentose phosphate pathway (PPP), glycine and serine metabolism, the GS/GOGAT cycle, alanine and aspartate metabolism, *de novo* purine biosynthesis, and allantoin biosynthesis. We used a systematic parameter sampling approach to explore the space of enzyme abundances that produce biologically reasonable predictions of the efficiency and rate of N-fixation. From this analysis, we evaluated the different metabolic characteristics of efficient versus inefficient nodules and identified the enzymes that were predicted to have the largest influence on the modeled N-fixation efficiency and rate. Finally, we simulated pairwise combinations of enzyme knockdowns or overexpression and identified potential engineering targets for improving N-fixation efficiency and rate.

## 2 Methods

### 2.1 Kinetic model construction

The determinate nodule kinetic model integrates reactions from ten major metabolic pathways: glycolysis, the tricarboxylic acid (TCA) cycle, the pentose phosphate pathway (PPP), glycine and serine metabolism, the GS/GOGAT cycle, alanine and aspartate metabolism, *de novo* purine biosynthesis, and allantoin biosynthesis. This model was constructed in three major steps (**Figure S1**): (1) Assembling known reactions in determinate nodules from all available databases, (2) generating and parameterizing the rate equations with literature-sourced datasets, and (3) building the kinetic model as a set of ordinary differential equations based on mass-balance principles.

#### 2.1.1 Assembling known reactions

Information on metabolic pathways, including reaction directionality, was sourced from the KEGG (Kanehisa and Goto, 2000) and BRENDA (Chang et al., 2021) databases, and further refined using biochemical principles reported in literature (**Text S1**). Each reaction was to ensure correct stoichiometric coefficients and atom balance. Empirical data for transmembrane transport kinetics and subcellular metabolite levels are limited, making precise compartmentalization and a full description of metabolite exchanges between subcellular compartments as well as infected and uninfected root nodule cells unrealistic. To address these gaps, reactions and metabolites were assumed to occur in a single, well-mixed compartment of a representative plant cell. While a more detailed subcellular compartmentalization could improve physiological fidelity, it would also increase model complexity and the number of unknown parameters. Given the limited experimental evidence to support such complexity, a simplified structure was adopted to maintain tractability while representing the core metabolic features of the *in vivo* system.

#### 2.1.2 Generating and parameterizing rate equations

Metabolic fluxes were modeled using Michaelis-Menten kinetics. Michaelis-Menten kinetics are simplified mass action kinetics that link substrate utilization to product formation assuming the reaction is operating under the quasi steady-state and conservation of enzyme and enzyme-complexes assumptions. Depending on the reaction, the Michaelis-Menten equations can take different forms. An example of a Michaelis-Menten equation describing a reaction featuring competitive inhibition is described in **Equation 2**.

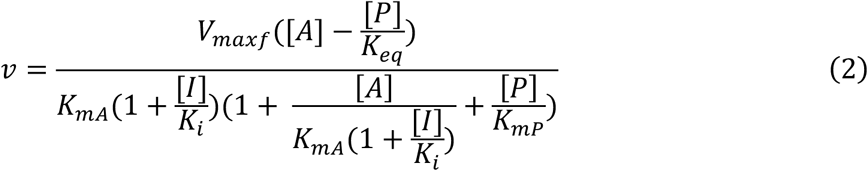

Where 𝑣 is the flux through a given reaction, *V_max_* is the maximum reaction rate when the enzyme is saturated with substrate, *K_mA_* is the substrate concentration at which the reaction rate is half of *V_max_*. A and P are respectively the substrate and product concentrations, *K_i_* is an inhibition constant, and *K_eq_* is the equilibrium constant of the reaction. Depending on the reversibility of a given reaction, the number of substrates and products involved in the reaction, and the form of inhibition known to act on it, different forms of **Equation 2** are used. Although data describing kinetics with respect to cofactors and coenzymes (e.g. ATP and NADPH) is available for some reactions in the network (Igamberdiev et al., 2014; Jin et al., 2020), most enzymes’ reaction kinetics were taken from experiments in which cofactors and coenzymes were at saturating concentration (Ashihara and Stupavska, 1984; Copeland and Morell, 1985). To ensure consistency, cofactors and coenzymes are assumed to be at saturating concentrations in all cases.

Malate, succinate, and fumarate serve as exchange metabolites between the plant cell and the bacteroid, whose transport across the peribacteroid membrane (PBM) is assumed to occur through a saturable, porin-facilitated diffusion process (Mendes et al., 2015; Millard et al., 2017). The transport through the PBM is assumed to be irreversible. The transport rate of glycine, which was predicted in a previous FBA model to be transported in small quantities across the PBM, is scaled relative to the export flux of malate, using a ratio of 0.000546 as indicated by the FBA model (Holland et al., 2023).

The rate equation for each reaction in the model is included in **Text S1**. The kinetic parameters for these rate equations were compiled from both the literature and curated biochemical databases, including BRENDA and SABIO-RK (Wittig et al., 2018). All of the kinetic parameter values used in these equations and their sources are compiled in **Table S1**.

#### 2.1.3 Building the kinetic model as a system of ordinary differential equations

The kinetic model was formulated as a system of ordinary differential equations (ODEs) shown in **Equation 3**:

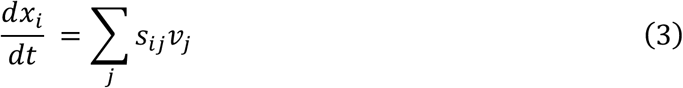

where *x_i_* denotes the concentration of metabolite *i*, s is the stoichiometric coefficient for metabolite *i* in reaction *j*, and *v_j_* represents the rate of reaction *j* which is determined by the enzyme activities. The mass balance equations for the metabolic network were constructed, resulting in a system of nonlinear differential equations provided in **Text S1**. The kinetic model comprises of 68 reactions, 60 metabolites, and 327 kinetic parameters.

Glucose is the input metabolite in our model, and we assumed that the carbohydrate intake from the root maintains glucose at a constant concentration in the nodule. Glucose concentrations in soybean nodules have been reported of 17.56 to 23.71mM (Streeter, 1987) and 28.89mM (Kouchi and Yoneyama, 1986), calculated assuming there are 20,000 cells per nodule, a cytosolic volume of 1.24×10^-10^ L per cell, and a nodule weight of 12.84 mg (Carroll et al., 1985). Based on the value in (Kouchi and Yoneyama, 1986), the glucose concentration in the model was set to 28.89mM.

### 2.2 Defining and sampling feasible and biologically reasonable parameter spaces

#### 2.2.1 Initial sampling

Due to variations in enzyme kinetics and network interactions, not all combinations of *V_max_* values allow the metabolic system to reach a steady state while preserving biological plausibility. Latin Hypercube Sampling (LHS) (Marino et al., 2008) was used to generate an initial 600,000 parameter sets across a broad range of *V_max_* values (1 < *V_max_* < 1000 mM s^-1^), sampled on a logarithmic scale to capture variation across orders of magnitude. LHS stratifies each parameter distribution into equiprobable intervals and samples once per interval, enabling efficient, space-filling exploration of the parameter space with lower variance than simple random sampling. For enzymes transketolase (TKT) and aconitate hydratase (ACN), each of them catalyzes two distinct reactions in the metabolic network, TKT1, TKT2, ACN1 and ACN2, respectively. Therefore, the common enzyme concentration ranges of TKT and ACN were determined based on the initial sampling results of their *V_max_* ranges divided by the corresponding *K_cat_* values. Then, the LHS matrix was regenerated, where values for TKT and ACN were instead sampled within TKT and ACN’s enzyme concentration ranges.

#### 2.2.2 Use of proteomic data to inform parameter space sampling

To constrain the relative enzyme levels in the model to better mirror those seen in actual legume nodules, proteomic datasets from *Glycine max* nodule mitochondria (Sin et al., 2024) and *Vicia faba* whole nodule extracts (Thal et al., 2018) were used to constrain the sampling range for some of the *V_max_* parameters. Intensity values for *Glycine max* mitochondria proteins from (Sin et al., 2024) and for *Vicia faba* whole nodule extracts from (Thal et al., 2018) were taken directly from published supplemental material or reanalyzed from publicly deposited raw data, respectively. Quantification of intensity for the *Vicia faba* dataset was performed with MaxQuant (version 2.6.7) with identical parameter settings as described in (Thal et al., 2018) and aligned to both the *Vicia faba* proteome UP001157006 and the *Rhizobium leguminosarum* proteome UP000076193, both obtained from UniProt (The UniProt Consortium, 2025). From the *Vicia faba* whole nodule dataset, only signals identified as belong to *V. faba*, as opposed to *R. leguminosarum*, were considered.

While *Vicia faba* forms indeterminate nodules (Abd-Alla et al., 2000), we hypothesized that the relative levels of TCA cycle and glycolysis enzymes, which are common to both indeterminate and determinate legume nodules, would be similar between the two nodule types (**Figure S3**). The intensity-based absolute quantification (iBAQ) (Schwanhäusser et al., 2011) and raw intensity values annotated as belonging to proteins that overlap with either the TCA cycle enzymes (Sin et al., 2024) or both the TCA cycle and glycolysis enzymes (Thal et al., 2018) were extracted, summed on a per enzyme basis, and then normalized by the total intensity of the subset of selected proteins (**Text S2**). The normalized enzyme intensity ratios were then scaled by the abundance of pyruvate dehydrogenase (PDH) in the faba bean dataset as PDH is present in both datasets. These scaled proportions were subsequently used to estimate the abundances of glycolytic enzymes for soybean, where raw enzyme intensity measurements were unavailable. Finally, the proportions of enzymes involved in both glycolysis and the TCA cycle were normalized to the abundance of PDH to enable a direct comparison between model-predicted and experimentally measured enzyme levels (**Figure S8**).

The *V_max_* sampling ranges for the other enzymes in glycolysis and the TCA cycle were estimated with their relative abundance proportions to enzyme PDH and their related *K_cat_* values (**Text S3**). Although HKI was not included in the proteomic dataset, the *V_max_* range of viable steady-state solutions appeared narrower after the initial sampling (**Figure S4**). Therefore, its range was further constrained based on those preliminary results. These focus the parameter exploration on enzyme concentration distributions that mirror those observed in the proteomic data while maintaining physiological relevance. A regression analysis of the *V_max_* values of successful steady state runs against the sampled proteomic abundances yielded a slope of 0.85 and an R^2^ of 0.76 (**Figure S8**), demonstrating that accounting for proteomic abundances influenced, but did not entirely dictate the ratios of *V_max_* values in our results.

Given the uncertainty introduced by combining data from both *G. max* and *V. faba*, as well as the uncertainty inherent in within-sample protein signal comparisons, which is addressed in part by using iBAQ values, we also ran our analysis without the proteomic weights applied but with all other steps of the pipeline kept the same. This analysis results in values for key metrics (i.e., carbon cost and rate of N-fixation) extremely similar to when the proteomic data is incorporated, with the only notable difference being that a higher proportion of sampled parameters sets result in the simulation successfully reaching steady state when using proteomic data **(Figure S5).**

From this we conclude that the incorporation of the proteomic data helps focus the parameter sampling on the viable subspace of feasible parameter values, allowing us to estimate the same values in a more computationally efficient way.

#### 2.2.3 Parameter space screening

The parameter sets resulting in steady-state solutions were further screened by a series of quantitative features to ensure that all analyses were performed on biologically/biochemically reasonable flux maps. First, we required that the export rate of malate from the plant cell to the bacteroid was greater than that of succinate, since malate is the main dicarboxylate used for nitrogen fixation (Booth et al., 2021; Mitsch et al., 2018; Udvardi and Poole, 2013). Second, we required that the N-fixation efficiency values, represented by the grams of carbon consumed per gram of nitrogen exported from the network (g C g^-1^ N) (**Equation 4**), fall in the interval [2, 8] given the available experimental data in the literature (Atkins, 1982; Layzell et al., 1988; Minchin et al., 1983; Minchin and Pate, 1973; Minchin and Witty, 2005; Ryle et al., 1979; Schubert, 1982).

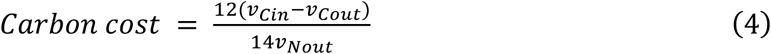

Third, we required that intracellular metabolite concentrations must remain below 200mM, considering the general physiological range of metabolite concentrations in organisms (Milo and Phillips, 2015). Since varying combinations of *V_max_* parameters lead to metabolite concentration changes of different magnitudes, this upper bound ensures that simulated concentrations remain within a physiologically acceptable range.

### 2.3. Sensitivity analysis for model validation

The efficiency with which the plant cell and bacterial cell exchange energy and nutrients was represented by the ratio of export fluxes of malate, glycine, and the import flux of NH₄^+^ in the FBA model. This value was predicted to be 1.36 in (Holland et al., 2023). However, it has been previously noted that several processes in the bacterial cell may vary in their energetic efficiency (Minchin and Witty, 2005). Based on the theoretical efficiency ranges reported in (Minchin and Witty, 2005), we estimated that the FBA-estimated ratio might realistically vary up or down by 23%. Therefore, we perturbed the ratio by this amount to characterize how sensitive the system is to changes in bacterial energy-use efficiency.

To evaluate model robustness, we performed two analyses. First, a local sensitivity analysis varied each kinetic parameter (*K_m_*, *K_i_*, *K_eq_*) by ±50% around its reference value while holding all others constant. This allowed us to assess the degree to which variations in individual parameters influence model outputs, thereby identifying parameters with disproportionate effects on system behavior. Second, to probe sensitivity to initial conditions and assess numerical and structural robustness, we used LHS to sample the initial metabolite concentrations, other than glucose which is held constant in our model, between 1 and 50 mM, and ran the model to steady-state.

### 2.5. Global sensitivity analysis for identifying influential enzyme parameters

Global sensitivity analysis was conducted through Latin Hypercube Sampling with Partial Rank Correlation Coefficient (LHS-PRCC) (Marino et al., 2008; Mckay et al., 2000) using parameter sets that passed the parameter space screening. PRCC quantifies the relationship between input parameters and model outputs by computing correlation coefficients that account for the effects of all other parameters. These coefficients provide a measure of the relative importance of each input parameter in shaping the corresponding model response. To capture the overall impact of each enzyme’s *V_max_* on its associated subnetwork, in addition to its impact on the behavior of the whole network, PRCCs over the reactions in each sub-pathway (**Figure S2**) for every enzyme were averaged and the resulting mean values and standard errors were calculated.

### 2.5. Software

The system of ODEs was solved using *ode15s* of the MATLAB package (MathWorks, ver2023a, Natick, MA, USA). The associated numerical simulations were performed through the BioCluster high performance computation facility (Carl R. Woese Institute for Genomic Biology, v3, Champaign, IL, USA). Model and analysis code to reproduce all results can be found at https://github.com/Matthews-Research-Group/DeterminateNoduleModel.git.

## 3 Results

### 3.1 A kinetic model of determinate nodule metabolism

The sampled *V_max_* parameter space resulted in 591 steady-state solutions (**Figure S4)**. These solutions have a median N-fixation efficiency of 4.72 g C g^-1^ N that ranges from 3.73 – 50.79 g C g^-1^ N (**Figure 2A**) and a median nitrogen fixation rate of 1.00 mM s^-1^ that ranges from 0.14 – 4.07 mM s^-1^ (**Figure 2B**). Limiting our steady-state solutions to those that we considered biologically plausible (n=471), the distributions remain qualitatively similar, and the median N-fixation efficiency decreases to 4.63 g C g^-1^ N (range: 3.73-7.96 g C g^-1^ N) and the rate increases slightly to 1.06 mM s^-1^ (range: 0.51-3.95 mM s^-1^) (**Figure 2C-D**). These distributions align well with previously published model and experimental determinate nodule N-fixation efficiency and rate estimates which range from 3 to 5 g C g^-1^ N and 0.04 to 0.20 mM s^-1^ (Bergersen, 1994; King et al., 1986) respectively, converted by a fresh-to-dry weight ratio of 5.76 for soybean nodules (Streeter, 1985). The predicted N-fixation rate was slightly larger, but a similar magnitude to the simulated and experimental values reported in (Bergersen, 1994; King et al., 1986). This discrepancy is likely due to differences in the nodule developmental stages (Bergersen, 1958) and to physiological assumptions incorporated into the model, including fixed nodule size, homogeneous internal conditions, and standardized environmental parameters (e.g., oxygen availability and carbon supply), which may not capture variability in growth conditions among nodules.

**Figure 2.**
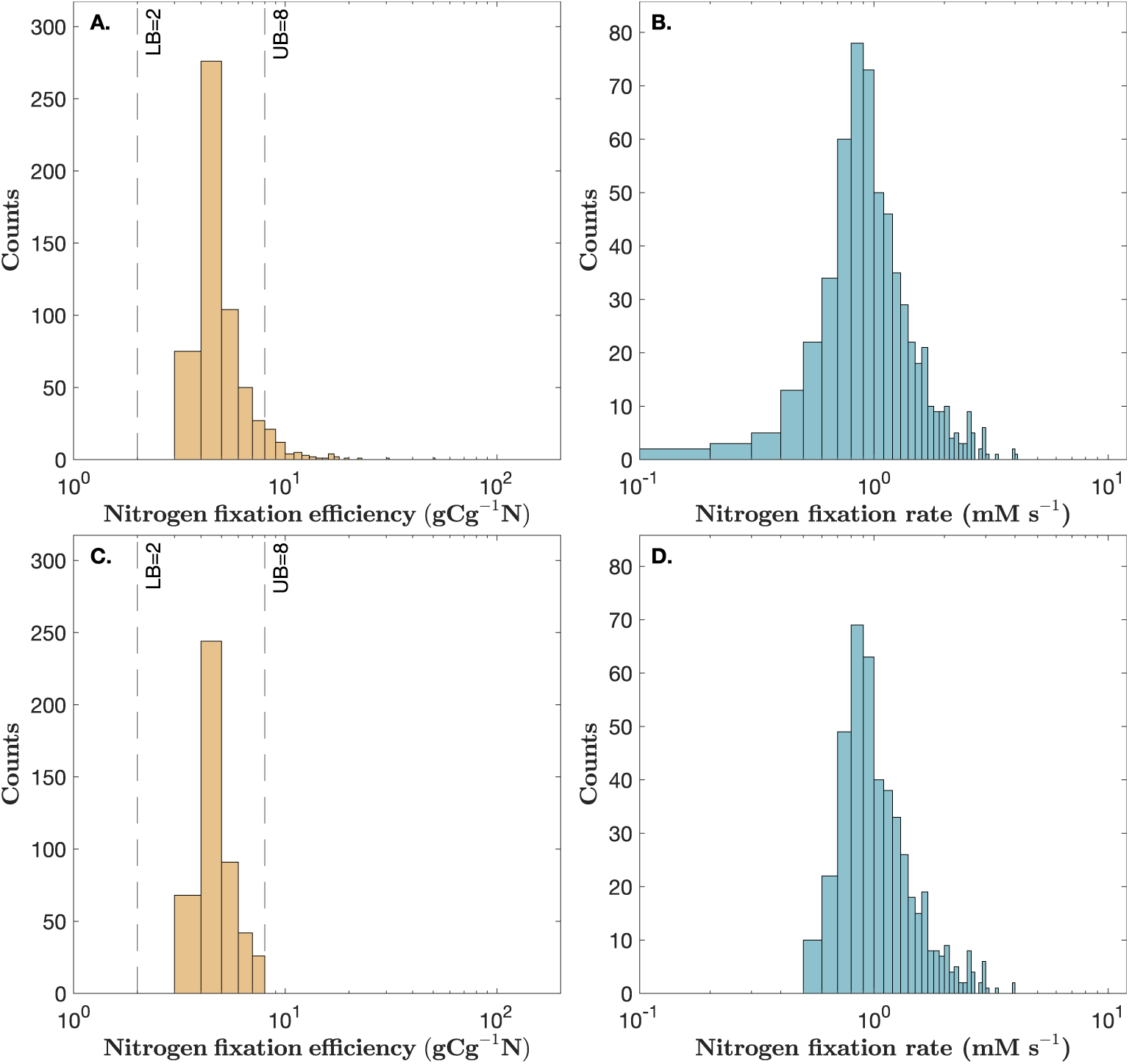
The predicted N-fixation efficiency and N-fixation rate distributions. Predicted N-fixation efficiency (yellow) and N-fixation rate (blue) distributions (A, B) before (n=591) and (C, D) after (n=471) limiting the solutions to those determined to be biologically plausible. Black dashed lines represent the lower and upper bounds of the N-fixation efficiency range considered biologically plausible.

Using the median *V_max_* of each enzyme from the set of values that result in biologically-plausible steady-state solutions, we generated a representative nodule flux map with an N-fixation efficiency of 4.17 g C g^-1^ N and an N-fixation rate of 1.14 mM s^-1^ (**Figure 3**). The main flux pathway in this representative map begins with glycolysis before entering the TCA cycle in the non-canonical direction where the dicarboxylates, primarily malate, are produced and then transported to the bacteroid and used to provide the energy needed to support nitrogenase activity. The nitrogenase output is then transported back to the nodule as NH4^+^ where it is used in the GS/GOGAT cycle and finally converted into allantoin through the *de novo* purine and allantoin synthesis pathways (**Figure 3**).

**Figure 3.**
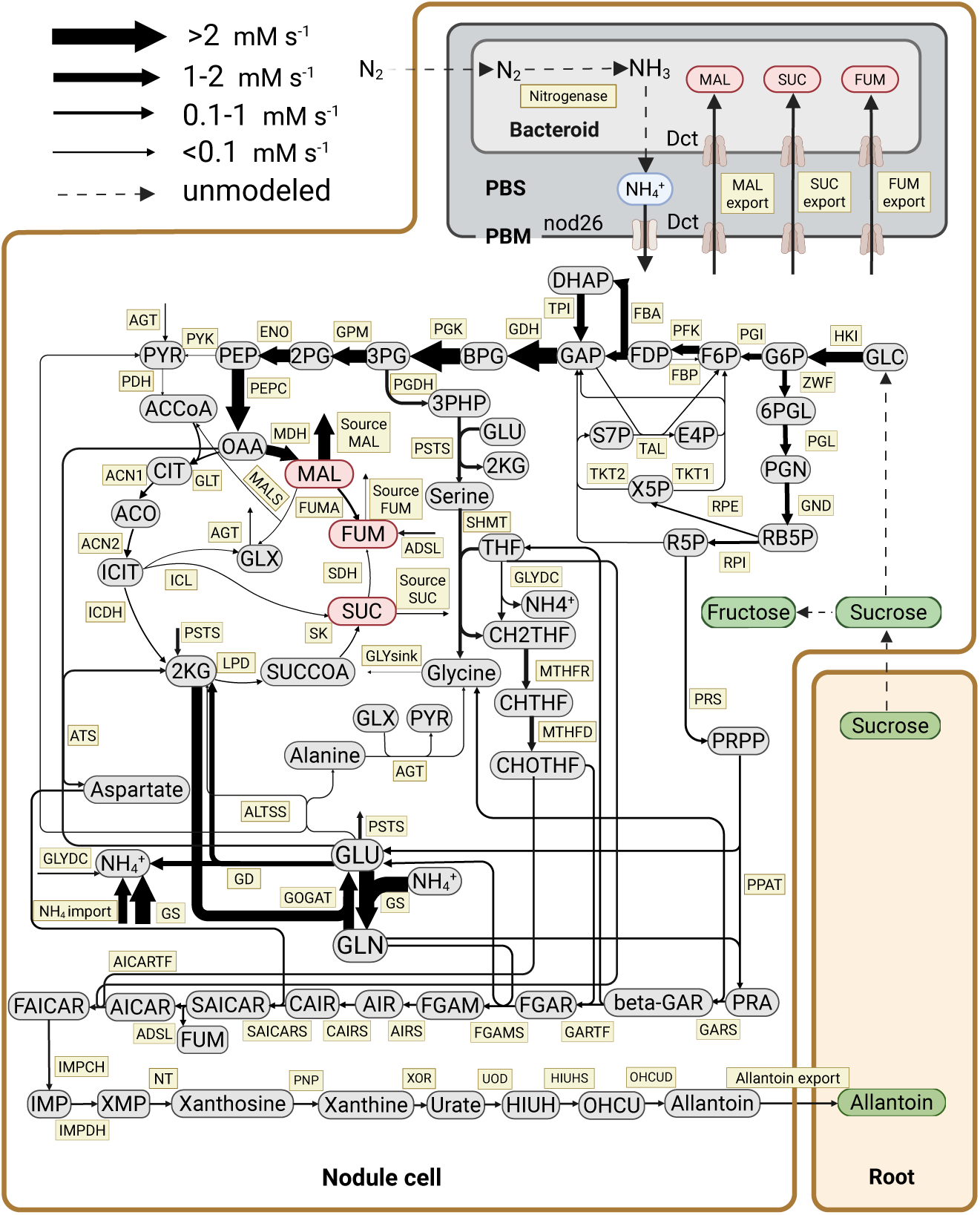
Schematic representation of flux distribution in the metabolic network of the determinate root nodule. The size of the arrows corresponds to net flux intensity. All flux values are simulated using the median *V_max_* values in unit of mM s^-1^. Arrows indicate the direction of metabolic reactions. Enzyme names are shown in yellow squares, and gray circles represent metabolites. All abbreviations used in the figure are defined in the Abbreviations section. For reversible reactions, arrows point in the direction of net flux in the flux map, and reversibility information of enzymes is shown in **Figure S2**. This flux map yields an N-fixation efficiency of 4.17 g C g^-1^ N and an N-fixation rate of 1.14 mM s^-1^.

The predicted flux of the C4-dicarboxylate exchange through the MDH shunt was observed to be substantially greater than the flux through the rest of the TCA cycle (**Figure 3**). This is consistent with previous experimental findings that malate production in nodules is primarily produced via reverse flux from oxaloacetate to malate through malate dehydrogenase (MDH) (Appels and Haaker, 1988; Day and Copeland, 1991; Vance, 2008). This results in most of the flux in the TCA cycle being non-cyclic flux from oxaloacetate to malate, with much smaller fluxes observed through the canonical cyclic TCA cycle.

Fluxes through the OPPP are necessary to generate pentose phosphate precursors to allow for *de novo* purine synthesis in combination with the ammonium received from the bacteroid. However, in the PPP, a small subset (2.97%) of parameter sets were found to result in flux maps with reversed (anabolic) flux through the PPP. Flux in this anabolic direction is associated with a slightly worse N-fixation efficiency of 4.48 ± 0.57 g C g^-1^ N, but a higher N-fixation rate of 1.83 ± 0.61 mM s-^1^, compared with the N-fixation efficiency of 4.17 g C g^-1^ N and N-fixation rate of 1.14 mM s^-1^ predicted for the representative nodule flux map (**Figure 3**).

We simulated the steady-state fluxes and intracellular metabolite concentrations for each of the screened parameter sets (**Figure S6, S7**). The simulated steady-state metabolite concentrations largely fell within the physiological range (<200 mM), with just a few scenarios that resulted in larger concentrations. The predicted steady-state concentrations of key nodule amino acids showed reasonable agreement with published measurements (**Table 1**), with the model predictions generally capturing both the correct order of magnitude and relative abundance patterns. The predicted median values of L-serine and L-glutamate overlap with experimental ranges, indicating close agreement with *in vivo* measurements. The predicted L-aspartate and L-glutamine concentrations are of similar magnitudes as their reported abundances. The predicted L-alanine concentrations are lower than empirical measurements. However, because bacteroids can excrete L-alanine into the cytosol as a potential alternative product of N-fixation (Day et al., 2001; Prell and Poole, 2006), empirical cytosolic concentrations may be overestimated if alanine export varies with physiological conditions or is enhanced under experimental settings.

**Table 1:** Experimental and model predicted amino acid concentrations from infected determinate legume nodule cells.

| Amino acid | Experimental concentration (mM) |  | Model Prediction (mM) <sup>b</sup> |  |
| --- | --- | --- | --- | --- |
|  | Cytosol <sup>a</sup><br>(Kouchi and Yoneyama, 1986) | Whole cell <sup>a</sup><br>(Vance and Heichel, 1991) | Median <sup>c</sup> | In median flux map (Figure 3) |
| L-Aspartate | 4.25 | [1.04, 7.77] | $0.87 \pm 0.82$ | 1.58 |
| L-Serine | 2.28 | [1.55, 3.62] | $2.94 \pm 2.76$ | 0.47 |
| L-Alanine | 1.71 | [1.55, 15.53] | $0.16 \pm 0.16$ | 0.07 |
| L-Glutamate | 4.45 | [1.55, 22.26] | $2.40 \pm 1.85$ | 1.01 |
| L-Glutamine | 0.16 | [1.55, 4.40] | $0.77 \pm 0.71$ | 0.30 |

<sup>a</sup> All concentrations are calculated with assumptions of 20,000 cells/nodule, $1.24 \times 10^{-10} \text{ L/cell}$ , 12.84 mg/fresh weight nodule (Carroll et al., 1985).
<sup>b</sup> Glucose concentration is taken from (Kouchi and Yoneyama, 1986) as 28.89 mM with assumptions in a. Normally, the glucose concentration in legume nodules is range from 17.56 to 23.71 mM (Streeter, 1987).
<sup>c</sup> The median concentrations are median ± MAD (median absolute deviation).

We used the representative flux map (**Figure 3**) to assess the stability of our model to changes in kinetic parameters (**Figure 4**) and initial metabolite concentrations (**Tables S2-S4**). We evaluated the sensitivity of the model predictions to the kinetic parameters using a local sensitivity analysis varying each kinetic parameter by ± 50%. The N-fixation efficiency and N-fixation rate showed some degree of sensitivity (% change ≥ 0.1%) to perturbations in 13.57% and 12.02% of the kinetic parameters respectively (**Figure 4**). The N-fixation efficiency and N-fixation rate were both most sensitive to the PEP Michaelis-Menten constant, parameter *K_mPEP_*, associated with enzymes PYK and PEPC. A 50% decrease in *K_mPEP_* for PYK caused an 5.19% increase in predicted N-fixation efficiency and a 3.41% decrease in N-fixation rate respectively. A 50% increase in *K_mPEP_* for PEPC resulted in a 2.58% increase in predicted N-fixation efficiency and a 1.72% decrease in N-fixation rate. Of the parameters identified as sensitive across both model outputs, 38.89% did not have experimentally determined kinetic values (bold parameters in **Figure 4**). The steady-state metabolite levels and reaction rates predicted using the kinetic parameters in our representative flux map were found to be numerically stable with respect to initial metabolite concentrations, further supporting model robustness (**Tables S2-S4**).

**Figure 4.**
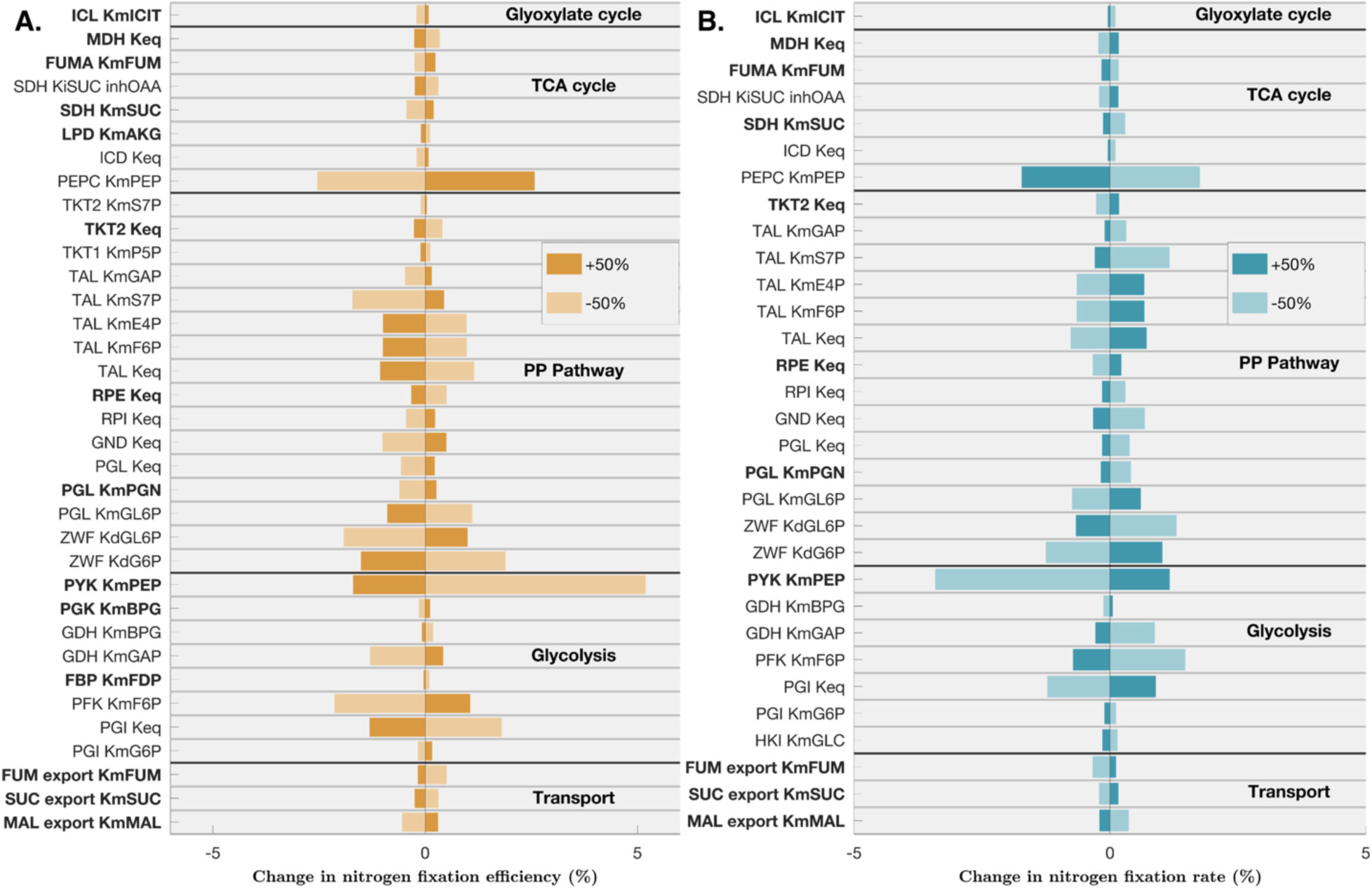
Model sensitivity to kinetic parameters. Percent change in (A) N-fixation efficiency and (B) N-fixation rate by increasing (dark bars) and decreasing (light bars) kinetic parameter values by 50%. Only kinetic parameters that resulted in a change in N-fixation efficiency or rate greater than or equal to 0.1% when perturbed by 50% are plotted. Bolded parameters lacked experimental values. *K_m_* is the Michaelis-Menten constant, *K_eq_* is the equilibrium constant, *K_d_* is the degradation constant, and *K_i_* such as *K_i_*_SUC_inhOAA_ for enzyme SDH represents the inhibition constant of substrate succinate inhibited by OAA.

### 3.2 Efficient and inefficient nodules have distinct differences in their flux maps

We refer to the top 5% (**Figure S11**) and bottom 5% (**Figure S12**) of simulations ranked by N-fixation efficiency as the high and low N-fixation efficiency subsets. Similarly, the top 5% (**Figure S13**) and bottom 5% (**Figure S14**) of simulations ranked by N-fixation rate are referred to as the high and low N-fixation rate subsets. To explore the flux patterns associated with high and low nitrogen fixation efficiency, we compared the flux maps of the high and low N-fixation efficiency (g C g^-1^ N) subsets, defined as the top and bottom 5% of simulations, respectively (**Figure S11, S12**) and examined the differences in their average fluxes (**Figure 5**) to better understand what metabolic characteristics are associated with high efficacy. The more efficient nodules exhibited considerable reduction in fluxes through the PPP and the majority of the TCA cycle reactions. As the main route into the TCA cycle, flux through the MDH shunt was substantially increased (by 0.88 mM s^-1^) in the efficient nodule (**Figure 5**). For example, in the high N-fixation efficiency nodules subset, fluxes from PEPC are directed predominantly toward MDH (1.68 mM s^-1^) and only minimally toward GLT (0.02 mM s^-1^), resulting in a better N-fixation efficiency of 3.78 g C g⁻¹ N (**Figure S11**). In contrast, the inefficient nodules in low N-fixation efficiency subset exhibit a greater allocation of PEPC flux toward GLT (0.38 mM s^-1^) and a reduced flux toward MDH (0.70 mM s^-1^), resulting in a worse N-fixation efficiency of 7.48 g C g⁻¹ N (**Figure S12**). This non-cyclic TCA flux, as shown in the flux maps where the N-fixation rate (**Figures S12**) and the efficiency (**Figure S11)** are increased compared to inefficient nodules (**Figures S12 and Figure S14)**, is also accompanied by increased fluxes through amino acid metabolic pathways, such as glycine and serine metabolism, GS/GOGAT, and *de novo* purine synthesis (**Figures S11, S13**). Additionally, flux through the PPP decreased markedly in high N-fixation efficiency nodules, with the exception of ribulose-5-phosphate isomerase (RPI), which functions at the interface between the PPP and OPPP. Although overall PPP activity declined, flux through the de novo purine synthesis pathway increased, implying that sustained or elevated RPI flux may help maintain ribose 5-phosphate (R5P) levels, which is consumed by both Transketolase 2 (TKT2) and Ribose-phosphate diphosphokinase (PRS) reactions. Therefore, an increase in RPI flux may compensate for the loss of R5P and thereby sustain purine biosynthesis despite the global reduction in overall PPP flux.

**Figure 5.**
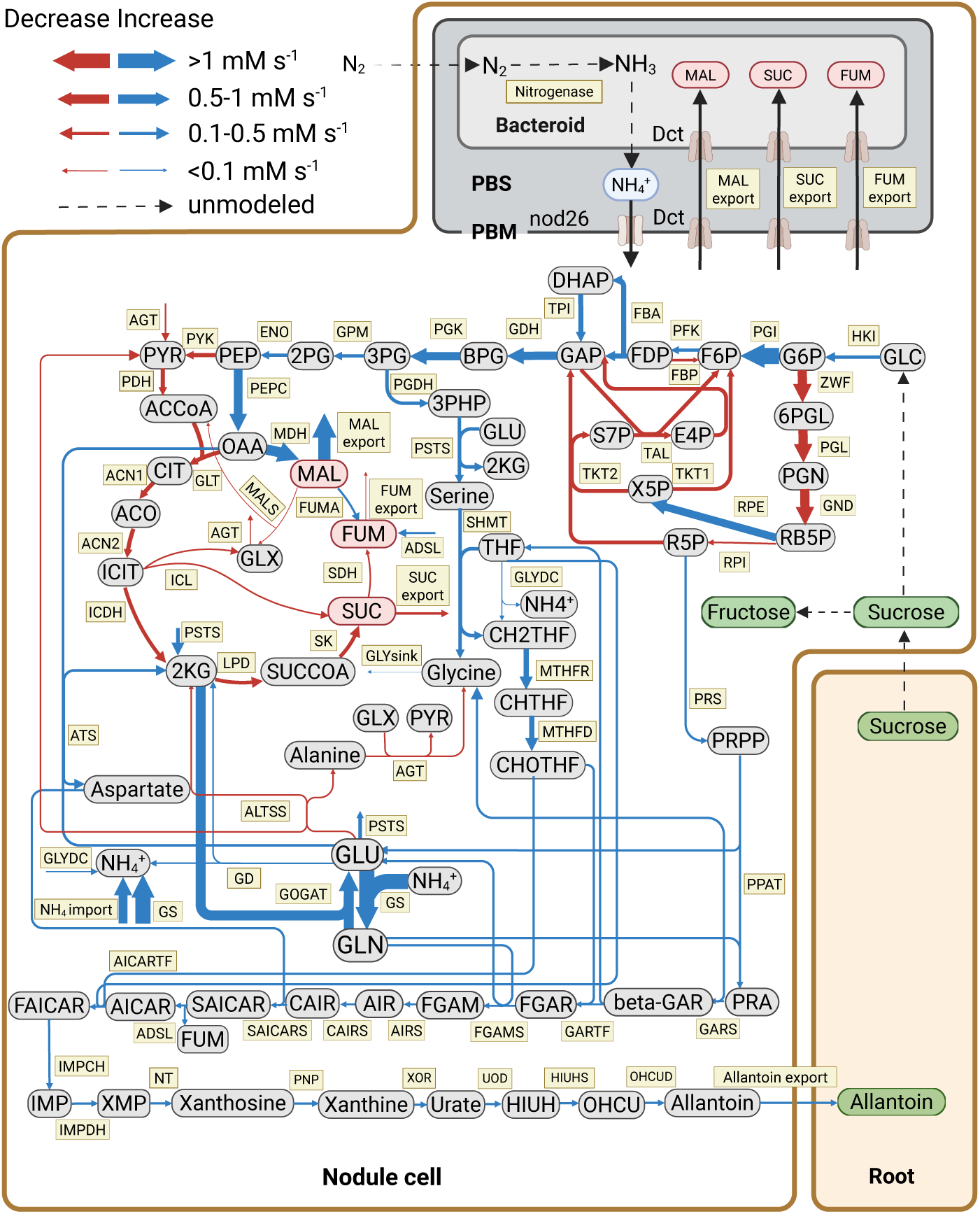
The flux change between an efficient versus inefficient determinate nodule. Flux map showing the absolute differences (efficient - inefficient) in flux between the bottom 5% of inefficient (high g C g^-1^ N) and the top 5% of most efficient (low g C g^-1^ N) nodules. The differences in flux (mM s^-1^) are presented with different arrow sizes, corresponding to their absolute difference of net flux intensity. Arrows indicate the direction of metabolic reactions, and arrow colors represent whether the efficient nodules had an increase (blue) or decrease (red) in the corresponding flux compared to the inefficient nodules.

The GS/GOGAT cycle consists of two counteracting reactions. The net flux through GS/GOGAT reflects the balance between these reactions and provides a measure of its overall contribution to nitrogen assimilation. In the high N-fixation efficiency subset, fluxes through both GS and GOGAT increased compared to their flux rates in the low N-fixation efficiency nodules, indicating coordinated enhancement of ammonium assimilation. While GS is the primary enzyme responsible for incorporating ammonium into amino acids and has been extensively characterized, GOGAT showed a parallel increase in the high N-fixation efficiency subset, consistent with its role in sustaining glutamate regeneration within the cycle. The flux through GS was lower in the low N-fixation efficiency subset compared to the high efficiency subset (**Figure 5)**, which is consistent with the reduction in GS enzyme activity reported by Anthon *et al* in the cytosol of ineffective soybean nodules (Anthon and Emerich, 1990). Despite GS and GOGAT fluxes individually showing different degrees of increased flux in the high vs. low N-fixation efficiency subset (**Figure 5**), the net GS/GOGAT flux also increased by 0.10 mM s^-1^ in nodules with better N-fixation efficiencies (**Figures S11-S12**) and by 1.11 mM s^-1^ in nodules with higher N-fixation rates (**Figures S13-S14**).

The differences between flux maps of nodules with high and low N-fixation efficiencies could be due to differences in their energy economy distributions which are not accounted for in this model as the ATP and reducing equivalent concentrations are considered to be constant. However, the presence of a non-negligible overlapping region between the net ATP balance distributions calculated from the efficient and inefficient flux maps (**Figure S19**) indicates that the differences between these flux maps are not due to differences in the energy economy. For these distributions (**Figure S19**), the net ATP balance was defined as the difference between the ATP/NADH/NADPH synthesis and consumption steady-state fluxes, with an assumed conversion efficiency of 2.5 between ATP and reducing equivalents. Additionally, this model treats the energetic efficiency of the bacteroid’s conversion of dicarboxylates into fixed nitrogen as a fixed value. We evaluated the impact of higher and lower bacteroid efficiencies on our model predictions, and corresponding changes in both N-fixation efficiency and rate were predicted, but no qualitative changes were observed in the flux distributions as a result of these changes (**Figure S9, S10**).

### 3.3 Key enzymes in central carbon metabolism have the strongest predicted influence over pathway fluxes and N-fixation efficiency and rate

We performed an LHS-PRCC global sensitivity analysis to quantify the influence of the *V_max_* parameters on the steady-state fluxes (**Figure 6**) and the N-fixation efficiency and rate (**Figure 7**). Several reactions in *de novo* purine metabolism and the allantoin synthesis pathway are primarily influenced by the *V_max_* parameters associated with enzymes outside of their subnetworks, particularly by enzymes in central carbon metabolism. The *V_max_* parameters associated with enzymes HKI, PGI, PFK, PYK, ZWF, PGL, TAL, PEPC, and LPD each have large PRCCs with different reactions across the network, indicating they are broadly influential controllers at the system level **(Figure 6)**. HKI and PGI are the entry-points for carbon and energy that are supplied by the rest of the plant in our model, as such, they shape the supply of hexose phosphates feeding both glycolysis and the PPP. PFK and PYK modulate glycolytic throughput, thereby influencing carbon flux into precursor pools. All four of these enzymes positively control the overall flux through glycolysis. ZWF, PGL, and TAL govern R5P availability and divert flux from glycolysis, directly affecting purine biosynthesis. PEPC functions as an anaplerotic entry point that channels carbon flux into malate via the TCA cycle shunt. LPD modulates the flux through the downstream portion of the TCA cycle by controlling the production of succinyl-CoA. Whereas PEPC exerts a broad, network-wide influence, LPD primarily affects fluxes within the remaining TCA reactions that generate dicarboxylates such as succinate and fumarate for export to the bacteroid. By diverting a portion of the upstream carbon flux into these downstream TCA reactions, LPD reduces the flux entering the GS/GOGAT pathway, thereby indirectly influencing nitrogen fixation and the synthesis of amino acids such as alanine and glycine.

**Figure 6.**
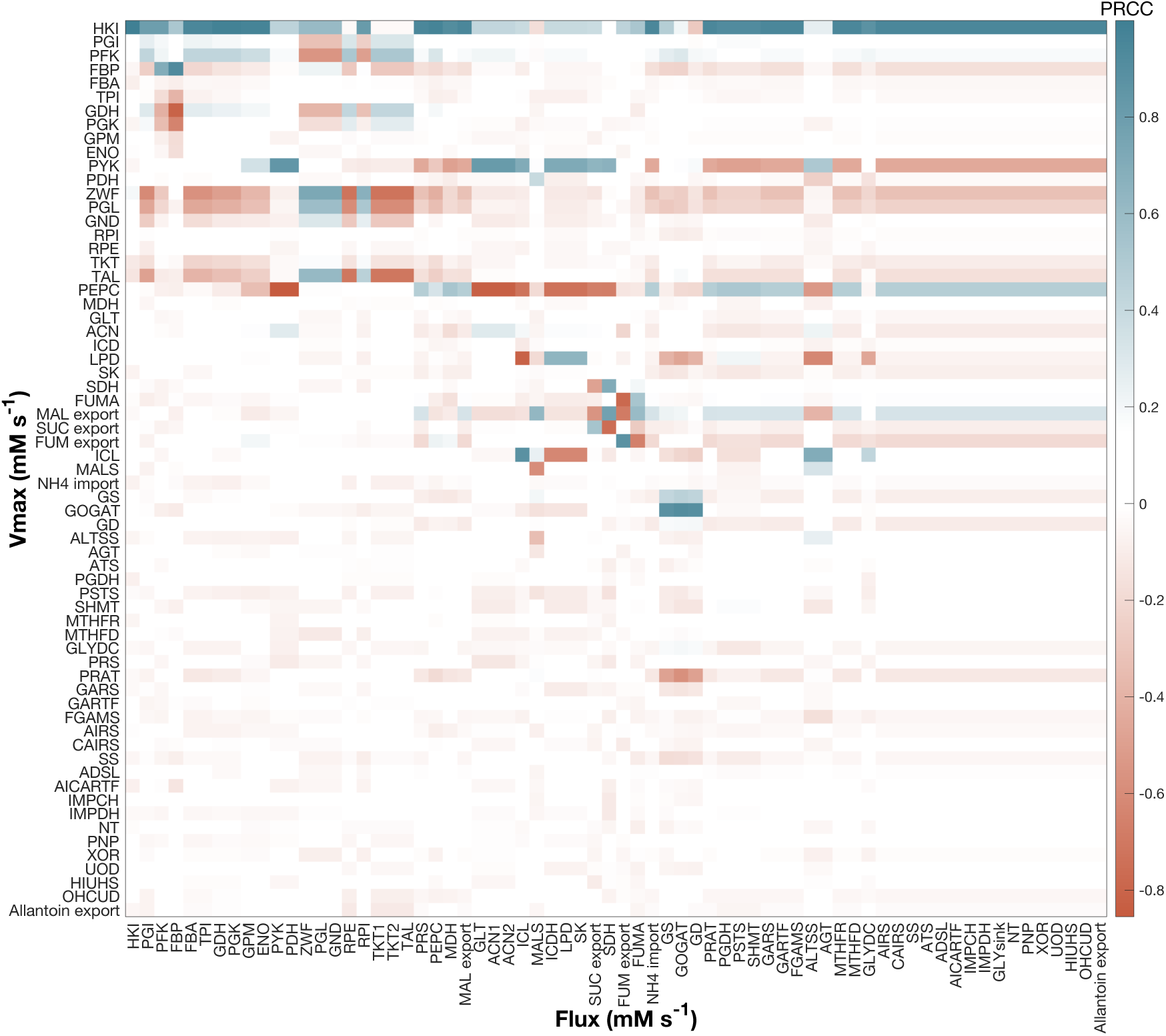
The influence of *V_max_* on nodule fluxes. Partial rank correlation coefficients between the steady-state fluxes (columns) and enzyme Vmax parameters (rows). A PRCC close to +1 or – 1 indicates a strong positive or negative influence respectively between that Vmax and steady-state flux.

**Figure 7.**
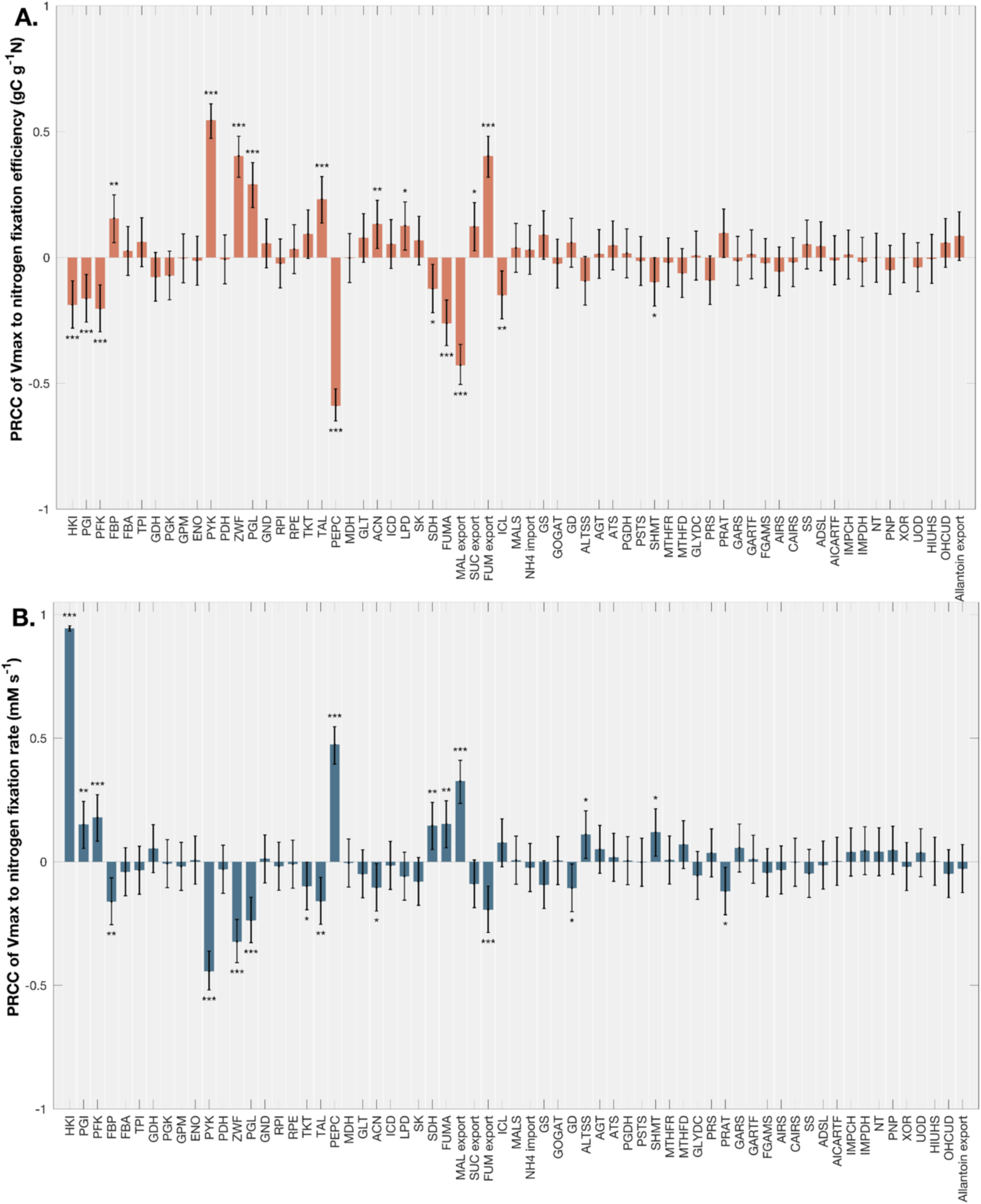
The influence of enzyme *V_max_* on N-fixation efficiency and N-fixation rate. The PRCC of the *V_max_* parameters to the (A) N-fixation efficiency (g C g^-1^ N) and (B) N-fixation rate (mM s^-1^) with the corresponding 95% confidence intervals. Asterisks denote significances: * 0.01 :s p < 0.05, ** 0.001 :s p < 0.01, *** p < 0.001.

Conversely, certain enzymes demonstrate strong control over their own subnetwork but have limited impact on reactions in other pathways, such as GS and GOGAT. Within the GS/GOGAT cycle, fluxes are more strongly driven by GOGAT (𝑝𝑟𝑐𝑐 = 0.90 ± 0.01) than by GS (𝑝𝑟𝑐𝑐 = 0.43 ± 0.02). In contrast, PRAT competes for glutamine from this cycle, diverting it into *de novo* purine biosynthesis (𝑝𝑟𝑐𝑐 = -0.52 ± 0.04). Both GOGAT and PRAT exert weak negative control over fluxes in other pathways, reflecting their specific roles in partitioning glutamine utilization, either into ureide synthesis to produce allantoin, into glutamate production to support nitrogen fixation, or into the synthesis of amino acids such as alanine and glycine.

PEPC and PYK control how carbon flows from glycolysis into the TCA cycle. These enzymes have opposite but equally strong influences on the fluxes throughout nodule metabolism and the efficiency and rate of N-fixation (**Figures 6-7**). Increased PEPC or decreased PYK result in enhanced flux through the MDH shunt of the TCA cycle (𝑝𝑟𝑐𝑐_)23_= -0.47 ± 0.02, 𝑝𝑟𝑐𝑐_)4)5_= 0.50 ± 0.03) which is a more direct route to producing malate than the canonical TCA cycle. Fluxes in *de novo* purine biosynthesis are also increased (𝑝𝑟𝑐𝑐_)23_ = -0.44, 𝑝𝑟𝑐𝑐_)4)5_ = 0.52) when PEPC and PYK are altered to enhance the flux through the MDH shunt, while the remaining TCA cycle fluxes are reduced (𝑝𝑟𝑐𝑐_)23_= 0.62 ± 0.25, 𝑝𝑟𝑐𝑐_)4)5_= -0.63 ± 0.25).

In addition to impacting nodule flux distribution, PEPC and PYK also strongly influence nodule N-fixation efficiency and nitrogen fixation rate (**Figure 7**). Overall, increasing the activities of PGI, PFK, PEPC, FUMA, and malate transport, or decreasing the activities of FBP, PYK, ZWF, PGL, TAL, SHMT and fumarate transport improve both the predicted efficiency and rate of N-fixation. Increasing ALTSS or decreasing GD and PRAT only improve the N-fixation rate and have no impact on the efficiency, while decreasing ICL or increasing LPD only improve the predicted efficiency and has no effect on the rate.

To identify potential realistic engineering strategies for improving the efficiency and rate of N-fixation, we simulated 3-fold changes in expression of pathway enzyme pairs from their median flux map expression (**Figure 3**). Among the 231 enzyme pairs analyzed, 78 resulted in a predicted improved efficiency and/or rate (**Figure 8**). The other 153 pairs exhibited no effect (**Table S5**). The largest improvements in both efficiency and rate were predicted for the strategy of down-regulating both PYK and ZWF (7.74% for efficiency, 5.51% for rate) and the strategy of over-expressing PEPC and down-regulating ZWF (7.72% for efficiency, 5.50% for rate) (**Figure 8**). Comparatively, individually down-regulating ZWF by itself resulted in only 3.98% and 1.17% improvements in efficiency and rate, and down-regulating PYK (or over-expressing PEPC) resulted in only 4.02% and 1.16% improvements in efficiency and rate respectively (**Figure S17, S18**).

**Figure 8.**
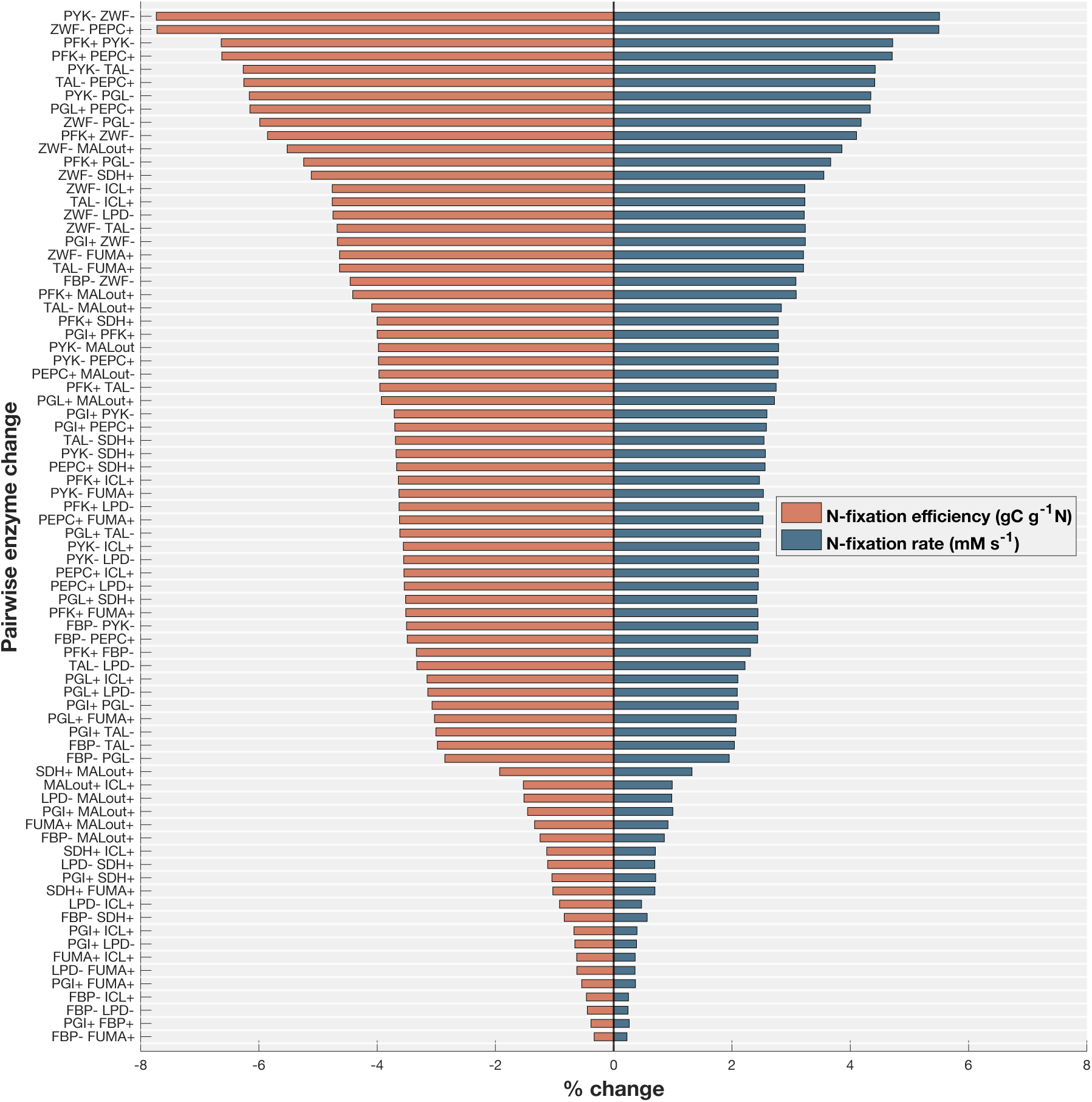
The pairwise impact of enzyme *V_max_* on N-fixation efficiency and N-fixation rate. Percent change in the N-fixation efficiency (orange) and N-fixation rate (blue) by increasing or decreasing the *V_max_* values by 3-fold. The figure includes only enzyme pairs that influence either carbon cost or N-fixation rate by more than 0.1%.

## 4 Discussion

In this study, we developed the first kinetic model of determinate nodule metabolism in legumes. The model exhibits robustness to kinetic parameter perturbations, produces intracellular metabolic concentration predictions that align with reported measurements, and reproduces biologically reasonable nitrogen fixation rates and efficiencies. This model provides a platform for analyzing pathway-level flux patterns and identifying metabolic reallocation associated with improved nitrogen fixation rate and efficiency.

### Coordinated over-/under-expression of ZWF and PEPC/PYK could enhance N-fixation

Enzymes in the model exhibit different levels of influence on N-fixation efficiency and rate (**Figure 7**). Our analysis found that the over-/under-expression of influential enzyme pairs could be used to achieve meaningful improvement in N-fixation efficiency and N-fixation rate (**Figure 8**). Specifically, coordinated adjustments of ZWF with either PEPC or PYK enabled the greatest enhancement of N-fixation (**Figure 8**). A simulated three-fold change in expression of only these enzymes was predicted to improve the N-fixation efficiency from 4.17 g C g^-1^ N to 3.84 g C g^-1^ N, achieving approximately 75% of the maximum possible improvement relative to the most efficient nodules identified in our simulations (3.73 g C g^-1^ N).

Notably, all three of these enzymes are located at branch points in the metabolic network. As the entry-point enzyme of the PPP, ZWF governs the production of pentose phosphate precursors required for *de novo* purine synthesis. PEPC and PYK determine the entry point of glycolytic intermediates into the TCA cycle, and have been shown to be regulated by cytosolic malate fluctuations in infected nodules (McCloud et al., 2001). PEPC and PYK have also been identified as key regulators of carbon metabolism in previous studies, particularly influencing carbohydrate uptake and carbon flux rerouting (Bettenbrock et al., 2006; Emmerling et al., 2002; Peskov et al., 2012). In our study, the preferential routing of flux through PEPC emerged from the kinetic structure of the model. Specifically, the relative catalytic capacities of enzymes surrounding PEPC create a kinetically favorable pathway, leading to PEPC carrying the predominant flux. This route provides a more direct entry into the reverse TCA cycle and the MDH shunt, enhancing the production of dicarboxylates transported to the bacteroid and thereby improving N-fixation efficiency. These findings suggest that metabolic engineering of nodules to alter PEPC or PYK expression could be a promising strategy to enhance N-fixation performance.

Since altering the expression of either of these enzymes will influence the flux through the other enzyme, model simulations indicate that targeting both enzymes together would mostly be redundant. Modulating either enzyme alone was predicted to increase N-fixation efficiency by 3.60% and N-fixation rate by 1.94% relative to the baseline model state. Targeting either PEPC or PYK in combination with ZWF, however, is predicted to provide the largest benefit (7.72% for efficiency, 5.50% for rate). This combination is particularly effective because these enzymes participate in carbon source partitioning toward downstream pathways, and all three exhibit substantial flux changes as the nodule becomes more efficient, consistent with the flux map differences observed between efficient and inefficient nodules (**Figure 5**).

### Nitrogen fixation is predominantly shaped by central carbon metabolism

Our analysis demonstrates that enzymes of central carbon metabolism constitute the primary control points of symbiotic N-fixation in nodules, exerting greater influence than enzymes in the *de novo* purine and allantoin synthesis pathways. Although flux maps comparing the high and low N-fixation efficiency and rate subsets reveal broad differences throughout the metabolic network (**Figure S11-S14**), our PRCC analysis indicates that only a subset of enzymes significantly impact N-fixation efficiency and rate (**Figure 7**), with most of those enzymes belonging to central carbon metabolism.

In the high N-fixation efficiency subset, central carbon metabolism shifts toward glycolysis with reduced PPP and TCA fluxes, indicating a metabolic reallocation for carbon sources that favors precursor generation over oxidative metabolism (**Figure 5**). This reallocation underscores the balance and integration among these central pathways in maintaining metabolic homeostasis. Such reorganization likely reflects adjustments in how imported sucrose is utilized within the nodule.

The role of the PPP is more complex. On one hand, it provides R5P and related precursors necessary for *de novo* purine synthesis, which underpins ureide biosynthesis in determinate nodules. On the other hand, our analysis reveals a negative relationship between PPP activity and fluxes through *de novo* purine metabolism (**Figure 6**). This discrepancy may reflect the need of the nodule to balance carbon and nitrogen fluxes. Excessive PPP activity can divert carbon away from glycolysis, reducing PEP availability for dicarboxylate production, which is critical for supplying the bacteroid with respiratory substrates. Conversely, too little flux through the PPP may fail to generate sufficient precursors for purine biosynthesis, thereby limiting ureide production and nitrogen transport. Although direct evidence remains limited for determinate nodules, our flux analysis suggests that PPP represents a pivotal control point at the nexus of C and N balance and represents a direction for future research, where flux partitioning must be precisely optimized to sustain both bacteroid energy supply and nitrogen export in assimilable forms.

### Fluxes through GS/GOGAT are highly correlated with improvements in N-fixation efficiency, but are controlled by enzymes in other areas of metabolism

Among the key reactions linking carbon and nitrogen metabolism, the GS/GOGAT cycle plays a central role in assimilating ammonium into organic forms of N. Our analysis shows that neither enzyme in this pathway exerts strong control over N-fixation rate or efficiency (**Figure 7**), despite fluxes through GS and GOGAT being highly correlated with N-fixation efficiency (**Figure 5**).

PYK, PEPC, LPD, GS, GOGAT, and PRAT exert the strongest control over the net flux through GS/GOGAT (**Figure S15**), but only PYK and PEPC showed strong controls over N-fixation efficiency or rate (**Figure 7**). Therefore, the observed increase in net flux through GS/GOGAT is an emergent property of the enhanced N-fixation efficiency and rate resulting from the modulation of PYK and PEPC. Together, these findings highlight the significant role that GS/GOGAT plays in N-fixation and associated internal flux adjustments under varying nodule efficiencies. However, they also indicate that the increased net flux through GS/GOGAT is primarily a secondary effect of broader metabolic changes, rather than a direct determinant of N-fixation efficiency or rate. This cross-pathway regulation highlights GS/GOGAT as a key metabolic junction, suggesting that engineering the surrounding network to modulate flux through GS/GOGAT may provide a promising strategy to enhance symbiotic N-fixation. The underlying mechanisms regulating flux through the GS/GOGAT cycle deserves further in-depth investigation.

## 5 Conclusion

We developed a detailed *in silico* model to investigate metabolism in determinate root nodules. This model provides critical insights into the metabolism underlying nitrogen fixation and highlights key regulatory nodes. Our pairwise perturbation results further reveal how nitrogen fixation performance might respond to targeted engineering strategies. The methodology developed in this study can be readily adapted to investigate other organisms with complex metabolic systems. A difficulty in developing kinetic models is the amount of biological information that is often required and prohibitive to measure in most plant systems. However, the parameter sampling approach used in this study demonstrates how these models can still provide informative insights into metabolic pathway dynamics.

## Abbreviations Metabolite

GLC: Glucose
G6P: glucose-6-phosphate
F6P: fructofuranose 6-phosphate
FDP: fructofuranose 1,6-bisphosphate
GAP: Glyceraldehyde 3-phosphate
DHAP: Dihydroxyacetone phosphate
BPG: 3-Phospho-D-glyceroyl phosphate
3PG: 3-Phospho-D-glycerate
2PG: 2-Phospho-D-glycerate
PEP: Phosphoenolpyruvate
PYR: Pyruvate
6PGL: Glucono-1,5-lactone 6-phosphate
PGN: 6-Phospho-D-gluconate
RB5P: Ribulose 5-phosphate
R5P: Ribose 5-phosphate
X5P: Xylulose 5-phosphate
E4P: Erythrose 4-phosphate
S7P: Sedoheptulose 7-phosphate
ACCOA: Acetyl-CoA
OAA: Oxaloacetate
CIT: Citrate
ACO: cis-Aconitate
ICIT: Isocitrate
2KG: 2-Oxoglutarate
SUCCOA: Succinyl-CoA
SUC: Succinate
FUM: Fumarate
MAL: Malate
GLX: Glyoxylate
3PHP: 3-P-hydroxypyruvate
CH2THF: 5,10-methylenetetrahydrofolate
THF: Tetrahydrofolate
CHTHF: 5,10-methenyltetrahydrofolate
CHOTHF: 10-formyltetrahydrofolate
PRPP: 5-phospho-alpha-D-ribose 1-diphosphate
PRA: 5-phospho-beta-D-ribosylamine
GAR: N1-(5-phospho-D-ribosyl)glycinamide
FGAR: N2-formyl-N1-(5-phospho-D-ribosyl)glycinamide
FGAM: 2-(formamido)-N1-(5-phospho-D-ribosyl)acetamidine
AIR: 5-amino-1-(5-phospho-D-ribosyl)imidazole
CAIR: 5-amino-1-(5-phospho-D-ribosyl)imidazole-4-carboxylate
SAICAR: (S)-2-[5-amino-1-(5-phospho-D-ribosyl)imidazole-4-carboxamido] succinate
AICAR: 5-amino-1-(5-phospho-D-ribosyl)imidazole-4-carboxamide
FAICAR: 5-formamido-1-(5-phospho-D-ribosyl)imidazole-4-carboxamide
IMP: Inosine monophosphate
XMP: Xanthosine 5’-phosphate
HIUH: 5-hydroxyisourate
OHCU: 5-hydroxy-2-oxo-4-ureido-2,5-dihydro-1H-imidazole-5-carboxylate

## Enzyme

Glycolysis: HKI: Hexokinase, PGI: Glucose-6-phosphate isomerase, PFK: phosphofructokinase, FBP: Fructose-1,6-bisphosphatase, FBA: Aldolase, TPI: Triosephosphate isomerase, GDH: lyceraldehyde-3-phosphate dehydrogenase, PGK: phosphoglycerate kinase, GPM: phosphoglycerate mutase, ENO: Enolase, PEPC: phosphoenolpyruvate carboxylase, PYK: Pyruvate kinase, PDH: Pyruvate dehydrogenase; ZWF: glucose-6-phosphate dehydrogenase, PGL: 6-phosphogluconolactonase, GND: 6-Phosphogluconate dehydrogenase, RPE: Ribulose-5-phosphate 3-epimerase, RPI: ribose-5-phosphate isomerase, TKT1: Transketolase 1, TKT2: Transketolase 2, TAL: Transaldolase; GLT: citrate synthase, ACN1: aconitate hydratase, ACN2: aconitate hydratase, ICDH: isocitrate dehydrogenase, LPD: Lipoamide dehydrogenase, SK: succinyl-CoA synthetase, SDH: succinate dehydrogenase, FUMA: fumarate hydratase, MDH: malate dehydrogenase, ICL: isocitrate lyase, MALS: malate synthase; GS: glutamine synthetase, GOGAT: glutamate synthase, GD: glutamate dehydrogenase; ALTSS: alanine transaminase, ATS: aspartate transaminase, AGT: alanine-glyoxylate transaminase; PGDH: phosphoglycerate dehydrogenase, PSTS: phosphoserine transaminase, SHMT: glycine hydroxymethyltransferase, GLYDC: Glycine decarboxylase, MTHFR: methylenetetrahydrofolate dehydrogenase, MTHFD: methenyltetrahydrofolate cyclohydrolase; PRS: ribose-phosphate diphosphokinase, PRAT: amidophosphoribosyltransferase, GARS: phosphoribosylamine-glycine ligase, GARTF: Glycinamide ribonucleotide transformylase, FGAMS: phosphoribosylformylglycinamidine synthase, AIRS: phosphoribosylformylglycinamidine cyclo-ligase, CAIRS: phosphoribosylaminoimidazole carboxylase, SAICARS: phosphoribosylaminoimidazolesuccinocarboxamide synthase, ADSL: adenylosuccinate lyase, AICARTF: phosphoribosylaminoimidazolecarboxamide formyltransferase, IMPCH: IMP cyclohydrolase, IMPDH: IMP pyrophosphorylase, NT: 5’-nucleotidase, PNP: purine-nucleoside phosphorylase, XOR: xanthine dehydrogenase, UOD: uricase, HIUHS: hydroxyisourate hydrolase, OHCUD: 2-oxo-4-hydroxy-4-carboxy-5-ureidoimidazoline decarboxylase; Peribacteroid space; PBM: peribacteroid membrane; DCT: Dicarboxylate Transport.

## Data availability

kinetic model was developed with the software *MATLAB_R2023a*. All data, models, and code to reproduce the work in this study can be found at https://github.com/Matthews-Research-Group/DeterminateNoduleModel.git and in the online supplemental material accompanying the manuscript.

## Acknowledgments

This work was supported by the project Enabling Nutrient Symbioses in Agriculture (ENSA), which is funded by Gates Agricultural Innovations (INV-57461) (RJ, JAMK, and MLM). Figures 1, 3, and 5 were created with BioRender.com.

## Author contributions

RJ and MLM conceived the study. RJ developed the kinetic models, performed the simulations, and analyzed the resulting data. JAMK processed the proteomic data. RJ, JAMK, and MLM interpreted the results. RJ wrote the manuscript. All authors contributed to revisions and approved the final version.

