## Supplemental Text S1 for "Kinetic model of a determinate legume root nodule reveals plant metabolic characteristics for more efficient nitrogen fixation symbiosis"

**2 ODE systems** **9**

**3 Reactions and kinetic parameters** **11**

**4 Supplemental figures and tables** **16**

### 1 Rate law equations

#### 1.1 Glycolysis

$$v_{HKI} = \frac{V_{\max}\left(GLC - \frac{G6P}{K_{\mathrm{eq}}}\right)}{K_{mGLC}\left(1 + \frac{GLC}{K_{mGLC}} + \frac{G6P}{K_{mG6P}}\right)} \tag{S1}$$

$$v_{PGI} = \frac{V_{\max}\left(G6P - \frac{F6P}{K_{\mathrm{eq}}}\right)}{K_{mG6P}\left(1 + \frac{G6P}{K_{mG6P}} + \frac{F6P}{K_{mF6P}}\right)} \tag{S2}$$

$$v_{PFK} = \frac{V_{\max} \cdot F6P}{K_{mF6P} + F6P} \tag{S3}$$

$$v_{FBP} = \frac{V_{\max} \cdot FDP}{K_{mFDP} + FDP} \tag{S4}$$

$$v_{FBA} = \frac{V_{\max}\left(FDP - \frac{GAP \cdot DAP}{K_{\mathrm{eq}}}\right)}{K_{mFDP}\left(1 + \frac{FDP}{K_{mFDP}} + \frac{GAP}{K_{mGAP}} + \frac{DAP}{K_{mDAP}} + \frac{GAP \cdot DAP}{K_{mGAP}K_{mDAP}}\right)} \tag{S5}$$

$$v_{TPI} = \frac{V_{\max} \cdot DAP\left(DAP - \frac{GAP}{K_{\mathrm{eq}}}\right)}{K_{mDAP}\left(1 + \frac{GAP}{K_{mGAP}} + \frac{DAP}{K_{mDAP}} + \frac{3PG}{K_{i3PG}} + \frac{PEP}{K_{iPEP}}\right)} \tag{S6}$$

$$v_{GDH} = \frac{V_{\max}\left(GAP - \frac{BPG}{K_{\mathrm{eq}}}\right)}{K_{mGAP}\left(1 + \frac{GAP}{K_{mGAP}} + \frac{BPG}{K_{mBPG}}\right)} \tag{S7}$$

$$v_{PGK} = \frac{V_{\max}\left(BPG - \frac{3PG}{K_{\mathrm{eq}}}\right)}{K_{mBPG}\left(1 + \frac{BPG}{K_{mBPG}} + \frac{3PG}{K_{m3PG}}\right)} \tag{S8}$$

$$v_{GPM} = \frac{V_{\max}\left(3PG - \frac{2PG}{K_{\mathrm{eq}}}\right)}{K_{m3PG}\left(1 + \frac{3PG}{K_{m3PG}} + \frac{2PG}{K_{m2PG}}\right)} \tag{S9}$$

$$v_{ENO} = \frac{V_{\max}\left(2PG - \frac{PEP}{K_{\mathrm{eq}}}\right)}{K_{m2PG}\left(1 + \frac{2PG}{K_{m2PG}} + \frac{PEP}{K_{mPEP}}\right)} \tag{S10}$$

$$v_{PYK} = \frac{V_{\max} \cdot PEP}{K_{mPEP} + PEP} \tag{S11}$$

$$v_{PDH} = \frac{V_{\max} \cdot PYR}{K_{mPYR}\left(1 + \frac{PYR}{K_{iPYR}}\right) + PYR} \tag{S12}$$

#### 1.2 PP pathway

$$v_{ZWF} = \frac{V_{\max}\left(G6P - \frac{GL6P}{K_{\mathrm{eq}}}\right)}{K_{dG6P}\left(1 + \frac{G6P}{K_{dG6P}} + \frac{GL6P}{K_{dGL6P}}\right)} \tag{S13}$$

$$v_{PGL} = \frac{V_{\max}\left(GL6P - \frac{PGN}{K_{\mathrm{eq}}}\right)}{K_{mGL6P}\left(1 + \frac{GL6P}{K_{mGL6P}} + \frac{PGN}{K_{mPGN}} + \frac{G6P}{K_{iG6P}}\right)} \tag{S14}$$

$$v_{GND} = \frac{V_{\max}\left(PGN - \frac{RB5P}{K_{\mathrm{eq}}}\right)}{K_{mPGN}\left(1 + \frac{PGN}{K_{mPGN}} + \frac{RB5P}{K_{dRB5P}}\right)} \tag{S15}$$

$$v_{RPE} = \frac{V_{\max}\left(X5P - \frac{RB5P}{K_{\mathrm{eq}}}\right)}{K_{mX5P}\left(1 + \frac{X5P}{K_{mX5P}} + \frac{RB5P}{K_{mRB5P}}\right)} \tag{S16}$$

$$Den_{RPI} = 1 + \frac{RB5P}{K_{mRB5P}\left(1 + \frac{E4P}{K_{iRB5P,\mathrm{inhE4P}}}\right)} + \frac{R5P}{K_{mR5P}\left(1 + \frac{E4P}{K_{iR5P,\mathrm{inhE4P}}} + \frac{GAP}{K_{iR5P,\mathrm{inhGAP}}} + \frac{PGA3}{K_{iR5P,\mathrm{inhPGA3}}}\right)} \tag{S17}$$

$$v_{RPI} = \frac{V_{\max}\left(RB5P - \frac{R5P}{K_{\mathrm{eq}}}\right)}{\left(1 + \frac{PGN}{K_{iR5P,\mathrm{inhPGN}}}\right) K_{mRB5P}\left(1 + \frac{E4P}{K_{iRB5P,\mathrm{inhE4P}}}\right) Den_{RPI}} \tag{S18}$$

$$\begin{aligned} Den_{TKT} = 1 &+ \left(1 + \frac{GAP}{TKT1_{K_{mGAP}}}\right)\left(\frac{F6P}{TKT1_{K_{mF6P}}} + \frac{S7P}{TKT2_{K_{mS7P}}}\right) \\ &+ \frac{GAP}{TKT2_{K_{mGAP}}} + \frac{X5P\left(1 + \frac{E4P \cdot R5P}{TKT1_{K_{mP5P}}}\right) + E4P + R5P}{TKT2_{K_{mP5P}}} \end{aligned} \tag{S19}$$

$$v_{TKT1} = \frac{V_{\max}\left(F6P \cdot GAP \cdot K_{\mathrm{eq}} - E4P \cdot X5P\right)}{TKT2_{K_{mP5P}} \cdot TKT1_{K_{mP5P}} \cdot Den_{TKT}} \tag{S20}$$

$$v_{TKT2} = \frac{V_{\max}\left(S7P \cdot GAP \cdot K_{\mathrm{eq}} - X5P \cdot R5P\right)}{TKT2_{K_{mP5P}} \cdot TKT1_{K_{mP5P}} \cdot Den_{TKT}} \tag{S21}$$

#### 1.3 TCA cycle

$$v_{GLT} = \frac{\frac{V_{\max}\left(ACCOA \cdot OAA - \frac{CIT}{K_{\mathrm{eq}}}\right)}{K_{mACCOA}K_{mOAA}\left(1 + \frac{OAA}{KiOAA_{inhOAA}} + \frac{AKG}{KiOAA_{inhAKG}}\right)}}{\left(1 + \frac{ACCOA}{K_{mACCOA}}\right)\left(1 + \frac{OAA}{K_{mOAA}\left(1 + \frac{OAA}{KiOAA_{inhOAA}} + \frac{AKG}{KiOAA_{inhAKG}}\right)}\right) + \left(1 + \frac{CIT}{K_{mCIT}}\right) - 1} \tag{S22}$$

$$v_{ACN1} = \frac{V_{\max}\left(CIT - \frac{ACO}{K_{\mathrm{eq}}}\right)}{K_{mCIT}\left(1 + \frac{CIT}{K_{mCIT}} + \frac{ACO}{K_{mACO}}\right)} \tag{S23}$$

$$v_{ACN2} = \frac{V_{\max}\left(ACO - \frac{ICIT}{K_{\mathrm{eq}}}\right)}{K_{mACO}\left(1 + \frac{ACO}{K_{mACO}} + \frac{ICIT}{K_{mICIT}}\right)} \tag{S24}$$

$$v_{ICD} = \frac{V_{\max}\left(ICIT - \frac{AKG}{K_{\text{eq}}}\right)}{K_{mICIT}\left(1 + \frac{ICIT}{K_{mICIT}} + \frac{AKG}{K_{mAKG}\left(1+\frac{ICIT}{KiAKG_{inhICIT}}\right)}\right)} \tag{S25}$$

$$v_{LPD} = \frac{V_{\max}\, AKG}{K_{mAKG} + AKG} \tag{S26}$$

$$v_{SK} = \frac{V_{\max}\, SUCCOA}{K_{mSUCCOA} + SUCCOA} \tag{S27}$$

$$v_{SDH} = \frac{V_{\max}\left(SUC - \frac{FUM}{K_{\text{eq}}}\right)}{K_{mSUC}\left(1 + \frac{OAA}{KiSUC_{inhOAA}}\right)\left[1 + \frac{SUC}{K_{mSUC}\left(1+\frac{OAA}{KiSUC_{inhOAA}}\right)} + \frac{FUM}{K_{mFUM}}\right]} \tag{S28}$$

$$\begin{aligned} Den_{FUMA} = 1 + \frac{MAL}{K_{mMAL}} + \frac{FUM}{K_{mFUM}} \\ + \frac{FUM}{1 + \frac{PEP}{KiFUM_{inhPEP}} + \frac{PYR}{KiFUM_{inhPYR}} + \frac{CIT}{KiFUM_{inhCIT}} + \frac{AKG}{KiFUM_{inhAKG}} + \frac{OAA}{KiFUM_{inhOAA}}} \end{aligned} \tag{S29}$$

$$v_{FUMA} = \frac{\frac{V_{\max}\left(FUM - \frac{MAL}{K_{\text{eq}}}\right)}{K_{mFUM} \cdot Den_{FUMA}}}{\left(1 + \frac{PEP}{KiFUM_{inhPEP}} + \frac{PYR}{KiFUM_{inhPYR}} + \frac{CIT}{KiFUM_{inhCIT}} + \frac{AKG}{KiFUM_{inhAKG}} + \frac{OAA}{KiFUM_{inhOAA}}\right)} \tag{S30}$$

$$v_{MDH} = \frac{V_{\max}\left(OAA - \frac{MAL}{K_{\text{eq}}}\right)}{K_{mOAA}\left(1 + \frac{OAA}{K_{mOAA}} + \frac{MAL}{K_{mMAL}}\right)} \tag{S31}$$

#### 1.4 Glyoxylate shunt

$$v_{ICL} = \frac{V_{\max} \cdot ICIT}{K_m ICIT + ICIT} \tag{S32}$$

$$Den_{MALS} = \left[(1 + \frac{ACCOA}{K_{mACCOA}})\left(1 + \frac{GLX}{K_{mGLX}\left(1 + \frac{PYR}{KiGLX_{inhPYR}} + \frac{OAA}{KiGLX_{inhOAA}}\right)}\right) + \left(1 + \frac{MAL}{K_{mMAL}}\right) - 1\right] \tag{S33}$$

$$v_{MALS} = \frac{V_{\max}\left(ACCOA \cdot GLX - \frac{MAL}{K_{\text{eq}}}\right)}{K_{mACCOA} K_{mGLX}\left(1 + \frac{PYR}{KiGLX_{inhPYR}} + \frac{OAA}{KiGLX_{inhOAA}}\right) Den_{MALS}} \tag{S34}$$

#### 1.5 GSGOGAT

$$v_{GD} = \frac{V_{\max}\left(GLU - \frac{AKG \cdot NH4}{K_{\text{eq}}}\right)}{K_{mGLU}\left[1 + \frac{GLU}{K_{mGLU}} + \frac{AKG}{K_{mAKG}\left(1+\frac{GLU}{KiAKG_{inhGLU}}\right)} + \frac{NH4}{K_{mNH4}} + \frac{AKG \cdot NH4}{K_{mAKG}\left(1+\frac{GLU}{KiAKG_{inhGLU}}\right)K_{mNH4}}\right]} \tag{S35}$$

$$Den_{GOGAT} = K_{mAKG}\left(1 + \frac{GLU}{KiAKG_{inhGLU}} + \frac{OAA}{KiAKG_{inhOAA}} + \frac{ASP}{KiAKG_{inhASP}}\right) GLN \tag{S36}$$

$$v_{GOGAT} = \frac{V_{\max} \cdot GLN \cdot AKG}{Den_{GOGAT} + K_{mGLN} \cdot AKG + GLN \cdot AKG} \tag{S37}$$

$$v_{GS} = \frac{\frac{V_{\max}\left(GLU \cdot NH4 - \frac{GLN}{K_{\text{eq}}}\right)}{(1 + \frac{GLN}{KiGLU_{inhGLN}} + \frac{GLN}{KiNH4_{inhGLN}})K_{mGLU}K_{mNH4}}}{(1 + \frac{GLU}{K_{mGLU}})(1 + \frac{NH4}{K_{mNH4}}) + \left(1 + \frac{GLN}{K_{mGLN}(1+\frac{ALN}{KiGLN_{inhALN}}+\frac{GLY}{KiGLN_{inhGLY}})}\right) - 1} \tag{S38}$$

#### 1.6 De novo purine synthesis

$$v_{PRS} = \frac{\frac{V_{\max}\left(R5P\left(1+\frac{PRPP}{KiiR5P_{inhPRPP}}\right)-\frac{PRPP}{K_{\mathrm{eq}}}\right)}{K_{mR5P}\left(1+\frac{PRPP}{KisR5P_{inhPRPP}}\right)}}{\left(1+\frac{R5P\left(1+\frac{PRPP}{KiiR5P_{inhPRPP}}\right)}{K_{mR5P}\left(1+\frac{PRPP}{KisR5P_{inhPRPP}}\right)}+\frac{PRPP}{K_{mPRPP}}\right)} \tag{S39}$$

$$\begin{aligned} Den_{PRAT} = 1 &+ \frac{PRPP}{K_{mPRPP}(1+\frac{IMP}{KiPRPP_{inhIMP}})} + \frac{GLN}{K_{mGLN}(1+\frac{NH4}{KiGLN_{inhNH4}}+\frac{GLU}{KiGLN_{inhGLU}})} \\ &+ \frac{PRA}{K_{mPRA}} + \frac{GLU}{K_{mGLU}} + \frac{PRA\cdot GLU}{K_{mPRA}K_{mGLU}} \\ &+ \frac{PRPP\cdot GLN}{K_{mPRPP}(1+\frac{IMP}{KiPRPP_{inhIMP}})K_{mGLN}(1+\frac{NH4}{KiGLN_{inhNH4}}+\frac{GLU}{KiGLN_{inhGLU}})} \end{aligned} \tag{S40}$$

$$v_{PRAT} = \frac{\frac{V_{\max}\left(PRPP\cdot GLN-\frac{PRA\cdot GLU}{K_{\mathrm{eq}}}\right)}{K_{mPRPP}\left(1+\frac{IMP}{KiPRPP_{inhIMP}}\right)K_{mGLN}\left(1+\frac{NH4}{KiGLN_{inhNH4}}+\frac{GLU}{KiGLN_{inhGLU}}\right)}}{Den_{PRAT}} \tag{S41}$$

$$v_{GARS} = \frac{\frac{V_{\max}\left(PRA\cdot GLY-\frac{GAR}{K_{\mathrm{eq}}}\right)}{K_{mPRA}K_{mGLY}}}{(1+\frac{PRA}{K_{mPRA}})(1+\frac{GLY}{K_{mGLY}})+\left(1+\frac{GAR}{K_{mGAR}}\right)-1} \tag{S42}$$

$$v_{GARTF} = \frac{V_{\max}\cdot GAR\cdot CHOTHF}{K_{mCHOTHF}\left(1+\frac{THF}{KiCHOTHF_{inhTHF}}\right)\cdot GAR+K_{mGAR}\cdot CHOTHF+GAR\cdot CHOTHF} \tag{S43}$$

$$v_{FGAMS} = \frac{V_{\max}\cdot FGAR\left(1+\frac{GLU}{KiiFGAR_{inhGLU}}\right)\cdot GLN}{K_{mGLN}\left(1+\frac{GLU}{KiGLN_{inhGLU}}\right)\cdot FGAR+K_{mFGAR}\left(1+\frac{GLU}{KisFGAR_{inhGLU}}\right)\cdot GLN+FGAR\cdot GLN} \tag{S44}$$

$$v_{AIRS} = \frac{V_{\max}\cdot FGAM}{K_mFGAM+FGAM} \tag{S45}$$

$$v_{CAIRS} = \frac{V_{\max}\cdot AIR}{K_mAIR+AIR} \tag{S46}$$

$$v_{SS} = \frac{V_{\max}\cdot CAIR\cdot ASP}{K_{mASP}\cdot CAIR+Km_{CIAR}\cdot ASP+CIAR\cdot ASP} \tag{S47}$$

$$v_{ADSL} = \frac{\frac{V_{\max}\left(SAICAR-\frac{FUM\cdot AICAR}{K_{\mathrm{eq}}}\right)}{K_{mSAICAR}}}{1+\frac{SAICAR}{K_{mSAICAR}}+\frac{FUM}{K_{mFUM}}+\frac{AICAR}{K_{mAICAR}}+\frac{FUM\cdot AICAR}{K_{mFUM}K_{mAICAR}}} \tag{S48}$$

$$v_{AICARTF} = \frac{V_{\max}\cdot AICAR\cdot CHOTHF}{K_{mCHOTHF}\cdot AICAR+K_{mAICAR}\cdot CHOTHF+AICAR\cdot CHOTHF} \tag{S49}$$

$$v_{IMPCH} = \frac{\frac{V_{\max}\left(FAICAR-\frac{IMP}{K_{\mathrm{eq}}}\right)}{K_{mFAICAR}}}{\left(1+\frac{FAICAR}{K_{mFAICAR}}+\frac{IMP}{K_{mIMP}}\right)} \tag{S50}$$

#### 1.7 Allantoin synthesis

$$v_{IMPDH} = \frac{V_{\max} \cdot IMP}{K_m IMP + IMP} \tag{S51}$$

$$v_{NT} = \frac{V_{\max} \cdot XMP}{K_m XMP + XMP} \tag{S52}$$

$$v_{PNP} = \frac{V_{\max} \cdot XAO}{K_m XAO + XAO} \tag{S53}$$

$$v_{XOR} = \frac{V_{\max} \cdot XAN}{K_{mXAN} \left(1 + \frac{URATE}{KiXAN_{inhURATE}}\right) + XAN} \tag{S54}$$

$$v_{UOD} = \frac{V_{\max} \cdot URATE}{K_{mURATE} \left(1 + \frac{XAN}{KiURATE_{inhXAN}}\right) + URATE} \tag{S55}$$

$$v_{HIUHS} = \frac{V_{\max} \cdot HIUH}{K_m HIUH + HIUH} \tag{S56}$$

$$v_{OHCUD} = \frac{V_{\max} \cdot OHCU}{K_m OHCU + OHCU} \tag{S57}$$

#### 1.8 Aspartate metabolism

$$\begin{aligned} Den_{ATS} = 1 &+ \frac{OAA}{K_{mOAA}(1 + \frac{OAA}{KiOAA_{inhOAA}})} + \frac{GLU}{K_{mGLU}} + \frac{ASP}{K_{mASP}(1 + \frac{GLU}{KiASP_{inhGLU}})} \\ &+ \frac{AKG}{K_{mAKG}(1 + \frac{OAA}{KiAKG_{inhOAA}})} + \frac{OAA \cdot GLU}{K_{mOAA}(1 + \frac{OAA}{KiOAA_{inhOAA}})K_{mGLU}} \\ &+ \frac{ASP \cdot AKG}{K_{mASP}(1 + \frac{GLU}{KiASP_{inhGLU}})K_{mAKG}(1 + \frac{OAA}{KiAKG_{inhOAA}})} \end{aligned} \tag{S58}$$

$$v_{ATS} = \frac{\frac{V_{\max}\left(OAA \cdot GLU - \frac{ASP \cdot AKG}{K_{\mathrm{eq}}}\right)}{K_{mOAA} K_{mGLU}}}{Den_{ATS}} \tag{S59}$$

#### 1.9 Alanine metabolism

$$v_{ALTSS} = \frac{\frac{V_{\max}\left(PYR \cdot GLU - \frac{ALN \cdot AKG}{K_{\mathrm{eq}}}\right)}{K_{mPYR} K_{mGLU}}}{1 + \frac{PYR}{K_{mPYR}} + \frac{GLU}{K_{mGLU}} + \frac{ALN}{K_{mALN}} + \frac{AKG}{K_{mAKG}} + \frac{PYR \cdot GLU}{K_{mPYR} K_{mGLU}} + \frac{ALN \cdot AKG}{K_{mALN} K_{mAKG}}} \tag{S60}$$

$$\begin{aligned} Den_{AGT} = 1 &+ \frac{ALN\left(1 + \frac{PYR}{KiALN_{inhPYR_{Kis}}}\right)}{K_{mALN}\left(1 + \frac{PYR}{KiALN_{inhPYR_{Kii}}}\right)} + \frac{GLX}{K_{mGLX}\left(1 + \frac{PYR}{KiGLC_{inhPYR}}\right)} + \frac{PYK}{K_{mPYK}} + \frac{GLY}{K_{mGLY}} \\ &+ \frac{ALN\left(1 + \frac{PYR}{KiALN_{inhPYR_{Kis}}}\right) \cdot GLX}{K_{mALN}\left(1 + \frac{PYR}{KiALN_{inhPYR_{Kii}}}\right) K_{mGLX}\left(1 + \frac{PYR}{KiGLC_{inhPYR}}\right)} + \frac{PYK \cdot GLY}{K_{mPYK} K_{mGLY}} \end{aligned} \tag{S61}$$

$$v_{AGT} = \frac{\frac{V_{\max}\left(ALN\left(1 + \frac{PYR}{KiALN_{inhPYR_{Kis}}}\right) \cdot GLX - \frac{PYK \cdot GLY}{K_{\mathrm{eq}}}\right)}{K_{mALN}\left(1 + \frac{PYR}{KiALN_{inhPYR_{Kii}}}\right) K_{mGLX}\left(1 + \frac{PYR}{KiGLC_{inhPYR}}\right)}}{Den_{AGT}} \tag{S62}$$

##### 1.10 Serine and glycine metabolism

$$v_{PGDH} = \frac{\frac{V_{\max}\left(3PG - \frac{PHP3}{K_{\text{eq}}}\right)}{K_{m3PG}}}{\left(1 + \frac{3PG}{K_{m3PG}} + \frac{PHP3}{K_{mPHP3}}\right)} \tag{S63}$$

$$v_{PSTS} = \frac{\frac{V_{\max}\left(PHP3 \cdot GLU - \frac{Serine \cdot AKG}{K_{\text{eq}}}\right)}{K_{mPHP3}K_{mGLU}}}{\left(1 + \frac{PHP3}{K_{mPHP3}} + \frac{GLU}{K_{mGLU}} + \frac{Serine}{K_{mSerine}} + \frac{AKG}{K_{mAKG}} + \frac{PHP3 \cdot GLU}{K_{mPHP3}K_{mGLU}} + \frac{Serine \cdot AKG}{K_{mSerine}K_{mAKG}}\right)} \tag{S64}$$

$$\begin{aligned} Den_{SHMT} = 1 &+ \frac{Serine}{K_{mSerine}(1 + \frac{Glycine}{KiSerine_{inhGlycine}})} + \frac{THF}{K_{mTHF}} + \frac{Glycine}{K_{mGlycine}} \\ &+ \frac{CH2THF}{K_{mCH2THF}} + \frac{Serine \cdot THF}{K_{mSerine}(1 + \frac{Glycine}{KiSerine_{inhGlycine}})K_{mTHF}} + \frac{Glycine \cdot CH2THF}{K_{mGlycine}K_{mCH2THF}} \end{aligned} \tag{S65}$$

$$v_{SHMT} = \frac{\frac{V_{\max}\left(Serine \cdot THF - \frac{Glycine \cdot CH2THF}{K_{\text{eq}}}\right)}{(1 + \frac{CH2THF}{KiTHF_{inhCH2THF}})K_{mSerine}(1 + \frac{Glycine}{KiSerine_{inhGlycine}})K_{mTHF}}}{Den_{SHMT}} \tag{S66}$$

$$v_{MTHFR} = \frac{V_{\max} \cdot CH2THF}{K_{mCH2THF}\left(1 + \frac{CH2THF}{KiCH2THF}\right) + CH2THF} \tag{S67}$$

$$v_{MTHFD} = \frac{V_{\max} \cdot CHTHF}{K_{mCHTHF} + CHTHF} \tag{S68}$$

$$v_{GLYDC} = \frac{V_{\max} \cdot Glycine \cdot THF}{K_{mTHF} \cdot Glycine + K_{mGlycine} \cdot THF + Glycine \cdot THF} \tag{S69}$$

##### 1.11 Metabolite exchange

$$v_{MAL_{out}} = \frac{V_{\max} \cdot MAL}{K_m MAL + MAL} \tag{S70}$$

$$v_{SUC_{out}} = \frac{V_{\max} \cdot SUC}{K_m SUC + SUC} \tag{S71}$$

$$v_{FUM_{out}} = \frac{V_{\max} \cdot FUM}{K_m FUM + FUM} \tag{S72}$$

$$v_{ALTN_{out}} = \frac{V_{\max} \cdot Allantoin}{K_m Allantoin + Allantoin} \tag{S73}$$

$$ToNH4 = \frac{V_{max}}{Km_{\text{Bac\_sink}} + Bac_{\text{sink}}} \tag{S74}$$

$$Glycine_{sink} = v_{MAL_{out}} \cdot \frac{0.005}{9.158} \tag{S75}$$

*Bac_sink is the intermediate in the bacteroid that accepts the decarboxylates from the cytosol (malate, glycine) and sends the nitrogen back to the cytosol.

#### 2 ODE systems

$$\frac{dGLC}{dt} = 0 \tag{S76}$$
$$\frac{dG6P}{dt} = v_{HKI} - v_{PGI} - v_{ZWF} \tag{S77}$$
$$\frac{dF6P}{dt} = v_{PGI} - v_{PFK} - v_{FBP} - v_{TKT1} - v_{TAL} \tag{S78}$$
$$\frac{dFDP}{dt} = v_{PFK} - v_{FBA} - v_{FBP} \tag{S79}$$
$$\frac{dGAP}{dt} = v_{FBA} + v_{TPI} - v_{GDH} + v_{TAL} - v_{TKT1} - v_{TKT2} \tag{S80}$$
$$\frac{dDAP}{dt} = v_{FBA} - v_{TPI} \tag{S81}$$
$$\frac{dBPG}{dt} = v_{GDH} - v_{PGK} \tag{S82}$$
$$\frac{d3PG}{dt} = v_{PGK} - v_{GPM} - v_{PGDH} \tag{S83}$$
$$\frac{d2PG}{dt} = v_{GPM} - v_{ENO} \tag{S84}$$
$$\frac{dPEP}{dt} = v_{ENO} - v_{PYK} - v_{PEPC} \tag{S85}$$
$$\frac{dPYR}{dt} = v_{PYK} - v_{PDH} - v_{ALTSS} - v_{AGT} \tag{S86}$$
$$\frac{dGL6P}{dt} = v_{ZWF} - v_{PGL} \tag{S87}$$
$$\frac{dPGN}{dt} = v_{PGL} - v_{GND} \tag{S88}$$
$$\frac{dRB5P}{dt} = v_{GND} - v_{RPE} - v_{RPI} \tag{S89}$$
$$\frac{dR5P}{dt} = v_{RPI} + v_{TKT2} - v_{PRS} \tag{S90}$$
$$\frac{dX5P}{dt} = v_{TKT1} + v_{TKT2} - v_{RPE} \tag{S91}$$
$$\frac{dE4P}{dt} = v_{TKT1} - v_{TAL} \tag{S92}$$
$$\frac{dS7P}{dt} = v_{TAL} - v_{TKT2} \tag{S93}$$
$$\frac{dOAA}{dt} = v_{PEPC} - v_{MDH} - v_{GLT} - v_{ATS} \tag{S94}$$
$$\frac{dMAL}{dt} = v_{MDH} - v_{MAL_{out}} + v_{FUMA} + v_{MALS} \tag{S95}$$
$$\frac{dACCOA}{dt} = v_{PDH} - v_{GLT} - v_{MALS} \tag{S96}$$
$$\frac{dCIT}{dt} = v_{GLT} - v_{ACN1} \tag{S97}$$
$$\frac{dACO}{dt} = v_{ACN1} - v_{ACN2} \tag{S98}$$
$$\frac{dGLX}{dt} = v_{ICL} - v_{MALS} - v_{AGT} \tag{S99}$$
$$\frac{dICIT}{dt} = v_{ACN2} - v_{ICDH} - v_{ICL} \tag{S100}$$
$$\frac{dAKG}{dt} = v_{ICDH} - v_{LPD} - v_{GOGAT} + v_{GD} + v_{PSTS} + v_{ALTSS} + v_{ATS} \tag{S101}$$
$$\frac{dSUCCOA}{dt} = v_{LPD} - v_{SK} \tag{S102}$$
$$\frac{dSUC}{dt} = v_{SK} - v_{SUC_{out}} + v_{ICL} - v_{SDH} \tag{S103}$$
$$\frac{dFUM}{dt} = v_{SDH} - v_{FUM_{out}} - v_{FUMA} + v_{ADSL} \tag{S104}$$

$$\frac{dBac}{dt} = v_{MAL_{out}} + v_{GLY_{sink}} - \frac{9.158 + 0.005}{6.735} \cdot v_{ToNH4} \tag{S105}$$

$$\frac{dNH4}{dt} = v_{ToNH4} - v_{GS} + v_{GD} + v_{GLYDC} \tag{S106}$$

$$\frac{dGLU}{dt} = -v_{GS} + 2 \cdot v_{GOGAT} - v_{GD} - v_{PSTS} + v_{PRAT} - v_{ALTSS} + v_{FGAMS} - v_{ATS} \tag{S107}$$

$$\frac{dGLN}{dt} = v_{GS} - v_{GOGAT} - v_{PRAT} - v_{FGAMS} \tag{S108}$$

$$\frac{dPRPP}{dt} = v_{PRS} - v_{PRAT} \tag{S109}$$

$$\frac{dPHP3}{dt} = v_{PGDH} - v_{PSTS} \tag{S110}$$

$$\frac{dSerine}{dt} = v_{PSTS} - v_{SHMT} \tag{S111}$$

$$\frac{dGlycine}{dt} = v_{SHMT} - v_{GARS} + v_{AGT} - v_{GLYDC} - v_{Glycine_{sink}} \tag{S112}$$

$$\frac{dTHF}{dt} = -v_{SHMT} + v_{GARTF} - v_{GLYDC} + v_{AICARTF} \tag{S113}$$

$$\frac{dCH2THF}{dt} = v_{SHMT} - v_{MTHFR} + v_{GLYDC} \tag{S114}$$

$$\frac{dCHTHF}{dt} = v_{MTHFR} - v_{MTHFD} \tag{S115}$$

$$\frac{dCHOTHF}{dt} = v_{MTHFD} - v_{GARTF} - v_{AICARTF} \tag{S116}$$

$$\frac{dPRA}{dt} = v_{PRAT} - v_{GARS} \tag{S117}$$

$$\frac{dGAR}{dt} = v_{GARS} - v_{GARTF} \tag{S118}$$

$$\frac{dFGAR}{dt} = v_{GARTF} - v_{FGAMS} \tag{S119}$$

$$\frac{dALN}{dt} = v_{ALTSS} - v_{AGT} \tag{S120}$$

$$\frac{dFGAM}{dt} = v_{FGAMS} - v_{AIRS} \tag{S121}$$

$$\frac{dAIR}{dt} = v_{AIRS} - v_{CAIRS} \tag{S122}$$

$$\frac{dSAICAR}{dt} = v_{SS} - v_{ADSL} \tag{S123}$$

$$\frac{dAICAR}{dt} = v_{ADSL} - v_{AICARTF} \tag{S124}$$

$$\frac{dFAICAR}{dt} = v_{AICARTF} - v_{IMPCH} \tag{S125}$$

$$\frac{dIMP}{dt} = v_{IMPCH} - v_{IMPDH} \tag{S126}$$

$$\frac{dXMP}{dt} = v_{IMPDH} - v_{NT} \tag{S127}$$

$$\frac{dXAO}{dt} = v_{NT} - v_{PNP} \tag{S128}$$

$$\frac{dXAN}{dt} = v_{PNP} - v_{XOR} \tag{S129}$$

$$\frac{dURATE}{dt} = v_{XOR} - v_{UOD} \tag{S130}$$

$$\frac{dHIUH}{dt} = v_{UOD} - v_{HIUHS} \tag{S131}$$

$$\frac{dOHCU}{dt} = v_{HIUHS} - v_{OHCUD} \tag{S132}$$

$$\frac{dALTN}{dt} = v_{OHCUD} - v_{Allantoin_{out}} \tag{S133}$$

$$\frac{dASP}{dt} = v_{ATS} - v_{SS} \tag{S134}$$

### 3 Reactions and kinetic parameters

This section details the list of reactions incorporated in the kinetic model, along with their associated kinetic parameters. While the majority of these values were sourced from established literature, parameters for which no experimental data were available, which accounted for 26% of the parameters (74/274, Table S1), were estimated within biochemically plausible ranges.

**Table S1. Parameters and reactions in the kinetic model**

| Subsystem | EC number | Reaction name | Reaction | Parameter | Value | Reference |
|---|---|---|---|---|---|---|
| Glycolysis | 2.7.1.1 | HKI | Glucose+ATP → G6P+ADP | $km_{\text{GLC}}$ | 0.075 | [1] |
| | | | | $km_{\text{G6P}}$ | 0.5 | estimated |
| | | | | $keq$ | 850 | [2] |
| | 5.3.1.9 | PGI | G6P ↔ F6P | $km_{\text{G6P}}$ | 0.27 | [3] |
| | | | | $km_{\text{F6P}}$ | 0.48 | [4] |
| | | | | $keq$ | 0.276 | [5] |
| | | | | $ki_{\text{G6P\_inh6PG}}$ | 0.013 | [3] |
| | 2.7.1.11 | PFK | ATP+F6P → FDP+ADP | $km_{\text{F6P}}$ | 1.5 | [6] |
| | 3.1.3.11 | FBP | FDP → F6P+$PO_4^-$ | $km_{\text{FDP}}$ | 0.25 | estimated |
| | 4.1.2.13 | FBA | FDP ↔ DAP+GAP | $km_{\text{FDP}}$ | 0.0167 | [7] |
| | | | | $km_{\text{GAP}}$ | 0.0605 | [7] |
| | | | | $km_{\text{DAP}}$ | 0.104 | [7] |
| | | | | $keq$ | 0.19 | [8] |
| | | | | $ki_{\text{FDP\_inhR5P}}$ | 2.2 | [7] |
| | 5.3.1.1 | TPI | DAP ↔ GAP | $km_{\text{DAP}}$ | 0.812 | [9] |
| | | | | $km_{\text{GAP}}$ | 0.245 | [9] |
| | | | | $ki_{\text{3PG}}$ | 0.4 | [9] |
| | | | | $ki_{\text{PEP}}$ | 0.661 | [9] |
| | | | | $kcat_{\text{DAP}}$ | 1080 | [10] |
| | | | | $kcat_{\text{GAP}}$ | 6170 | [10] |
| | 1.2.1.12 | GDH | GAP+NAD+$PO_4^-$ ↔ BPG+NADH | $km_{\text{GAP}}$ | 0.074 | [11] |
| | | | | $km_{\text{PO4}}$ | 9 | [11] |
| | | | | $km_{\text{BPG}}$ | 0.036 | [12] |
| | | | | $keq$ | 20 | [13] |
| | 2.7.2.3 | PGK | ADP+BPG ↔ ATP+PGA3 | $km_{\text{BPG}}$ | 0.1 | estimated |
| | | | | $km_{\text{PGA3}}$ | 0.146 | [2] |
| | | | | $keq$ | 100 | estimated |
| | 5.4.2.12 | GPM | PGA3 ↔ PGA2 | $km_{\text{PGA3}}$ | 0.1 | estimated |
| | | | | $km_{\text{PGA2}}$ | 0.369 | [14] |
| | | | | $kcat_{\text{PGA2}}$ | 3.01 | [15] |
| | | | | $kcat_{\text{PGA2}}$ | 4.01 | [15] |
| | 4.2.1.11 | ENO | PGA2 ↔ PEP | $km_{\text{PGA2}}$ | 0.19 | [16] |
| | | | | $km_{\text{PEP}}$ | 0.534 | [17] |
| | | | | $keq$ | 3 | [13] |
| | 2.7.1.40 | PYK | PEP+ADP → PYR+ATP | $km_{\text{PEP}}$ | 0.15 | estimated |
| | 1.2.4.1 | PDH | PYR+COA+NAD → ACCOA+NADH+$HCO_3^-$ | $km_{\text{PYR}}$ | 0.4 | estimated |
| | | | | $km_{\text{COA}}$ | 0.00061 | [18] |
| | | | | $ki_{\text{PYK}}$ | 31 | [19] |
| | 4.1.1.31 | PEPC | PEPC+$HCO_3^-$ → OAA+Pi | $km_{\text{PEP}}$ | 0.09 | [20] |
| | | | | $ki_{\text{MAL}}$ | 37.1 | [20] |
| | | | | $ki_{\text{GLU}}$ | 11.5 | [20] |
| | | | | $ki_{\text{ASP}}$ | 12.6 | [20] |
| Pentose phosphate pathway | 1.1.1.49 | ZWF | G6P+NADP ↔ GL6P+NADPH | $km_{\text{G6P}}$ | 0.1073 | [21] |
| | | | | $ki_{\text{G6P\_inhNH4}}$ | 0.00226 | [22] |
| | | | | $km_{\text{GL6P}}$ | 0.329 | [23] |
| | | | | $keq$ | 1000 | [23] |
| | | | | $kd_{\text{G6P}}$ | 0.192 | [23] |
| | | | | $kd_{\text{GL6P}}$ | 0.02 | [23] |
| | 3.1.1.31 | PGL | GL6P+$H_2O$ ↔ PGN | $km_{\text{GL6P}}$ | 0.83 | [24] |
| | | | | $km_{\text{PGN}}$ | 1 | estimated |
| | | | | $ki_{\text{G6P}}$ | 2 | [23] |
| | | | | $keq$ | 42.7 | [23] |

| Subsystem | EC number | Reaction name | Reaction | Parameter | Value | Reference |
|---|---|---|---|---|---|---|
| | 1.1.1.44 | GND | NADP+PGN → NADPH+RB5P+$HCO_3^-$ | $km_{\mathrm{PGN}}$ | 0.01 | estimated |
| | | | | $km_{\mathrm{RB5P}}$ | 45.2 | [23] |
| | | | | $kd_{\mathrm{RB5P}}$ | 0.044 | [23] |
| | | | | $keq$ | 10 | [23] |
| | 5.3.1.6 | RPI | RB5P ↔ R5P | $km_{\mathrm{RB5P}}$ | 0.01 | estimated |
| | | | | $km_{\mathrm{R5P}}$ | 0.88 | [25] |
| | | | | $keq$ | 0.333 | [26] |
| | | | | $ki_{\mathrm{R5P\_inhE4P}}$ | 0.1 | [25] |
| | | | | $ki_{\mathrm{RB5P\_inhE4P}}$ | 0.14 | [25] |
| | | | | $ki_{\mathrm{R5P\_inhGAP}}$ | 0.5 | [25] |
| | | | | $ki_{\mathrm{R5P\_inhPGA3}}$ | 1.2 | [25] |
| | | | | $ki_{\mathrm{R5P\_inhPGN}}$ | 7 | [25] |
| | 5.1.3.1 | RPE | RB5P ↔ X5P | $km_{\mathrm{X5P}}$ | 0.067 | estimated |
| | | | | $km_{\mathrm{RB5P}}$ | 2.5 | estimated |
| | | | | $keq$ | 1 | estimated |
| | 2.2.1.2 | TAL | S7P+GAP ↔ E4P+F6P | $km_{\mathrm{F6P}}$ | 1.2 | [27] |
| | | | | $km_{\mathrm{E4P}}$ | 0.09 | [27] |
| | | | | $km_{\mathrm{S7P}}$ | 0.285 | [27] |
| | | | | $km_{\mathrm{GAP}}$ | 0.038 | [27] |
| | | | | $keq$ | 26.6266 | [27] |
| | 2.2.1.1 | TKT1 | GAP+F6P ↔ E4P+X5P | $km_{\mathrm{TKT1\_P5P}}$ | 0.616 | [28] |
| | | | | $km_{\mathrm{TKT2\_P5P}}$ | 0.118 | [28] |
| | | | | $km_{\mathrm{TKT1\_GAP}}$ | 0.2727 | [28] |
| | | | | $km_{\mathrm{TKT1\_F6P}}$ | 0.5443 | [28] |
| | | | | $keq$ | 1 | [28] |
| | | | | $kcat$ | 131.2 | [29] |
| | 2.2.1.1 | TKT2 | GAP+S7P ↔ R5P+X5P | $km_{\mathrm{S7P}}$ | 0.01576 | [28] |
| | | | | $km_{\mathrm{GAP}}$ | 0.09078 | [28] |
| | | | | $keq$ | 1000 | estimated |
| | | | | $kcat$ | 69.05 | [29] |
| TCA cycle | 2.3.3.1 | GLT | ACCOA+OAA+$H_2O$ ↔ CIT+COA | $km_{\mathrm{ACCOA}}$ | 0.12 | estimated |
| | | | | $km_{\mathrm{OAA}}$ | 0.02 | estimated |
| | | | | $km_{\mathrm{CIT}}$ | 1.16 | [23] |
| | | | | $km_{\mathrm{COA}}$ | 0.0001 | [23] |
| | | | | $ki_{\mathrm{OAA\_inhOAA}}$ | 0.033 | [30] |
| | | | | $ki_{\mathrm{OAA\_inh2kg}}$ | 0.76 | [30] |
| | | | | $keq$ | 8300 | [31] |
| | 4.2.1.3 | ACN1 | CIT ↔ ACO+$H_2O$ | $km_{\mathrm{CIT}}$ | 0.1 | estimated |
| | | | | $km_{\mathrm{ACO}}$ | 0.3 | estimated |
| | | | | $kcat_{\mathrm{CIT}}$ | 4.3 | [32] |
| | | | | $kcat_{\mathrm{ACO}}$ | 5 | [32] |
| | 4.2.1.3 | ACN2 | ACO ↔ ICIT | $km_{\mathrm{ACO}}$ | 0.2 | estimated |
| | | | | $km_{\mathrm{ICIT}}$ | 0.58 | [33] |
| | | | | $kcat_{\mathrm{ACO}}$ | 5.2 | [32] |
| | | | | $kcat_{\mathrm{ICIT}}$ | 1.1 | estimated |
| | 1.1.1.42 | ICDH | ICIT+NADP ↔ Akg+$CO_2$+NADPH+$H^+$ | $km_{\mathrm{ICIT}}$ | 0.01 | estimated |
| | | | | $km_{\mathrm{AKG}}$ | 0.5 | estimated |
| | | | | $ki_{\mathrm{AKG\_inhICIT}}$ | 0.012 | [34] |
| | | | | $kcat_{\mathrm{ICIT}}$ | 51.8 | [35] |
| | | | | $kcat_{\mathrm{AKG}}$ | 172.667 | [35] |
| | 1.2.4.2 | LPD | COA+AKG+NAD → NADH+SUCCOA+$HC)_3^-$ | $km_{\mathrm{AKG}}$ | 10 | estimated |
| | 6.2.1.5 | SK | ADP+SUCCOA+$PO_4^-$ ↔ ATP+SUC+COA | $km_{\mathrm{SUCCOA}}$ | 0.41 | estimated |
| | | | | $km_{PO\_4^-}$ | 0.72 | [36] |
| | | | | $km_{\mathrm{SUC}}$ | 1.5 | [37] |
| | | | | $km_{\mathrm{COA}}$ | 0.1 | [37] |
| | | | | $kcat_{\mathrm{SUCCOA}}$ | 22 | [38] |
| | | | | $kcat_{\mathrm{SUC}}$ | 29 | [38] |
| | | | | $keq$ | 1000 | estimated |
| | 1.3.5.1 | SDH | Q+SUC↔ FUM+$QH_2$ | $km_{\mathrm{SUC}}$ | 0.005 | estimated |
| | | | | $km_{\mathrm{FUM}}$ | 0.02 | [39] |
| | | | | $ki_{\mathrm{SUC\_inhOAA}}$ | 0.00007 | [39] |
| | | | | $kcat_{\mathrm{SUC}}$ | 15 | estimated |

| Subsystem | EC number | Reaction name | Reaction | Parameter | Value | Reference |
|---|---|---|---|---|---|---|
| | | | | $kcat_{\text{FUM}}$ | 32 | estimated |
| | | | | $keq$ | 1000 | estimated |
| | 4.2.1.2 | FUMA | FUM↔MAL | $km_{\text{FUM}}$ | 0.1 | estimated |
| | | | | $km_{\text{MAL}}$ | 0.3 | estimated |
| | | | | $ki_{\text{FUM\_inhPEP}}$ | 0.7 | [40] |
| | | | | $ki_{\text{FUM\_inhPYR}}$ | 1.6 | [40] |
| | | | | $ki_{\text{FUM\_inhCIT}}$ | 0.5 | [40] |
| | | | | $ki_{\text{FUM\_inhAKG}}$ | 0.8 | [40] |
| | | | | $ki_{\text{FUM\_inhOAA}}$ | 1.2 | [40] |
| | | | | $kcat_{\text{FUM}}$ | 21 | [40] |
| | | | | $kcat_{\text{MAL}}$ | 15.3 | [40] |
| | | | | $keq$ | 310 | [41] |
| | 1.1.1.37 | MDH | OAA+$NAD^+$ ↔ MAL+NADH+$H^+$ | $km_{\text{OAA}}$ | 0.018 | [42] |
| | | | | $km_{\text{MAL}}$ | 2.6 | [42] |
| | | | | $kcat_{\text{OAA}}$ | 960 | [43] |
| | | | | $kcat_{\text{MAL}}$ | 250 | [43] |
| | | | | $keq$ | 0.5 | estimated |
| Glyoxylate cycle | 4.1.3.1 | ICL | ICIT → GLX+SUC | $km_{\text{ICIT}}$ | 8 | estimated |
| | 2.3.3.9 | MALS | ACCOA+GLX ↔ MAL+COA | $km_{\text{MAL}}$ | 8 | estimated |
| | | | | $km_{\text{ACCOA}}$ | 0.8 | estimated |
| | | | | $km_{\text{GLX}}$ | 30 | estimated |
| | | | | $ki_{\text{GLX\_inhPYR}}$ | 0.54 | [44] |
| | | | | $ki_{\text{GLX\_inhOAA}}$ | 1.5 | [44] |
| | | | | $keq$ | 100 | estimated |
| GS/GOGAT | 1.4.1.3 | GD | GLU+$H_2O$+NAD(P) ↔ AKG+NH3+NAD(P)H+$H^+$ | $km_{\text{AKG}}$ | 2.1 | [45] |
| | | | | $km_{\text{NH4}}$ | 15.8 | [45] |
| | | | | $km_{\text{GLU}}$ | 1.67 | estimated |
| | | | | $kcat_{\text{AKG}}$ | 165.3 | [46] |
| | | | | $kcat_{\text{GLU}}$ | 121.2 | [46] |
| | | | | $ki_{\text{AKG\_inhGLU}}$ | 20 | [45] |
| | 6.3.1.2 | GS | GLU+NH4+ATP ↔ GLN+ADP+$PO_4^-$ | $km_{\text{GLU}}$ | 9 | [47] |
| | | | | $km_{\text{NH4}}$ | 5.2 | [47] |
| | | | | $km_{\text{GLN}}$ | 1.3 | [48] |
| | | | | $ki_{\text{GLU\_inhGLN}}$ | 6.6 | [47] |
| | | | | $ki_{\text{NH4\_inhGLN}}$ | 7.4 | [47] |
| | | | | $ki_{\text{GLN\_inhALN}}$ | 0.03 | [48] |
| | | | | $ki_{\text{GLN\_inhGLY}}$ | 0.1 | [48] |
| | | | | $keq$ | 1000 | estimated |
| | 1.4.1.14 | GOGAT | GLN+AKG+NADH+$H^+$ ↔ 2GLU+$NAD^+$ | $km_{\text{AKG}}$ | 0.039 | [49] |
| | | | | $km_{\text{GLN}}$ | 4 | estimated |
| | | | | $ki_{\text{AKG\_inhGLU}}$ | 0.7 | [50] |
| | | | | $ki_{\text{AKG\_inhOAA}}$ | 5 | [50] |
| | | | | $ki_{\text{AKG\_inhASP}}$ | 2.7 | [50] |
| Alanine metabolism | 2.6.1.2 | ALTSS | ALN+AKG ↔ PYR+GLU | $km_{\text{PYR}}$ | 0.11 | [51] |
| | | | | $km_{\text{GLU}}$ | 0.68 | [51] |
| | | | | $km_{\text{ALN}}$ | 0.4 | [51] |
| | | | | $km_{\text{AKG}}$ | 0.25 | [51] |
| | | | | $kcat_{\text{ALN}}$ | 4.98 | [51] |
| | | | | $kcat_{\text{PYR}}$ | 6.86 | [51] |
| | 2.6.1.44 | AGT | ALN+GLX ↔ PYR+GLY | $km_{\text{ALN}}$ | 0.31 | estimated |
| | | | | $km_{\text{GLX}}$ | 0.23 | [52] |
| | | | | $km_{\text{PYR}}$ | 0.21 | [52] |
| | | | | $km_{\text{GLY}}$ | 22 | [52] |
| | | | | $ki_{\text{GLX\_inhPYR}}$ | 2.3 | [52] |
| | | | | $ki_{\text{ALN\_inhPYR\_Kis}}$ | 15.8 | [52] |
| | | | | $ki_{\text{ALN\_inhPYR\_Kii}}$ | 22.8 | [52] |
| | | | | $kcat_{\text{ALN}}$ | 45 | [52] |
| | | | | $kcat_{\text{GLY}}$ | 10.33 | [52] |
| Aspartate metabolism | 2.6.1.1 | ATS | OAA+GLU ↔ ASP+AKG | $km_{\text{OAA}}$ | 0.02 | [53] |

| Subsystem | EC number | Reaction name | Reaction | Parameter | Value | Reference |
|---|---|---|---|---|---|---|
| | | | | $km_{\text{GLU}}$ | 12 | [53] |
| | | | | $km_{\text{ASP}}$ | 0.05 | estimated |
| | | | | $km_{\text{AKG}}$ | 0.2 | [53] |
| | | | | $ki_{\text{AKG\_inhGLU}}$ | 10 | [53] |
| | | | | $ki_{\text{ASP\_inhGLU}}$ | 12 | [53] |
| | | | | $ki_{\text{ASP\_inhOAA}}$ | 0.027 | [53] |
| | | | | $ki_{\text{AKG\_inhOAA}}$ | 0.056 | [53] |
| | | | | $ki_{\text{OAA\_inhOAA}}$ | 0.2 | [53] |
| | | | | $Vmax_{\text{OAA}}$ | 320 | [53] |
| | | | | $Vmax_{\text{AKG}}$ | 160 | [53] |
| Serine and Glycine metabolism | 1.1.1.95 | PGDH | 3PG+$NAD^+$ ↔ 3PHP+NADH+$H^+$ | $km_{\text{3PG}}$ | 0.29 | [54] |
| | | | | $km_{\text{3PHP}}$ | 0.1 | estimated |
| | | | | $keq$ | 10 | estimated |
| | 2.6.1.52 | PSTS | 3PHP+GLU ↔ Serine+AKG | $km_{\text{3PHP}}$ | 0.08 | estimated |
| | | | | $km_{\text{GLU}}$ | 5 | estimated |
| | | | | $km_{\text{Serine}}$ | 0.225 | [55] |
| | | | | $km_{\text{AKG}}$ | 0.0651 | [53] |
| | | | | $kcat_{\text{3PHP}}$ | 1.75 | [56] |
| | | | | $kcat_{\text{AKG}}$ | 0.63 | [56] |
| | 2.1.2.1 | SHMT | Serine+THF ↔ GLY+5, $10CH_2THF$+$H_2O$ | $km_{\text{Serine}}$ | 2.5 | estimated |
| | | | | $km_{\text{THF}}$ | 0.25 | [57] |
| | | | | $km_{\text{GLY}}$ | 0.66 | [58] |
| | | | | $km_{\text{CH2THF}}$ | 0.98 | [58] |
| | | | | $ki_{\text{Serine\_inhGLY}}$ | 3 | [57] |
| | | | | $ki_{\text{THF\_inhCH2THF}}$ | 2.9 | [57] |
| | | | | $kcat_{\text{THF}}$ | 14.17 | [58] |
| | | | | $kcat_{\text{GLY}}$ | 0.188 | [58] |
| | | | | $keq$ | 10 | estimated |
| | 1.5.1.15 | MTHFR | $CH_2THF$+$NAD^+$ → CHTHF+NADH+$H^+$ | $km_{CH_2THF}$ | 0.0004 | [59] |
| | | | | $ki_{CH_2THF}$ | 0.061 | [59] |
| | 3.5.4.9 | MTHFD | $CHTHF$+$H_2O$ → CHOTHF | $km_{\text{CHTHF}}$ | 0.026 | [60] |
| | 1.4.4.2 | GLYDC | GLY+THF+$NAD^+$ → $CH_2THF$+NADH+$CO_2$+$NH_3$ | $km_{\text{GLY}}$ | 6 | [61] |
| | | | | $km_{\text{THF}}$ | 7 | estimated |
| | | | | $ki_{\text{GLY\_inhSerine}}$ | 4 | [62] |
| de novo purine synthesis | 2.7.6.1 | PRS | R5P+ATP → PRPP+AMP | $km_{\text{R5P}}$ | 0.11 | [63] |
| | | | | $km_{\text{PRPP}}$ | 0.5 | estimated |
| | | | | $kis_{\text{R5P\_inhPRPP}}$ | 0.82 | [63] |
| | | | | $kii_{\text{R5P\_inhPRPP}}$ | 1.7 | [63] |
| | | | | $keq$ | 0.1 | estimated |
| | 2.4.2.14 | PRAT | GLN+PRPP+H2O → PRA+Ppi+GLU | $km_{\text{GLN}}$ | 18 | [64] |
| | | | | $km_{\text{PRPP}}$ | 0.4 | [64] |
| | | | | $km_{\text{PRA}}$ | 0.8 | estimated |
| | | | | $km_{\text{GLU}}$ | 20 | estimated |
| | | | | $ki_{\text{GLN\_inhNH4}}$ | 16 | [64] |
| | | | | $ki_{\text{GLN\_inhGLU}}$ | 30 | [64] |
| | | | | $ki_{\text{PRPP\_inhIMP}}$ | 1.7 | [64] |
| | | | | $ki_{\text{PRPP\_inhXMP}}$ | 1.2 | [64] |
| | | | | $keq$ | 1000 | estimated |
| | 6.3.4.13 | GARS | PRA+ATP+GLY ↔ ADP+$PO_4^-$+beta-GAR | $km_{\text{PRA}}$ | 7 | estimated |
| | | | | $km_{\text{GLY}}$ | 0.27 | [65] |
| | | | | $km_{\text{GAR}}$ | 0.3 | estimated |
| | | | | $km_{\text{PO4}}$ | 0.54 | [65] |
| | | | | $kcat_{\text{GLY}}$ | 7 | [66] |
| | | | | $kcat_{\text{GAR}}$ | 2.3 | [66] |
| | 2.1.2.2 | GARTF | CHOTHF10+beta-GAR → Tthdf+FGAR | $km_{\text{CHOTHF}}$ | 0.2 | estimated |
| | | | | $km_{\text{GAR}}$ | 19.2 | estimated |
| | | | | $ki_{\text{CHOTHF\_inhTHF}}$ | 0.012 | [67] |
| | 6.3.5.3 | FGAMS | FGAR+GLN+ATP+$H_2$) → GLU+FGAM+ADP+$PO_4^-$ | $km_{\text{FGAR}}$ | 4.68 | estimated |

| Subsystem | EC number | Reaction name | Reaction | Parameter | Value | Reference |
|---|---|---|---|---|---|---|
| | | | | $km_{\text{GLN}}$ | 0.03 | [68] |
| | | | | $ki_{\text{GLN\_inhGLU}}$ | 1.6 | [68] |
| | | | | $kis_{\text{FGAR\_inhGLU}}$ | 13 | [68] |
| | | | | $kii_{\text{FGAR\_inhGLU}}$ | 52 | [68] |
| | 6.3.3.1 | AIRS | FGAM+ATP $\rightarrow$ AIR+$PO_4^-$ | $km_{\text{FGAM}}$ | 2.7 | estimated |
| | | | | $kis_{\text{FGAM\_inhAIR}}$ | 0.0647 | [69] |
| | | | | $kii_{\text{FGAM\_inhAIR}}$ | 0.0654 | [69] |
| | | | | $km_{\text{AIR}}$ | 0.76 | estimated |
| | | | | $km_{\text{CAIR}}$ | 1 | [70] |
| | | | | $kcat_{\text{AIR}}$ | 40 | [70] |
| | 4.1.1.21 | CAIRS | AIR+$CO_2$ $\leftrightarrow$ CAIR | $kcat_{\text{CAIR}}$ | 20 | [70] |
| | 6.3.2.6 | SS | ATP+CAIR+ASP $\leftrightarrow$ ADP+SAICAR+$PO_4^-$ | $km_{\text{CAIR}}$ | 0.5 | estimated |
| | | | | $km_{\text{ASP}}$ | 1.4 | [70] |
| | 4.3.2.2 | ADSL | SAICAR $\leftrightarrow$ FUM+AICAR | $km_{\text{SAICAR}}$ | 1 | estimated |
| | | | | $km_{\text{FUM}}$ | 0.52 | [71] |
| | | | | $km_{\text{AICAR}}$ | 0.009 | [72] |
| | | | | $kcat_{\text{SAICAR}}$ | 337 | [73] |
| | | | | $kcat_{\text{FUM}}$ | 2.9 | [74] |
| | 2.1.2.3 | AICARTF | CHOTHF+AICAR $\rightarrow$ FAICAR+TTHDF | $km_{\text{CHOTHF}}$ | 0.2 | [75] |
| | | | | $km_{\text{AICAR}}$ | 0.31 | [75] |
| | | | | $kcat$ | 4.97 | [76] |
| | 3.5.4.10 | IMPCH | FAICAR $\leftrightarrow$ IMP+$H_2$O | $km_{\text{FAICAR}}$ | 0.78 | estimated |
| | | | | $km_{\text{IMP}}$ | 0.5 | estimated |
| | | | | $kcat_{\text{FAICAR}}$ | 1.32 | [77] |
| | | | | $kcat_{\text{IMP}}$ | 0.5 | estimated |
| Allantoin biosynthesis | 1.1.1.205 | IMPDH | IMP+NAD+$H_2$O $\rightarrow$ XMP+NADH+$H^+$ | $km_{\text{IMP}}$ | 0.5 | estimated |
| | | | | $kcat$ | 6.1 | [78] |
| | 3.1.3.5 | NT | XMP+$H_2$O $\rightarrow$ Xanthosine+$PO_4^-$ | $km_{\text{XMP}}$ | 0.77 | [79] |
| | 2.4.2.1 | PNP | Xanthosine+$PO_4^-$ $\rightarrow$ Xanthine+R1P | $km_{\text{XAO}}$ | 0.51 | estimated |
| | | | | $km_{\text{PO4}}$ | 0.76 | [80] |
| | 1.17.1.4 | XOR | Xanthine+NAD+$H_2$O $\rightarrow$ Urate+NADH+$H^+$ | $km_{\text{XAN}}$ | 0.005 | [81] |
| | | | | $ki_{\text{XAN\_inhUrate}}$ | 0.18 | [81] |
| | 1.7.3.3 | UOD | Urate+$O_2$+$H_2$O $\rightarrow$ HDSS+$H_2$O$_2$ | $km_{\text{Urate}}$ | 0.01 | [82] |
| | | | | $km_{\text{O2}}$ | 0.031 | [82] |
| | | | | $ki_{\text{Urate\_inhXAN}}$ | 0.01 | [82] |
| | 3.5.2.17 | HIUHS | HDS5+$H_2$O $\rightarrow$ OHCU | $km_{\text{HIUH}}$ | 0.015 | [83] |
| | 4.1.1.97 | OHCUD | OHCU $\rightarrow$ Allantoin+$CO_2$ | $km_{\text{OHCU}}$ | 0.151 | [84] |
| Metabolite exchange | NA | MAL_out | $MAL_{\text{cytosol}} \rightarrow MAL_{\text{bacteroid}}$ | $km_{\text{MAL}}$ | 1 | estimated |
| | NA | SUC_out | $SUC_{\text{cytosol}} \rightarrow SUC_{\text{bacteroid}}$ | $km_{\text{SUC}}$ | 2.5 | estimated |
| | NA | FUM_out | $FUM_{\text{cytosol}} \rightarrow FUM_{\text{bacteroid}}$ | $km_{\text{FUM}}$ | 3.91 | estimated |
| | NA | ALTN_out | $ALTN$cytosol $\rightarrow ALTN_{\text{root}}$ | $km_{\text{ALTN}}$ | 0.0762 | [85] |
| | NA | $ToNH_4$ | $Bac_{\text{sink}} \rightarrow NH_4$ | $km_{Bac_{\text{sink}}}$ | 0.5 | estimated |

### 4 Supplemental figures and tables

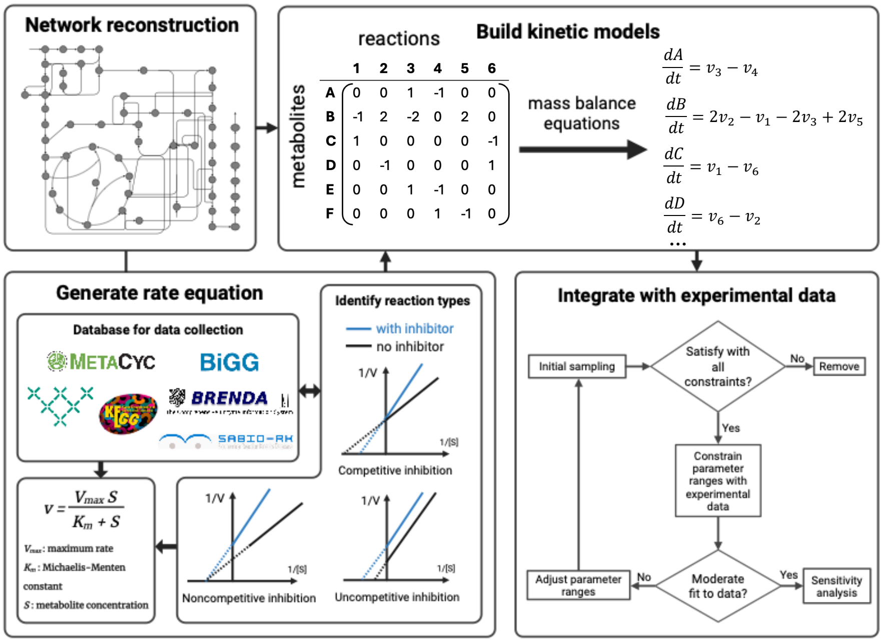

**Figure S1. Overall framework used to construct the kinetic model for the determinate nodule and incorporate experimental data.**

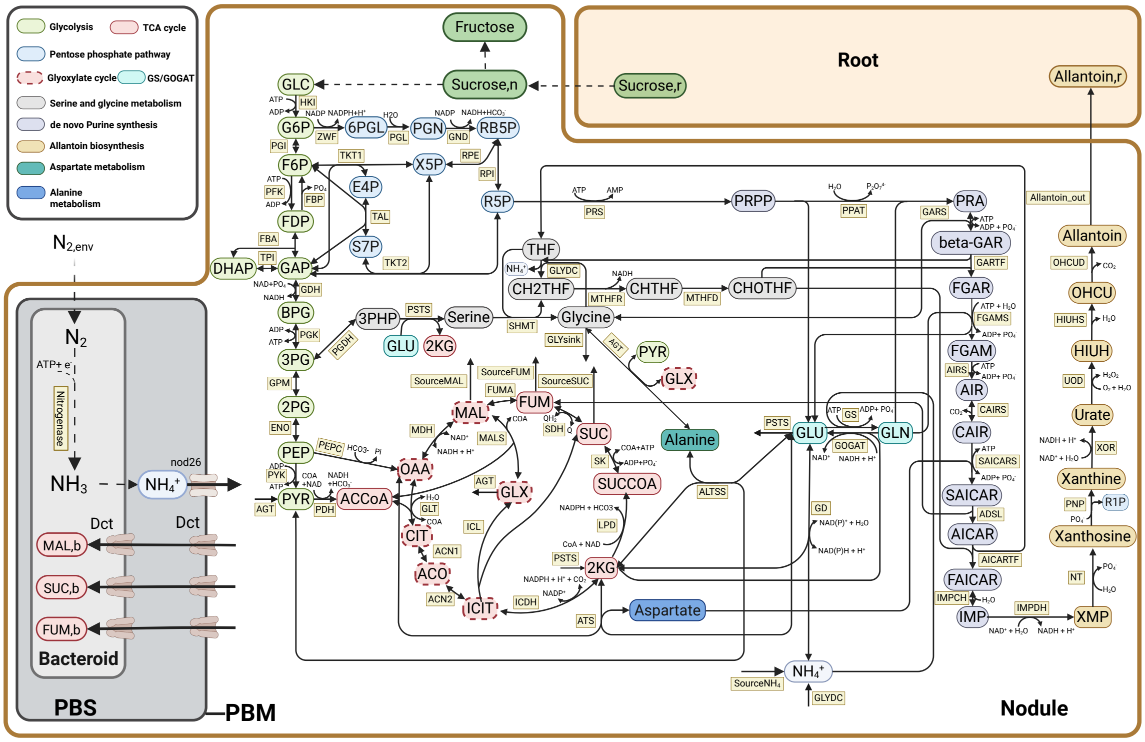

**Figure S2. Pathway reactions included in the kinetic models of determinate root nodule.** All pathways' reversibility information is included in this flux map. Metabolites that participated in each subnetwork are shown in the corresponding colors. Enzymes are shown in yellow. Black arrows denote reactions and regulatory interactions, respectively.

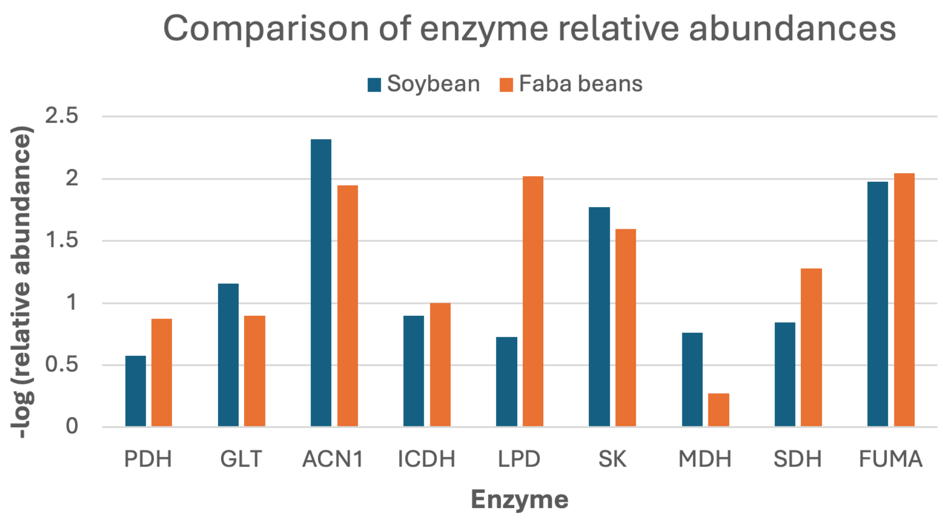

**Figure S3. Relative abundance of enzymes in the TCA cycle in soybean [86] and faba beans [87] datasets.**

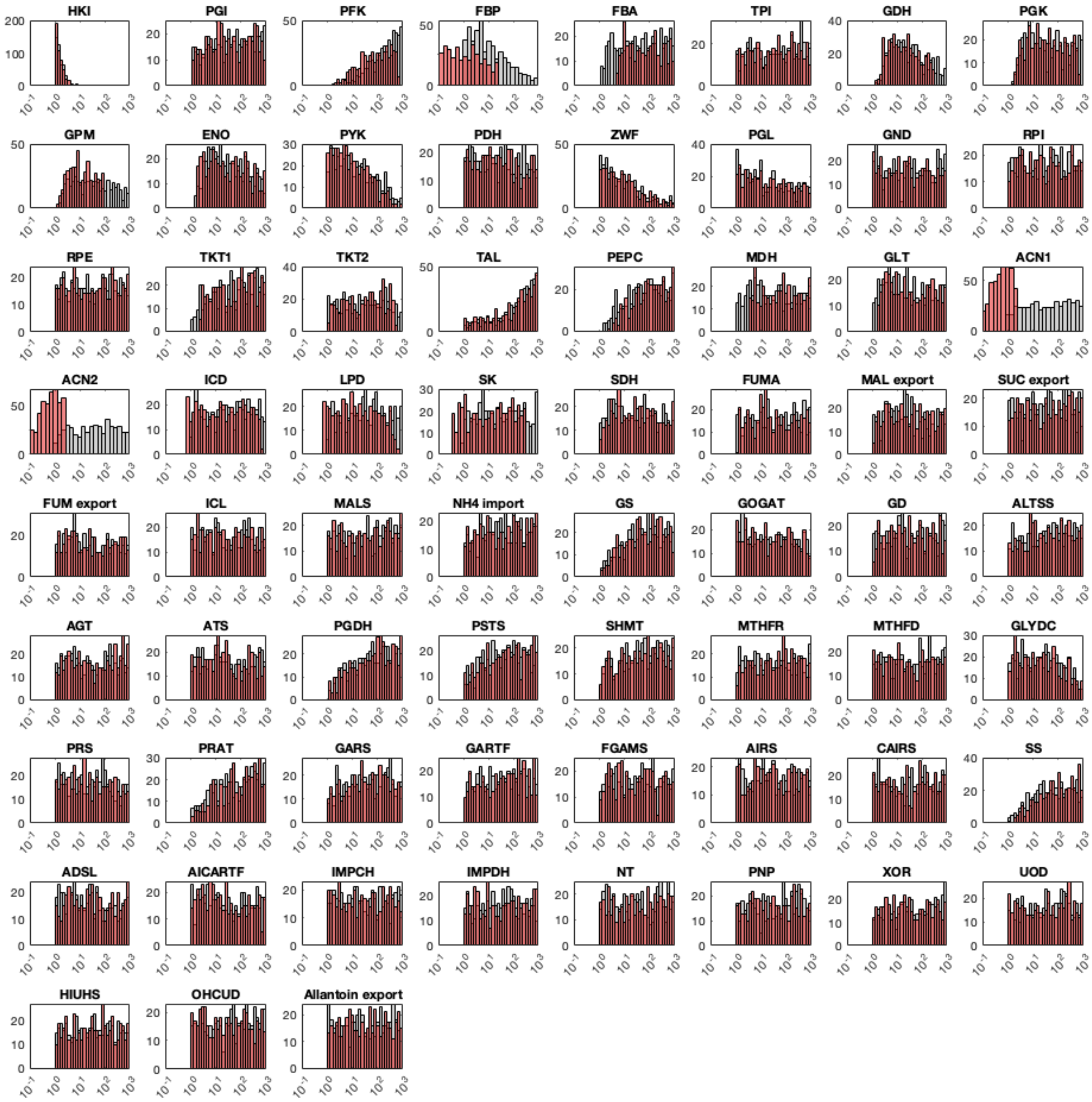

**Figure S4. Comparison of Vmax distributions before (grey) and after (red) applying soybean and faba bean proteomic data.** In the distribution plots, enzymes HKI exhibited the most significant skewness within the original parameter space. The sampling process effectively captures the full range of biologically meaningful enzyme distributions with the applied constraints.

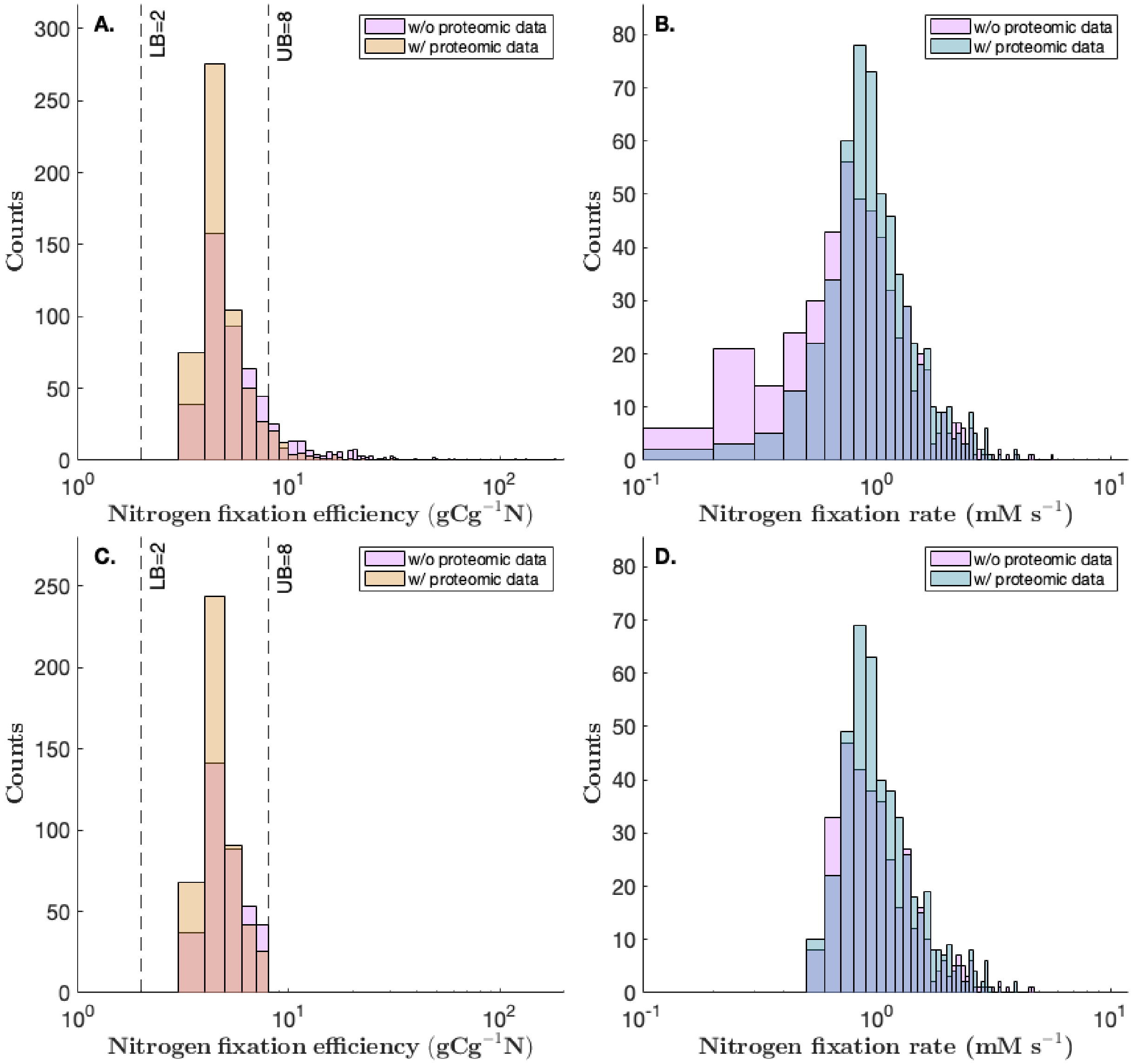

**Figure S5. The predicted N-fixation efficiency and N-fixation rate distributions comparing to whether incorporating proteomic data (pink) or not.**

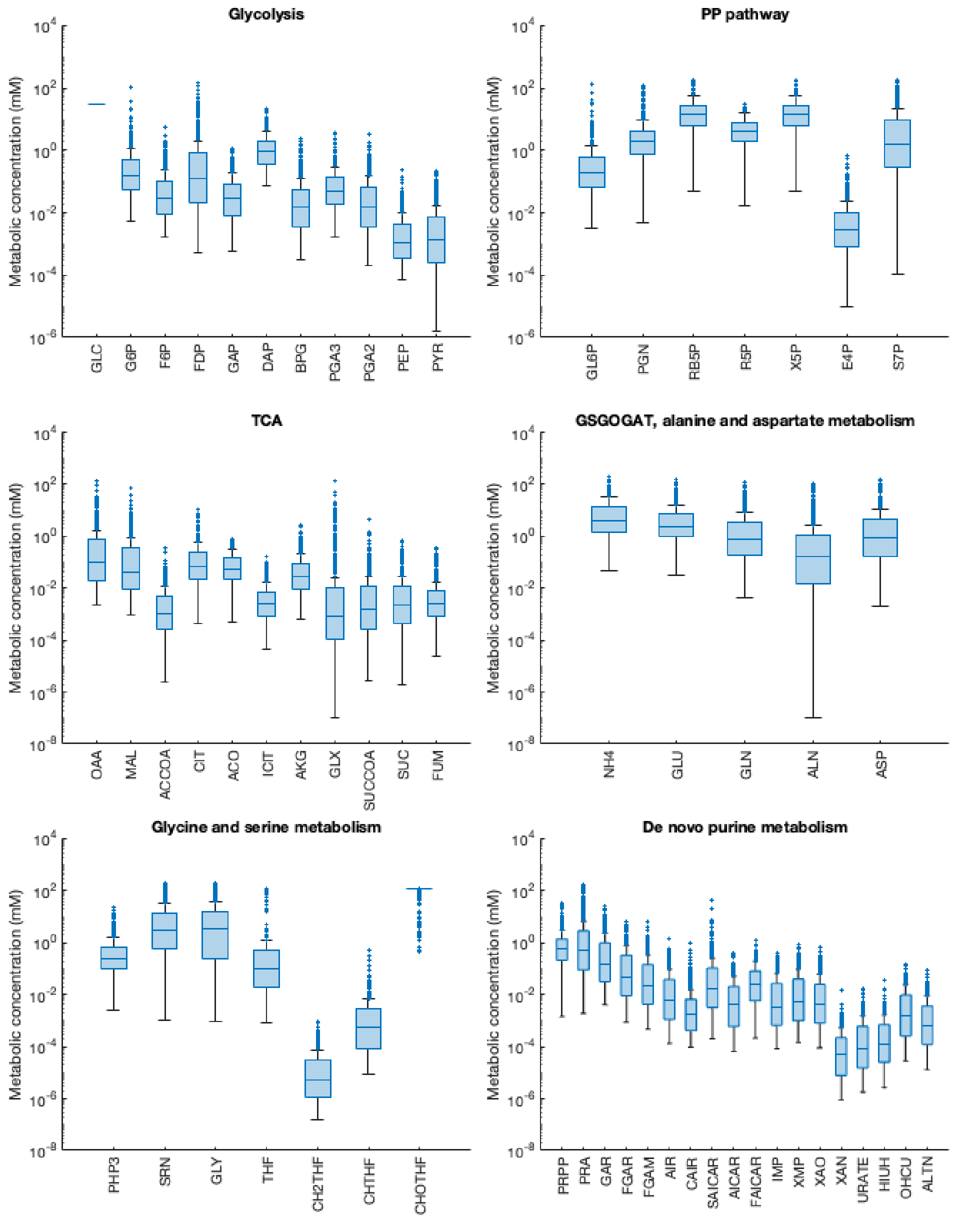

**Figure S6. The distribution of metabolic concentration at steady state.**

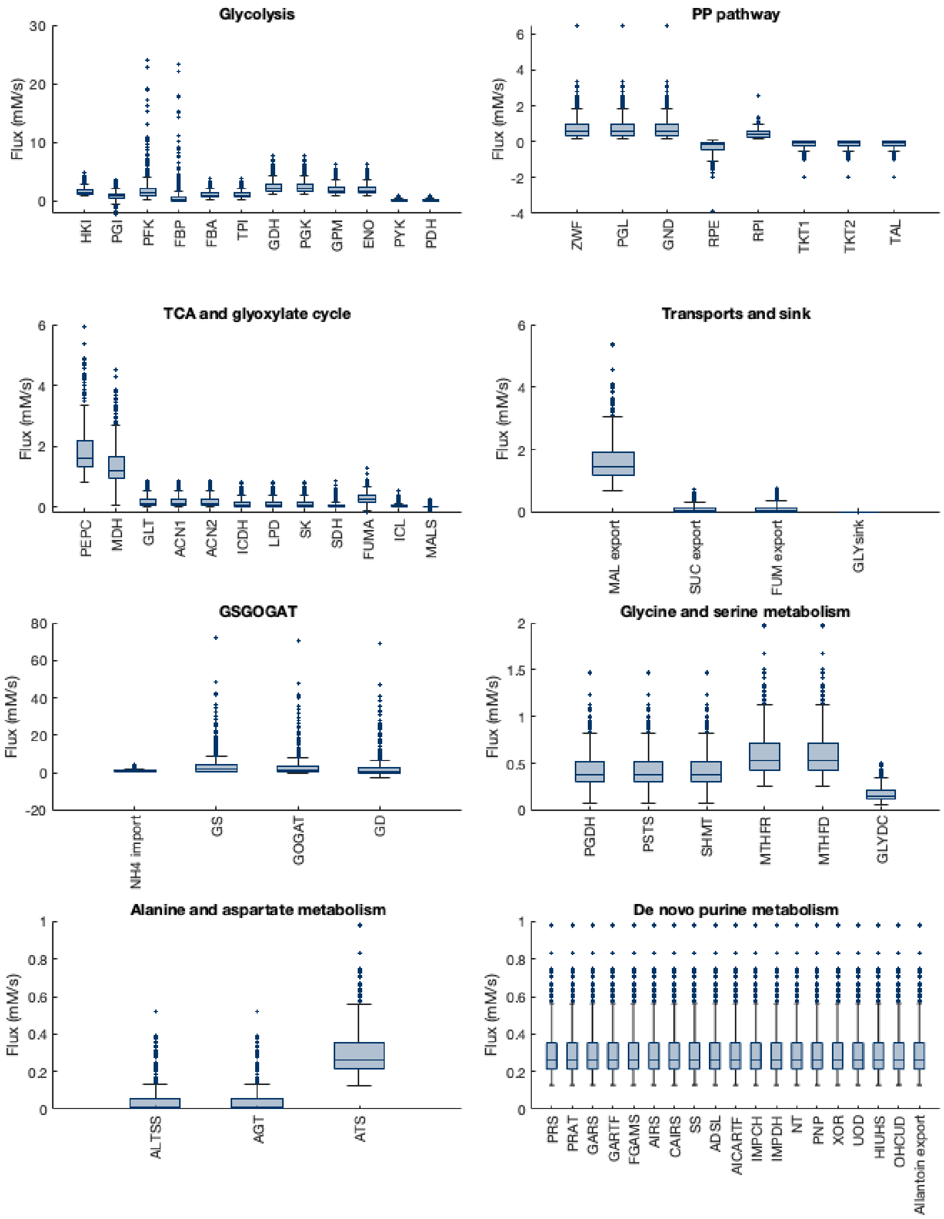

**Figure S7. The distribution of reaction's flux at steady state.**

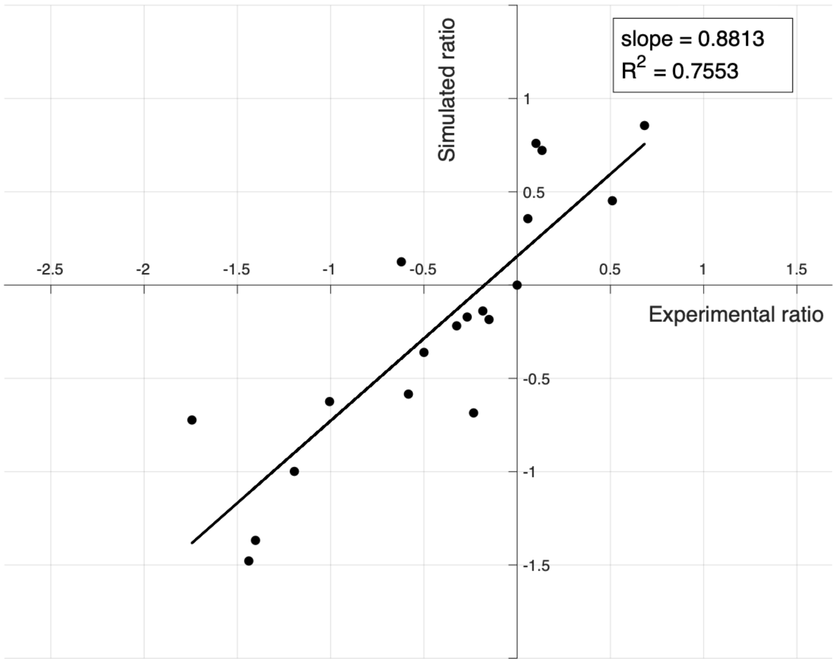

**Figure S8. Model fitness with experimental and simulation data of protein relative proportions.** The ratios are calculated by normalizing enzyme concentration by PDH.

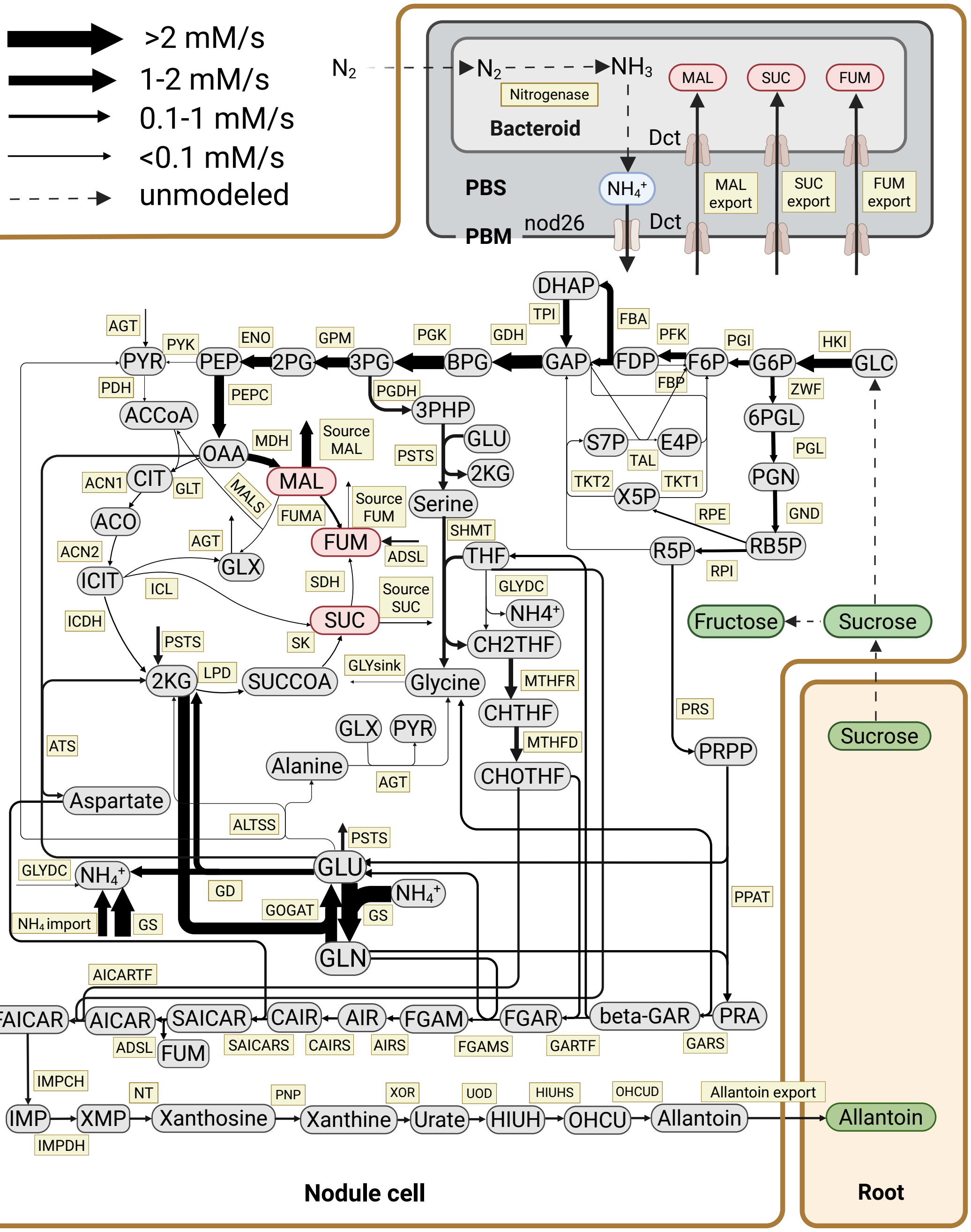

**Figure S9. The flux map for the bacteroid with less dicarboxylate demands (23% increase in bacteroid efficiency ($gCg^{-1}N = 3.25$), N-fixation rate = 1.33 $mMs^{-1}$).** Reaction arrows shown in different colors represent the different magnitudes of fluxes.

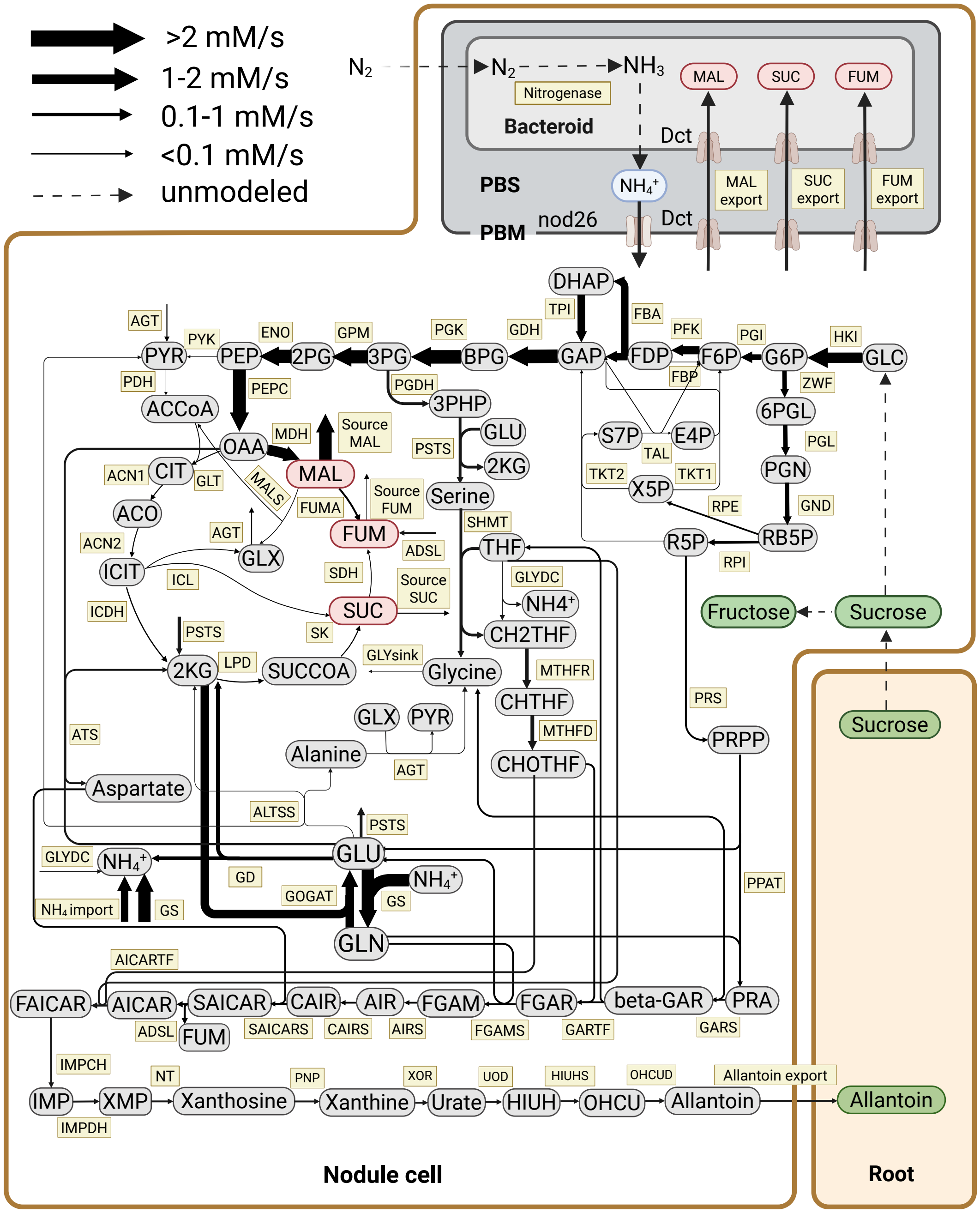

**Figure S10. The flux map for the bacteroid with more dicarboxylate demands (23% decrease in bacteroid efficiency ($gCg^{-1}N = 5.08$), N-fixation rate = 0.99 $mMs^{-1}$).** Reaction arrows shown in different colors represent the different magnitudes of fluxes.

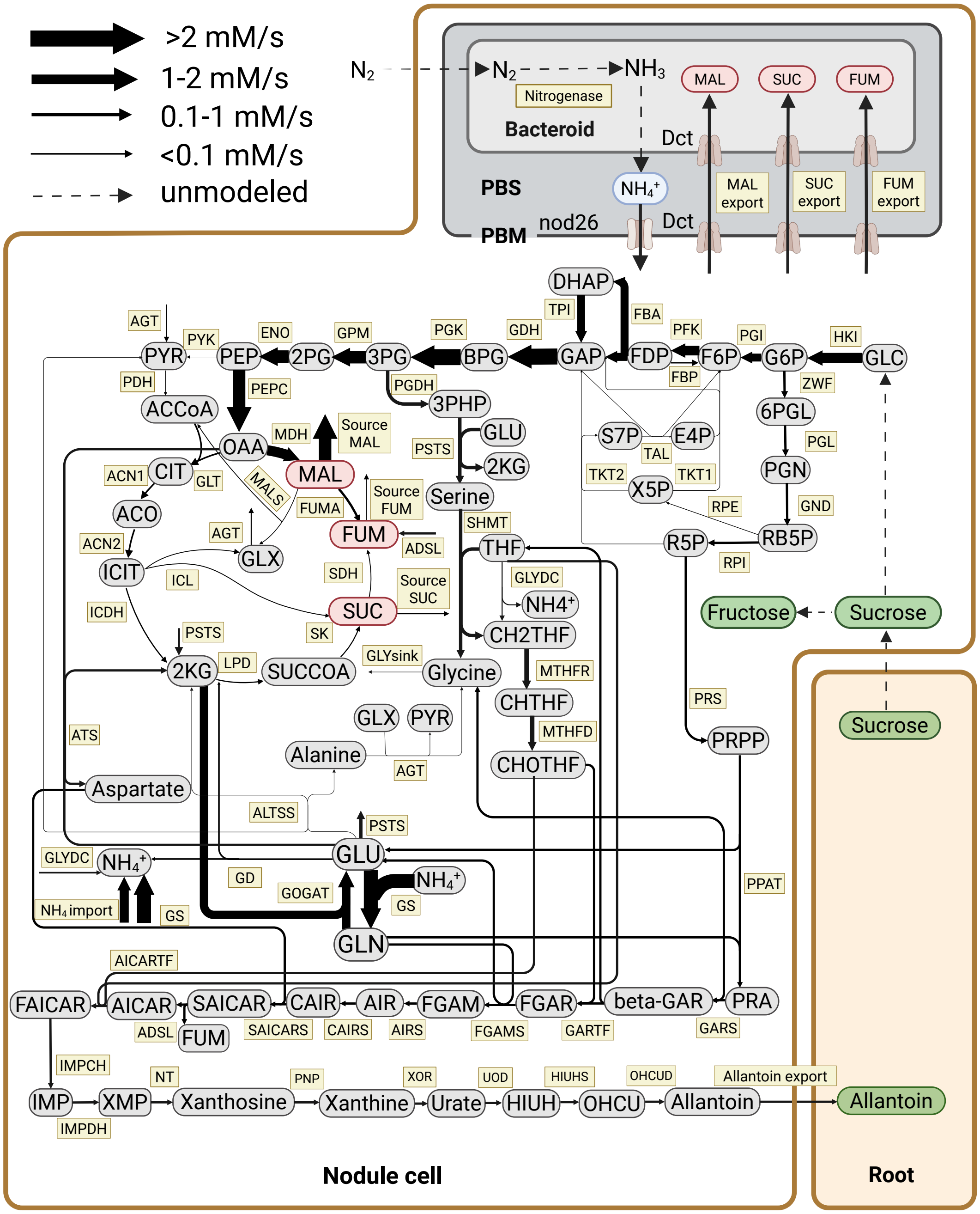

**Figure S11. Average flux map of the subset of flux maps with the best nitrogen fixation efficiency (i.e., lowest gCg$^{-1}$N).** Reaction arrows shown in different colors represent the different magnitudes of fluxes. gCg$^{-1}$N = 3.78.

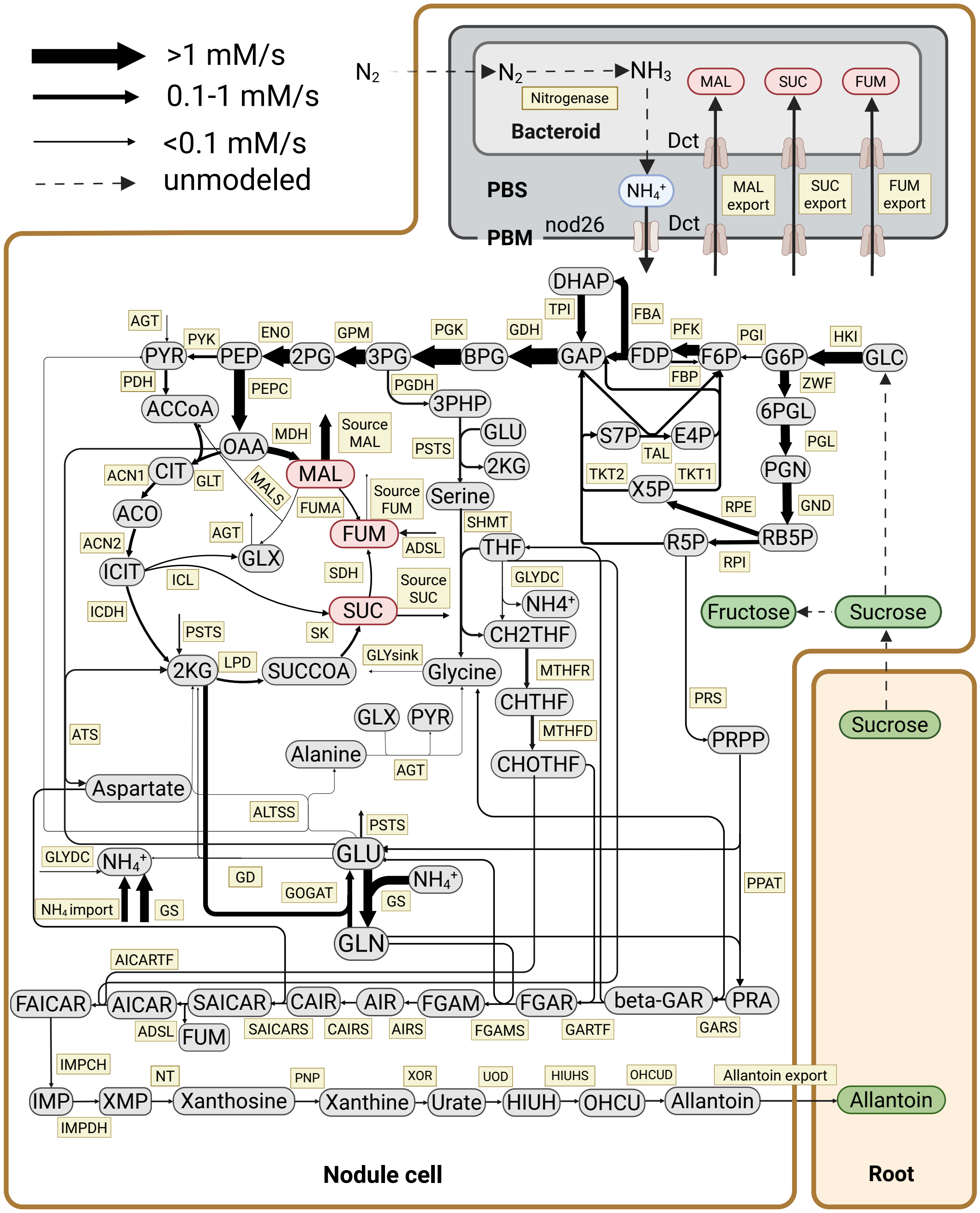

**Figure S12. Average flux map of the subset of flux maps with the worst nitrogen fixation efficiency (i.e., highest $gCg^{-1}N$).** Reaction arrows shown in different colors represent the different magnitudes of fluxes. $gCg^{-1}N = 7.48$.

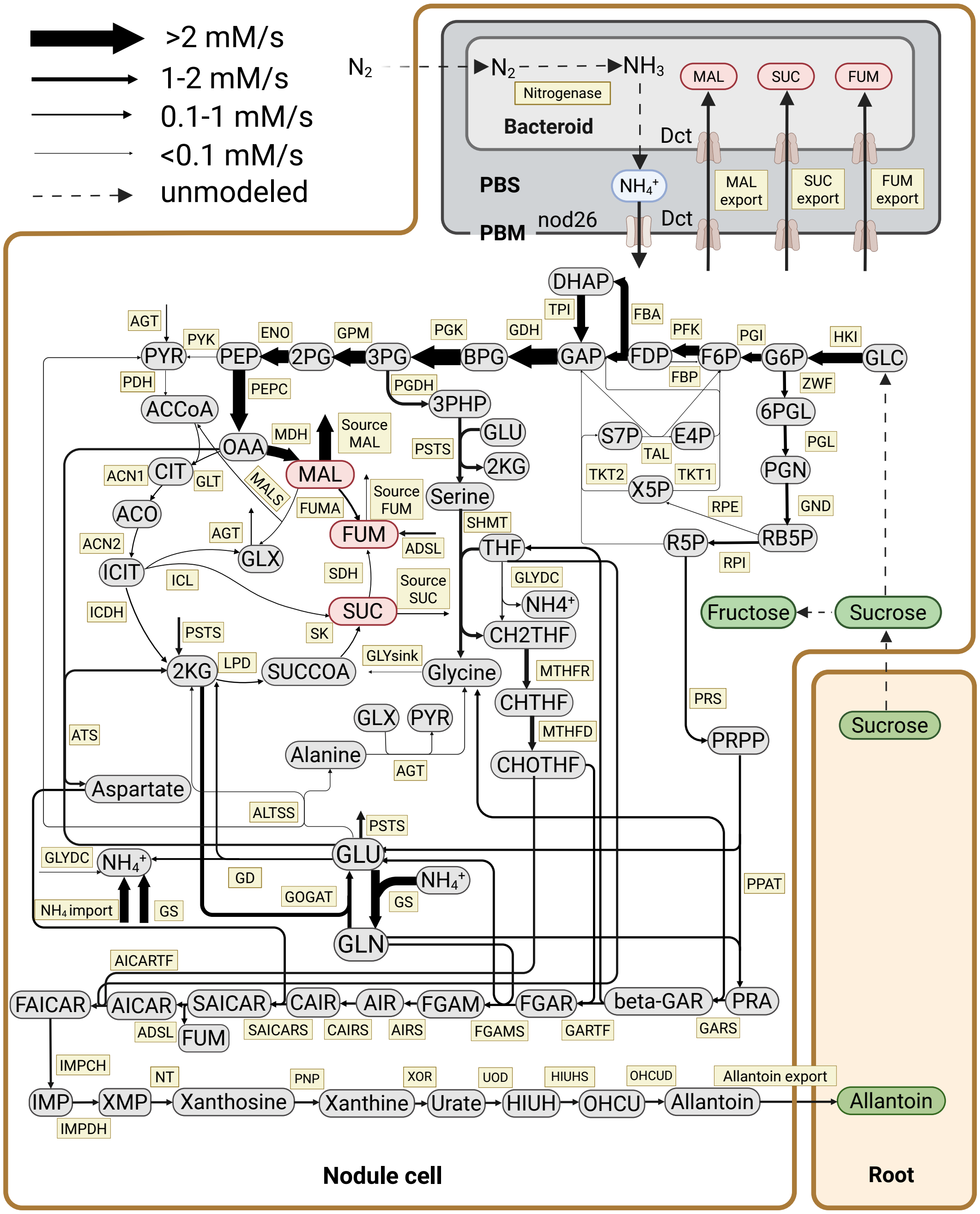

**Figure S13. Average flux map of the subset of flux maps with the highest nitrogen fixation rate (2.76 $mMs^{-1}$).** Reaction arrows shown in different colors represent the different magnitudes of fluxes.

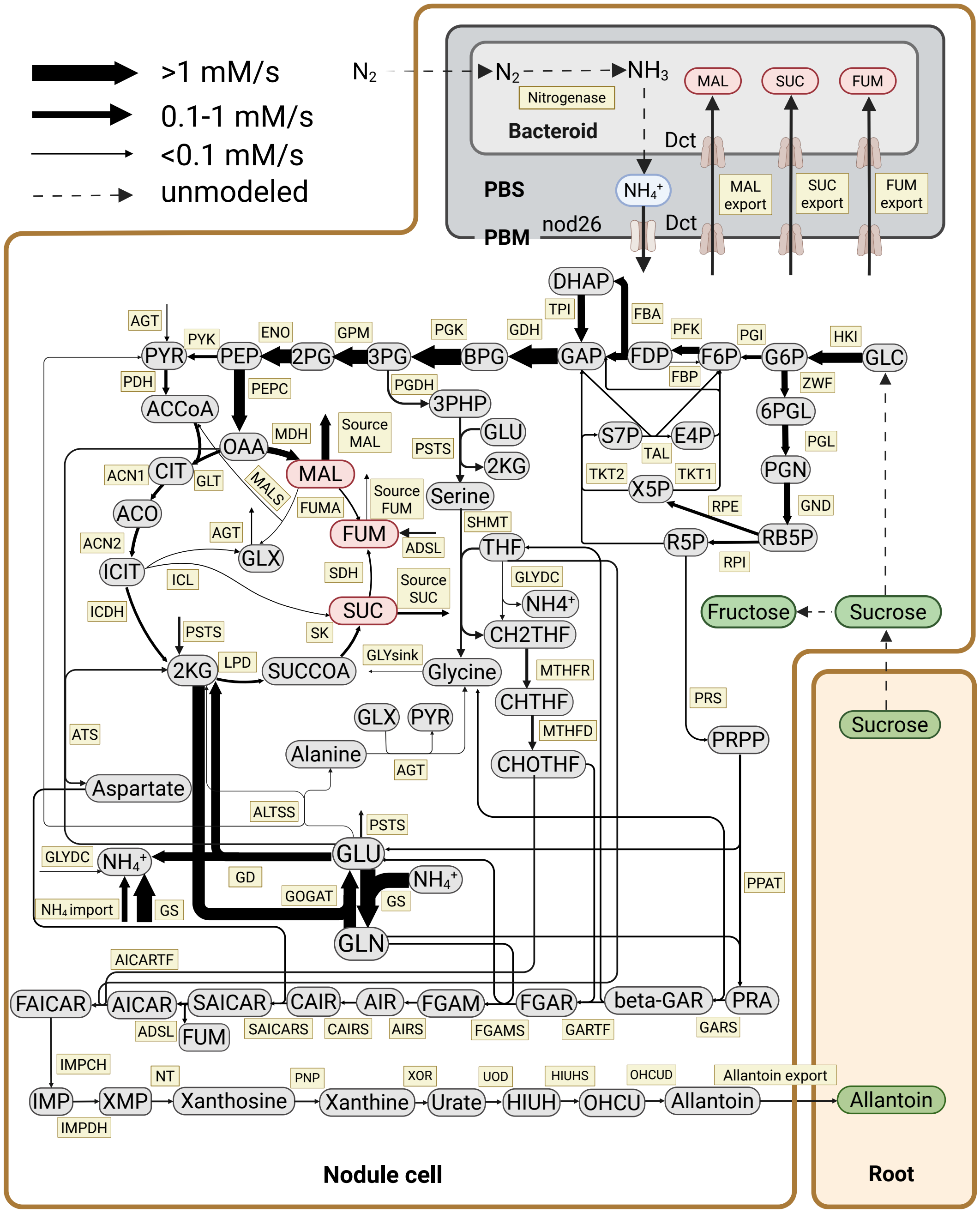

**Figure S14. Average flux map of the subset of flux maps with the lowest nitrogen fixation rate (0.60 mMs$^{-1}$).** Reaction arrows shown in different colors represent the different magnitudes of fluxes.

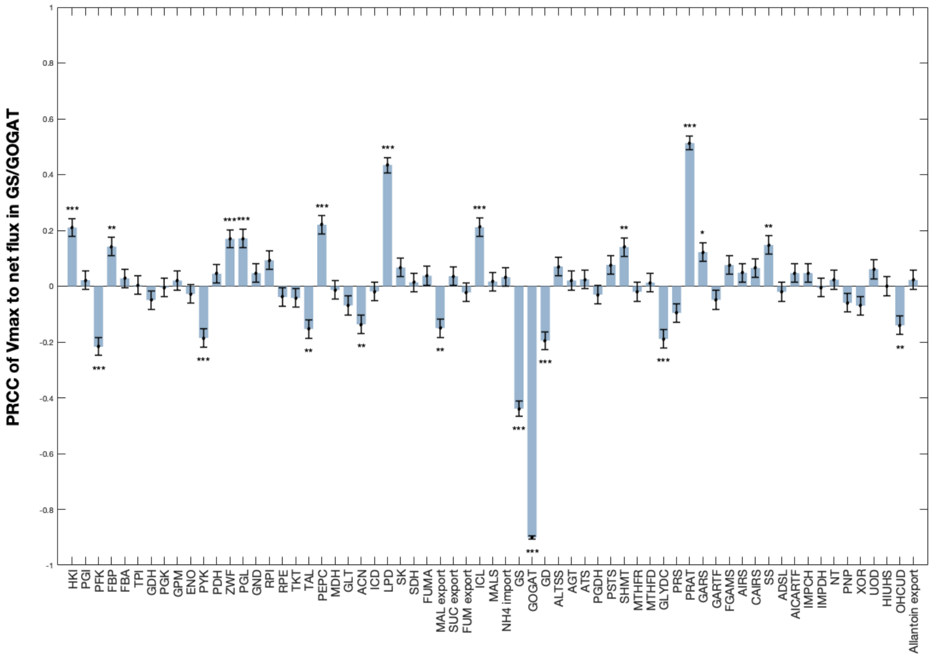

**Figure S15. The impact of enzyme Vmax on net flux in GS/GOGAT.**

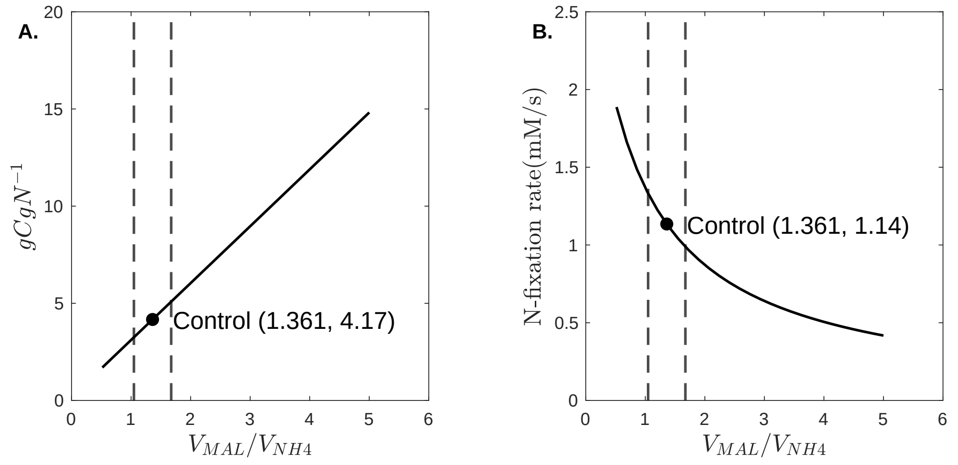

**Figure S16. FBA ratio changes.** The dashed lines represent the range of adjusting bacteroid efficiency ratio by ±23%.

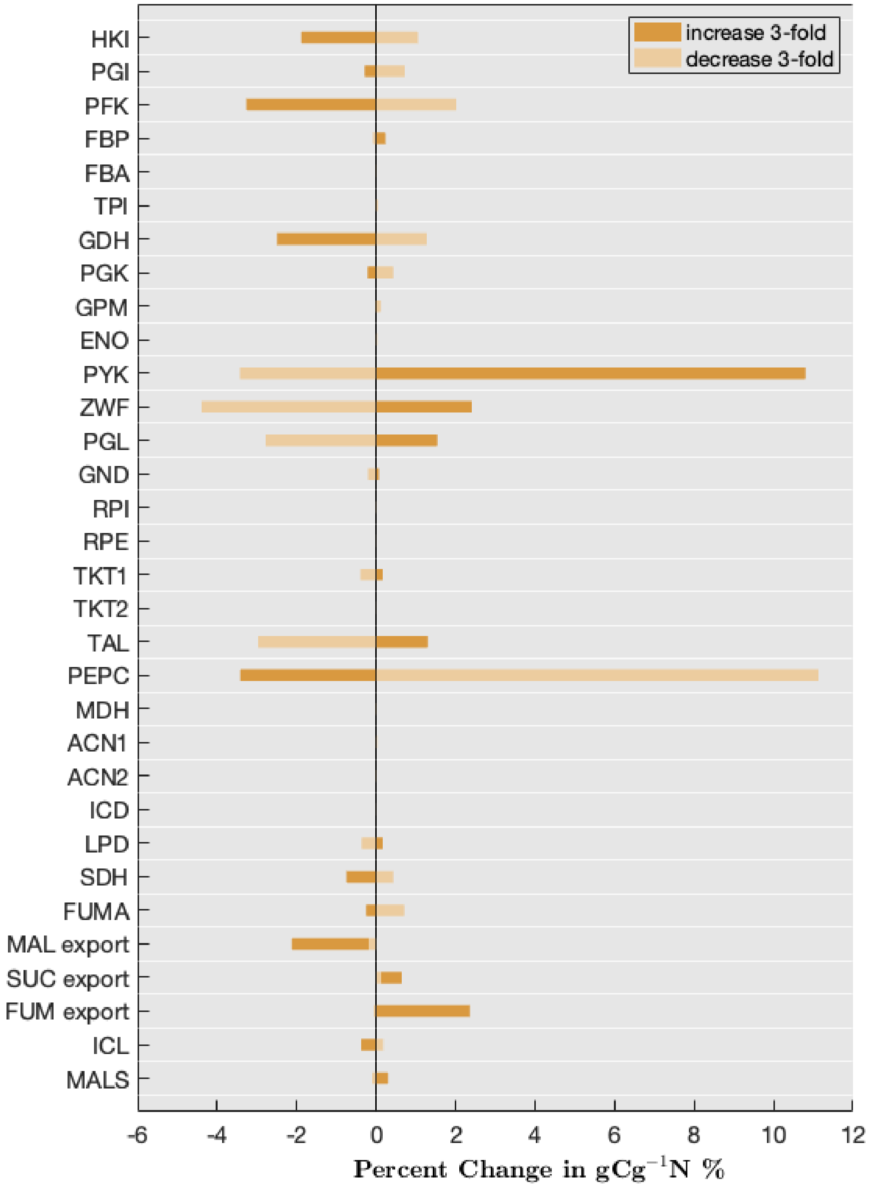

**Figure S17. Percent change in N-fixation efficiency by adjusting Vmax ±3 fold.** Enzymes that have % changes less than 0.1% are removed.

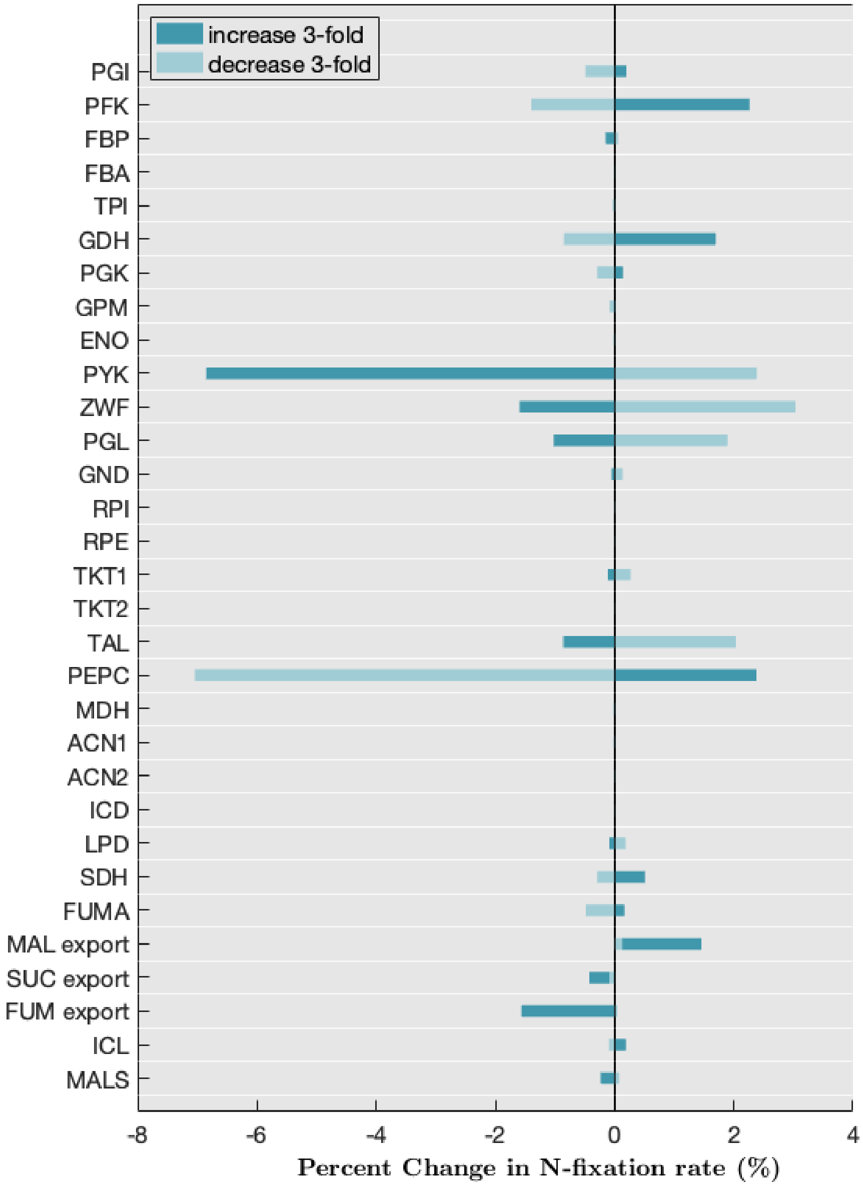

**Figure S18. Percent change in N-fixation rate by adjusting Vmax ±3 fold.** Enzymes that have % changes less than 0.1% are removed.

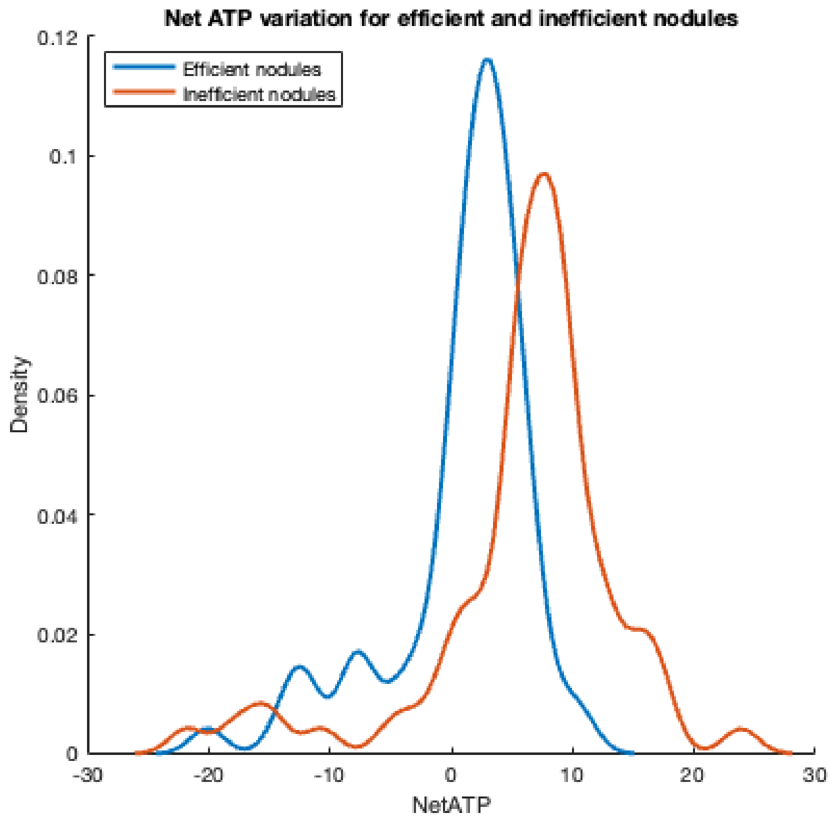

**Figure S19. Net ATP distribution by different levels of N-fixation efficiency (carbon cost).**

**Table S2. The influence of changing initial metabolite concentrations on the metabolite concentrations at steady-state.**

| Metabolite name | Concentration at steady-state (mM) | Metabolite name | Concentration at steady-state (mM) |
|---|---|---|---|
| GLC | $28.89 \pm 1.78E{-}14$ | NH4 | $2.67 \pm 5.47E{-}06$ |
| G6P | $0.067 \pm 2.81E{-}12$ | GLU | $0.99 \pm 2.53E{-}06$ |
| F6P | $0.016 \pm 6.01E{-}13$ | GLN | $0.28 \pm 1.77E{-}06$ |
| FDP | $0.020 \pm 1.25E{-}12$ | PRPP | $0.20 \pm 2.62E{-}10$ |
| GAP | $0.012 \pm 4.11E{-}13$ | PHP3 | $0.17 \pm 5.03E{-}12$ |
| DAP | $0.32 \pm 9.24E{-}12$ | SRN | $1.24 \pm 3.39E{-}06$ |
| BPG | $0.0077 \pm 2.50E{-}13$ | GLY | $2.91 \pm 6.19E{-}06$ |
| PGA3 | $0.022 \pm 6.99E{-}13$ | PRA | $0.11 \pm 7.30E{-}06$ |
| PGA2 | $0.0095 \pm 2.68E{-}13$ | GAR | $0.12 \pm 1.65E{-}05$ |
| PEP | $9.97E{-}04 \pm 2.71E{-}14$ | FGAR | $0.040 \pm 4.97E{-}09$ |
| PYR | $8.42E{-}04 \pm 2.27E{-}14$ | ALN | $0.066 \pm 1.72E{-}07$ |
| GL6P | $0.059 \pm 1.53E{-}11$ | THF | $0.055 \pm 1.92E{-}09$ |
| PGN | $0.60 \pm 4.37E{-}10$ | CH2THF | $4.83E{-}06 \pm 1.27E{-}16$ |
| RB5P | $5.53 \pm 4.20E{-}09$ | CHTHF | $5.43E{-}04 \pm 1.44E{-}14$ |
| R5P | $1.81 \pm 1.37E{-}09$ | CHOTHF | $50.05 \pm 25.03$ |
| X5P | $5.50 \pm 4.19E{-}09$ | FGAM | $0.026 \pm 6.74E{-}13$ |
| E4P | $0.0023 \pm 9.75E{-}13$ | AIR | $0.0066 \pm 1.73E{-}13$ |
| S7P | $0.86 \pm 2.61E{-}09$ | CAIR | $0.0015 \pm 4.04E{-}11$ |
| OAA | $0.092 \pm 2.60E{-}12$ | ASP | $0.42 \pm 1.15E{-}06$ |
| MAL | $0.045 \pm 1.26E{-}12$ | SAICAR | $0.013 \pm 2.28E{-}07$ |
| ACCOA | $2.88E{-}04 \pm 6.49E{-}15$ | AICAR | $0.0042 \pm 2.26E{-}07$ |
| CIT | $0.018 \pm 5.79E{-}13$ | FAICAR | $0.0091 \pm 2.41E{-}13$ |
| ACO | $0.018 \pm 5.33E{-}13$ | IMP | $0.0035 \pm 9.18E{-}14$ |
| ICIT | $0.0021 \pm 5.12E{-}14$ | XMP | $0.0056 \pm 1.46E{-}13$ |
| AKG | $0.031 \pm 7.55E{-}13$ | XAO | $0.0046 \pm 1.22E{-}13$ |
| GLX | $3.56E{-}04 \pm 6.64E{-}10$ | XAN | $5.09E{-}05 \pm 1.34E{-}15$ |
| SUCCOA | $0.0014 \pm 3.52E{-}14$ | URATE | $7.90E{-}05 \pm 2.08E{-}15$ |
| SUC | $0.0025 \pm 7.98E{-}14$ | HIUH | $1.24E{-}04 \pm 3.26E{-}15$ |
| FUM | $0.0014 \pm 4.30E{-}14$ | OHCU | $0.0015 \pm 3.82E{-}14$ |
| NH4_sink | $0.013 \pm 3.67E{-}13$ | ALTN | $6.40E{-}04 \pm 1.68E{-}14$ |

**Table S3. The influence of changing initial metabolite concentrations on the model output at reaction rates at steady-state.**

| Reaction name | Flux at steady-state ($mMs^{-1}$) | Reaction name | Flux at steady-state ($mMs^{-1}$) |
|---|---|---|---|
| HKI | $1.37 \pm 2.01E{-}14$ | FUM_out | $0.01 \pm 2.77E{-}13$ |
| PGI | $0.77 \pm 6.19E{-}11$ | FUMA | $0.29 \pm 7.01E{-}12$ |
| PFK | $1.02 \pm 3.73E{-}11$ | ToNH4 | $1.14 \pm 3.02E{-}11$ |
| FBP | $0.04 \pm 2.37E{-}12$ | GS | $1.94 \pm 6.17E{-}06$ |
| FBA | $0.98 \pm 1.14E{-}10$ | GOGAT | $1.37 \pm 6.17E{-}06$ |
| TPI | $0.98 \pm 3.42E{-}11$ | GD | $0.66 \pm 6.17E{-}06$ |
| GDH | $2.07 \pm 5.44E{-}11$ | PRAT | $0.28 \pm 2.21E{-}10$ |
| PGK | $2.07 \pm 5.49E{-}11$ | PGDH | $0.42 \pm 1.08E{-}11$ |
| GPM | $1.65 \pm 4.37E{-}11$ | PSTS | $0.42 \pm 1.63E{-}11$ |
| ENO | $1.65 \pm 4.42E{-}11$ | SHMT | $0.42 \pm 1.14E{-}11$ |
| PYK | $0.05 \pm 1.38E{-}12$ | GARS | $0.28 \pm 1.27E{-}11$ |
| PDH | $0.05 \pm 1.38E{-}12$ | GARTF | $0.28 \pm 7.38E{-}12$ |
| ZWF | $0.60 \pm 8.45E{-}11$ | FGAMS | $0.28 \pm 7.33E{-}12$ |
| PGL | $0.60 \pm 9.35E{-}11$ | ALTSS | $0.01 \pm 1.18E{-}12$ |
| GND | $0.60 \pm 2.04E{-}10$ | AGT | $0.01 \pm 8.90E{-}13$ |
| RPE | $-0.21 \pm 9.74E{-}11$ | MTHFR | $0.57 \pm 1.47E{-}11$ |
| RPI | $0.39 \pm 8.49E{-}11$ | MTHFD | $0.57 \pm 1.47E{-}11$ |
| TKT1 | $-0.10 \pm 6.72E{-}11$ | GLYDC | $0.15 \pm 3.66E{-}12$ |
| TKT2 | $-0.10 \pm 6.82E{-}08$ | AIRS | $0.28 \pm 7.37E{-}12$ |
| TAL | $-0.10 \pm 1.13E{-}10$ | CAIRS | $0.28 \pm 7.37E{-}12$ |
| PRS | $0.28 \pm 1.03E{-}08$ | SS | $0.28 \pm 7.43E{-}12$ |
| PEPC | $1.59 \pm 4.29E{-}11$ | ATS | $0.28 \pm 9.93E{-}11$ |
| MDH | $1.26 \pm 3.22E{-}11$ | ADSL | $0.28 \pm 7.29E{-}12$ |
| MAL_out | $1.55 \pm 4.11E{-}11$ | AICARTF | $0.28 \pm 7.37E{-}12$ |
| GLT | $0.05 \pm 1.41E{-}12$ | IMPCH | $0.28 \pm 7.37E{-}12$ |
| ACN1 | $0.05 \pm 1.39E{-}12$ | IMPDH | $0.28 \pm 7.37E{-}12$ |
| ACN2 | $0.05 \pm 1.40E{-}12$ | Glycine_sink | $8.44E{-}04 \pm 2.24E{-}14$ |
| ICL | $0.01 \pm 1.76E{-}13$ | NT | $0.28 \pm 7.37E{-}12$ |
| MALS | $-6.71E{-}04 \pm 2.95E{-}13$ | PNP | $0.28 \pm 7.37E{-}12$ |
| ICDH | $0.04 \pm 1.22E{-}12$ | XOR | $0.28 \pm 7.37E{-}12$ |
| LPD | $0.04 \pm 1.10E{-}12$ | UOD | $0.28 \pm 7.37E{-}12$ |
| SK | $0.04 \pm 1.10E{-}12$ | HIUHS | $0.28 \pm 7.37E{-}12$ |
| SUC_out | $0.04 \pm 1.25E{-}12$ | OHCUD | $0.28 \pm 7.37E{-}12$ |
| SDH | $0.01 \pm 7.66E{-}14$ | Allantoin_out | $0.28 \pm 7.37E{-}12$ |

**Table S4. The influence of changing initial CHOTHF concentrations on the flux at steady-state.**

| Reaction name | Flux at steady-state ($mMs^{-1}$) | Reaction name | Flux at steady-state ($mMs^{-1}$) |
|---|---|---|---|
| HKI | $1.37 \pm 1.73E{-14}$ | FUM_out | $0.0088 \pm 2.32E{-13}$ |
| PGI | $0.77 \pm 5.34E{-11}$ | FUMA | $0.29 \pm 5.85E{-12}$ |
| PFK | $1.02 \pm 3.15E{-11}$ | ToNH4 | $1.14 \pm 2.54E{-11}$ |
| FBP | $0.041 \pm 1.99E{-12}$ | GS | $1.94 \pm 3.59E{-07}$ |
| FBA | $0.98 \pm 9.81E{-11}$ | GOGAT | $1.37 \pm 3.59E{-07}$ |
| TPI | $0.98 \pm 2.89E{-11}$ | GD | $0.66 \pm 3.59E{-07}$ |
| GDH | $2.07 \pm 4.58E{-11}$ | PRAT | $0.28 \pm 1.78E{-10}$ |
| PGK | $2.07 \pm 4.62E{-11}$ | PGDH | $0.42 \pm 9.09E{-12}$ |
| GPM | $1.65 \pm 3.68E{-11}$ | PSTS | $0.42 \pm 1.22E{-11}$ |
| ENO | $1.65 \pm 3.72E{-11}$ | SHMT | $0.42 \pm 9.84E{-12}$ |
| PYK | $0.051 \pm 1.17E{-12}$ | GARS | $0.28 \pm 9.60E{-12}$ |
| PDH | $0.051 \pm 1.16E{-12}$ | GARTF | $0.28 \pm 6.18E{-12}$ |
| ZWF | $0.60 \pm 7.29E{-11}$ | FGAMS | $0.28 \pm 6.12E{-12}$ |
| PGL | $0.60 \pm 8.06E{-11}$ | ALTSS | $0.0078 \pm 4.68E{-13}$ |
| GND | $0.60 \pm 1.76E{-10}$ | AGT | $0.0078 \pm 2.26E{-13}$ |
| RPE | $-0.21 \pm 8.81E{-11}$ | MTHFR | $0.57 \pm 1.23E{-11}$ |
| RPI | $0.39 \pm 7.73E{-11}$ | MTHFD | $0.57 \pm 1.23E{-11}$ |
| TKT1 | $-0.10 \pm 5.85E{-11}$ | GLYDC | $0.15 \pm 3.04E{-12}$ |
| TKT2 | $-0.10 \pm 5.91E{-08}$ | AIRS | $0.28 \pm 6.17E{-12}$ |
| TAL | $-0.10 \pm 9.77E{-11}$ | CAIRS | $0.28 \pm 6.17E{-12}$ |
| PRS | $0.28 \pm 1.03E{-08}$ | SS | $0.28 \pm 6.22E{-12}$ |
| PEPC | $1.59 \pm 3.61E{-11}$ | ATS | $0.28 \pm 8.93E{-12}$ |
| MDH | $1.26 \pm 2.71E{-11}$ | ADSL | $0.28 \pm 6.07E{-12}$ |
| MAL_out | $1.55 \pm 3.46E{-11}$ | AICARTF | $0.28 \pm 6.17E{-12}$ |
| GLT | $0.052 \pm 1.17E{-12}$ | IMPCH | $0.28 \pm 6.17E{-12}$ |
| ACN1 | $0.052 \pm 1.16E{-12}$ | IMPDH | $0.28 \pm 6.17E{-12}$ |
| ACN2 | $0.052 \pm 1.16E{-12}$ | Glycine_sink | $8.44E{-04} \pm 1.89E{-14}$ |
| ICL | $0.0071 \pm 1.44E{-13}$ | NT | $0.28 \pm 6.17E{-12}$ |
| MALS | $-6.71E{-04} \pm 2.27E{-14}$ | PNP | $0.28 \pm 6.17E{-12}$ |
| ICDH | $0.045 \pm 1.02E{-12}$ | XOR | $0.28 \pm 6.17E{-12}$ |
| LPD | $0.045 \pm 8.98E{-13}$ | UOD | $0.28 \pm 6.17E{-12}$ |
| SK | $0.045 \pm 8.98E{-13}$ | HIUHS | $0.28 \pm 6.17E{-12}$ |
| SUC_out | $0.040 \pm 1.04E{-12}$ | OHCUD | $0.28 \pm 6.17E{-12}$ |
| SDH | $0.012 \pm 1.80E{-14}$ | Allantoin_out | $0.28 \pm 6.17E{-12}$ |

**Table S5. Uninfluential enzyme pairs on N-fixation efficiency and rate.**

| | | | | |
|---|---|---|---|---|
| FBP-, FGAMS | FBP-, FUMout | FBP-, SUCout | FBP-, IMPCH | FBP-, IMPDH |
| FBP-, ALTSS | FBP-, GD | FBP-, PRAT | FBP-, SHMT | FGAMS, IMPCH |
| FGAMS, IMPDH | FUMA+, ALTSS | FUMA+, FGAMS | FUMA+, FUMout | FUMA+, GD |
| FUMA+, IMPCH | FUMA+, IMPDH | FUMA+, PRAT | FUMA+, SHMT | FUMA+, SUCout |
| FUMA+, SUCout | ICL+, ALTSS | ICL+, FGAMS | ICL+, GD | ICL+, IMPCH |
| ICL+, IMPDH | ICL+, PRAT | ICL+, SHMT | FUMout, ALTSS | FUMout, FGAMS |
| FUMout, GD | FUMout, ICL | FUMout, IMPCH | FUMout, IMPDH | FUMout, PRAT |
| FUMout, SHMT | IMPCH, IMPDH | LPD-, ALTSS | LPD-, FGAMS | LPD-, FUMout |
| LPD-, GD | LPD-, IMPCH | LPD-, IMPDH | LPD-, PRAT | LPD-, SHMT |
| LPD-, SUCout | PGL-, ALTSS | PGL-, FGAMS | PGL-, FUCout | PGL-, GD |
| PGL-, IMPCH | PGL-, IMPDH | PGL-, PRAT | PGL-, SHMT | PGL-, SUCout |
| PGI+, ALTSS | PGI+, FGAMS | PGI+, FUMout | PGI+, GD | PGI+, IMPCH |
| PGI+, IMPDH | PGI+, PRAT | PGI+, SHMT | PGI+, SUCout | ZWF-, ALTSS |
| ZWF-, FGAMS | ZWF-, FUMout | ZWF-, GD | ZWF-, IMPCH | ZWF-, IMPDH |
| ZWF-, PRAT | ZWF-, SHMT | ZWF-, SUCout | MALout+, ALTSS | MALout+, FGAMS |
| MALout+, GD | MALout+, FUMout | MALout+, IMPCH | MALout+, IMPDH | MALout+, PRAT |
| MALout+, SHMT | MALout+, SUCout | PEPC+, ALTSS | PEPC+, FGAMS | PEPC+, FUMout |
| PEPC+, GD | PEPC+, IMPCH | PEPC+, IMPDH | PEPC+, PRAT | PEPC+, SHMT |
| PEPC+, SUCout | PFK+, ALTSS | PFK+, FGAMS | PFK+, FUMout | PFK+, GD |
| PFK+, IMPCH | PFK+, IMPDH | PFK+, PRAT | PFK+, SHMT | PFK+, SUCout |
| PRAT, FGAMS | PRAT, IMPCH | PRAT, IMPDH | SDH+, ALTSS | SDH+, FGAMS |
| SDH+, FUMout | SDH+, GD | SDH+, IMPCH | SDH+, IMPDH | SDH+, SUCout |
| SDH+, PRAT | SDH+, SHMT | ALTSS, FGAMS | ALTSS, IMPCH | ALTSS, IMPDH |
| ALTSS, PRAT | ALTSS, SHMT | GD, ALTSS | GD, FGAMS | GD, IMPCH |
| GD, IMPDH | GD, PRAT | GD, SHMT | SHMT, FGAMS | SHMT, IMPCH |
| SHMT, IMPDH | SHMT, PRAT | SUCout, IMPCH | SUCout, IMPDH | SUCout, FUMout |
| SUCout, GD | SUCout, ICL+ | SUCout, IMPCH | SUCout, IMPDH | SUCout, PRAT |
| SUCout, SHMT | TAL-, ALTSS | TAL-, FGAMS | TAL-, FUMout | TAL-, GD |
| TAL-, IMPCH | TAL-, IMPDH | TAL-, PRAT | TAL-, SHMT | TAL-, SUCout |
| PYK-, ALTSS | PYK-, FGAMS | PYK-, FUMout | PYK-, GD | PYK-, IMPCH |
| PYK-, IMPDH | PYK-, PRAT | PYK-, SHMT | PYK-, SUCout | |
